# Massive inter-animal horizontal gene transfer contributes to stress tolerance in the fungus gnat *Bradysia coprophila*

**DOI:** 10.64898/2026.11.06.764122

**Authors:** Rishabh R. Kapoor, Tarun Kumar, Piyush Nanda, Supanat Phuangphong, Andrew W. Murray, Cassandra G. Extavour

## Abstract

Horizontal gene transfer (HGT) in animals is thought to be rare and largely restricted to transfers from non-animal sources^1–3^. In contrast to this expectation, we discovered that 9.2% of the protein-coding genes in the genome of the dark-winged fungus gnat *Bradysia coprophila* (order: Diptera, family: Sciaridae) are of potential HGT origin. Most of these genes have phylogenetic affinity with sequences from other distantly related arthropods that inhabit a shared soil-dwelling niche with *B. coprophila*. We show that these findings are inconsistent with genomic contamination or widespread gene loss and instead suggest a level of inter-animal HGT that is, to our knowledge, unprecedented. Consistent with their potential functional importance, ∼75% of putative *B. coprophila* horizontally transferred genes (HTGs) are transcribed, and nearly all are shared across multiple sciarid fungus gnat species that diverged tens of millions of years ago. Further, domain annotations suggest that HTGs participate in diverse processes potentially important to *B. coprophila* ecology. We tested the functions of eight putative *B. coprophila* HTGs through transgenic expression in the fly *Drosophila melanogaster* and the yeast *Saccharomyces cerevisiae*, and found evidence that HGT has likely endowed *B. coprophila* with the novel capability to biosynthesize NAD^+^, as well as enhanced capacity to withstand stressors imposed by arsenic, cyanide, alkylating agents, and reactive oxygen species. Our findings reveal a previously underappreciated contribution of inter-animal genetic exchange to the evolution of genomic, physiological, and ecological novelty.

## Introduction

Horizontal gene transfer (HGT), the non-Mendelian movement of DNA between organisms, has varying prevalence across the tree of life. On average, ∼10% of protein-coding genes in bacterial genomes have been acquired from other bacteria via recent HGT^4,5^. HGT from bacteria to animals has been increasingly reported and has likely contributed to adaptation in multiple lineages^1,6,7^. In contrast to HGT from non-animal sources, HGT between metazoans (animals) has rarely been reported and appears largely restricted to cases involving transposons^8–10^, host-parasite interactions^11–13^ or marine fish^3,14^. Most existing studies of HGT have been designed to detect transfers from non-metazoan genomes^2,15–20^. Thus, the apparent rarity of inter-animal HGT might in part reflect methodological biases.

We and others studying specific gene families have recently uncovered signatures of HGT in dark-winged fungus gnats (order: Diptera, family: Sciaridae)^21–31^, including putative inter-metazoan transfers (Table S1)^29,31–33^. Our recent screen of 319 arthropod genomes for genes formed by in-frame fusion of horizontally transferred sequences with pre-existing coding genes revealed that the fungus gnat *Bradysia coprophila* contained more than ten times the average number of these chimeras across all surveyed arthropods^32^. This suggested that the genome of this gnat might be particularly prone to horizontal gene acquisition. To our knowledge, however, no systematic analysis of HGT in sciarids has been conducted to date.

To that end, we used sequence similarity searches to detect HGT across dipteran (fly and mosquito) genomes, including *B. coprophila*. Subsequent phylogenetic analyses revealed that 9.2% of *B. coprophila* protein-coding genes are likely of HGT origin, and that most of those genes were likely obtained from other soil-dwelling animals rather than from microbes. We present multiple lines of evidence that *B. coprophila* horizontally transferred genes (HTGs) are functional, including expression data, evolutionary conservation, and domain annotations. Finally, with *in vivo* experimental analysis of eight HTGs via transgenic expression assays in *Drosophila* and yeast, we show that these HTGs can change the physiological capabilities of their hosts. Collectively, this study illuminates an underappreciated contribution of inter-animal HGT to innovation in animal evolution.

## Results

### Massive HGT shaped the B. coprophila genome

We searched fly and mosquito (order Diptera) genomes for signs of HGT from any non-dipteran donor. Because standard HGT detection methodologies can only detect HGT from non-animal sources^2,16,17,19^, we modified the Alien Index^16^ metric to search for genes with non-dipteran affinity using DIAMOND BLASTp^34^ (Figure 1A). We applied this methodology to protein sequences from every protein-coding gene in 54 dipteran genomes. Whereas the mean number and percentage of putative HTGs in non-sciarid dipteran genomes are 260.8 and 1.7%, respectively, the *B. coprophila* genome contains 1,721 putative HTGs constituting 10.5% of its protein-coding genes. We observed a greater number and percentage of putative HTGs for *B. coprophila* than for any other dipteran genome examined in this or previous studies^2,15^ (Figure 1B; Table S2).

**Figure 1.**
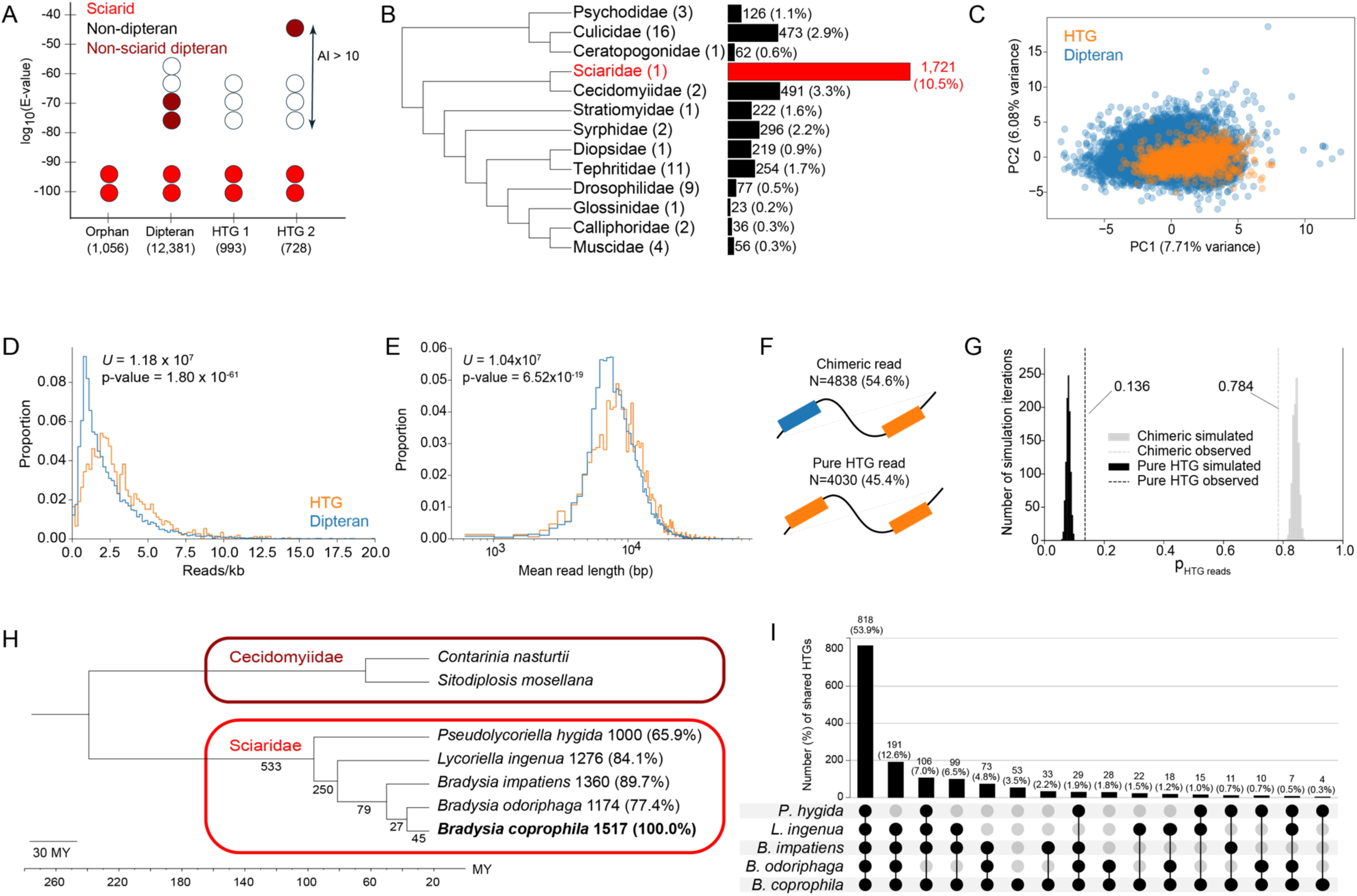
Signatures of massive HGT in the *B. coprophila* genome. **(A)** Illustration of our methodology for inference of gene ancestry from the taxonomic distribution of DIAMOND hits, with numbers beneath each ancestry annotation (see Methods) indicating the number of corresponding *B. coprophila* proteins. Circles represent DIAMOND hits colored by taxonomic source. HTG 1 and HTG 2 represent patterns interpreted as transfers from non-dipteran sources. Dipteran-affinity genes are assumed to be vertically inherited. **(B)** Phylogram^126^ of 13 dipteran families subjected to DIAMOND-based HTG inference, with the number of searched RefSeq genomes per family indicated in parentheses. Bar graph and adjacent numbers show the number and percentage of protein-coding HTGs per genome averaged across all genomes analyzed per family, rounded to the nearest integer and tenth of a percentage point, respectively. **(C)** Principal component analysis of coding region codon usage. Each dot represents codon usage of one gene, colored by gene ancestry. **(D)** Pooled Oxford Nanopore (ONT) read depth overlapping *B. coprophila* genes per kilobase, separated by gene ancestry (μ_HTG_= 3.09; σ_HTG_=2.14; μ_Dipt_= 2.38; σ_Dipt_=2.12). **(E)** Mean ONT read length overlapping each gene (log scale), separated by ancestry type (μ_HTG_= 1.05×10^4^; σ_HTG_=5.00×10^3^; μ_Dipt_= 9.54×10^3^; σ_Dipt_=4.37×10^3^). Asymptotic two-sided Mann-Whitney test statistics are indicated for (D)-(E). **(F)** “Chimeric reads” = ONT reads containing at least one putative HTG (orange rectangle) and one dipteran gene (blue rectangle). “Pure HTG reads” = ONT reads containing only putative HTGs. Percentages of the total number of putative HTG-containing reads in each category are indicated. **(G)** Fraction of HTGs found exclusively in ONT reads containing only HTGs or in at least one read containing a dipteran gene (dashed lines). Histograms show simulated null distributions. **(H)** Fossil-calibrated sciarid species tree (Figure S8A). Numbers at branches indicate the number of transfer events (of 934) inferred to have occurred in the corresponding interval. Number and % of *B. coprophila* HTGs shared with other sciarid species are indicated to the right of species names. **(I)** UpSet plot^127^ of the presence of homologs of HTGs from *B. coprophila* in other sciarid genomes.

To assess phylogenetic support for these putative HTGs, we first clustered all 1,721 HTGs by sequence similarity, then built maximum-likelihood gene trees for the resulting 924 clusters. We then inspected each clade of sciarid sequences for evidence of phylogenetic grouping with non-dipteran sequences with > 70% bootstrap support at the relevant node. This manual inspection (Table S3) provided support for 934 putative transfer events from non-dipterans to sciarids. When accounting for post-transfer gene family expansions (see SI Note 1; Table S4), we found a total of 1,517 HTGs, representing 9.2% of all protein-coding genes in the *B. coprophila* genome. This final set of putative HTGs has a mean length coverage of 87.7% and mean identity of 49.0% with top non-dipteran hits (Figure S1D). These findings are consistent with the hypothesis that these putative HTGs are closely related to non-dipteran sequences. Further, this set included 79.2% (42/53) of *B. coprophila* genes that were previously identified as HTGs based on phylogenetic evidence (Table S1).

### HGT signatures in B. coprophila are unlikely to result from genome contamination

The high level of HGT inferred in the *B. coprophila* genome could in principle reflect contamination of the *B. coprophila* genome assembly with reads from other taxa^1,35^. However, several lines of evidence suggest the *B. coprophila* reference genome we analyzed (GCF_014529535.2)^36,37^ is not extensively contaminated. First, gene annotations for 95.9% (1,455/1,517) of HTGs could be transferred with an intact open reading frame to an independently sequenced *B. coprophila* genome (GCA_965233685.1)^38^. Genes from contaminants are expected to be sequenced at lower depth^35^ and in shorter reads than endogenous genes^39^, but we detected no decrease in read depth (Figure 1D) or overlapping read length (Figure 1E) for HTGs relative to dipteran-affinity genes, which we assume are vertically inherited. Further, 54.6% of Oxford Nanopore long reads^36^ spanning an HTG also contain a dipteran gene (Figure 1F), and 78.4% of HTGs are found in at least one read containing a dipteran gene (Figure 1G), supporting the inference that HTGs are integrated into the *B. coprophila* genome. In contrast to the prediction that contaminants resemble the sequence composition of their taxon of origin^35^, the codon usage profile of HTGs overlaps that of dipteran genes (Figure 1C). Further, HTG codon usage and GC content distributions more closely resemble those of dipteran-ancestry genes than those of genes from the top putative donor lineage (Figure S1A-C).

We next used our gene trees to search for homologs of *B. coprophila* HTGs in the genomes of four other sciarid species (Table S5)^38,40,41^. We found that 95.2% (889/934) of inferred transfer events gave rise to homologous HTGs shared across at least two independently sequenced sciarid genomes (Figure 1H), and 96.5% (1,464/1,517) of *B. coprophila* HTGs have a homolog in at least one other sciarid genome (Figure 1I). In addition, 83.3% (701/842) of *B. coprophila* HTGs with a detectable reciprocal best BLASTp hit in *Bradysia odoriphaga* are syntenic with at least one adjacent gene. HTG synteny decreases with increasing genomic distance and species divergence from *B. coprophila*, as is also observed for dipteran genes (Figure S1E-F). In sum, we find no evidence that the widespread signature of HGT we detect in the *B. coprophila* genome is an artifact of contamination of its genome assembly.

### Most B. coprophila HTGs are of putative metazoan origin

To determine the potential origins of our putative HTGs, we examined both the most similar non-sciarid sequence to each HTG via DIAMOND (Table S6, Figure 2A), and the closest non-sciarid phylogenetic relative(s) of each HTG in its respective gene tree (Table S3, Figure S2A). Contrary to the preponderance of bacteria-to-insect HGT in prior studies^1,2,42^, both analyses suggest that at least 78.3% (1,188/1,517) of *B. coprophila* HTGs have affinity with sequences from other animals. Specifically, we found that at least 70.9% (1,075/1,517) of HTGs are of non-dipteran arthropod affinity (Figures 2A-B, S2A). The class Collembola (springtails), which are hexapods that diverged from insects over 450 million years ago^43^, accounts for at least 51.7% (785/1,517) of these HTGs.

**Figure 2.**
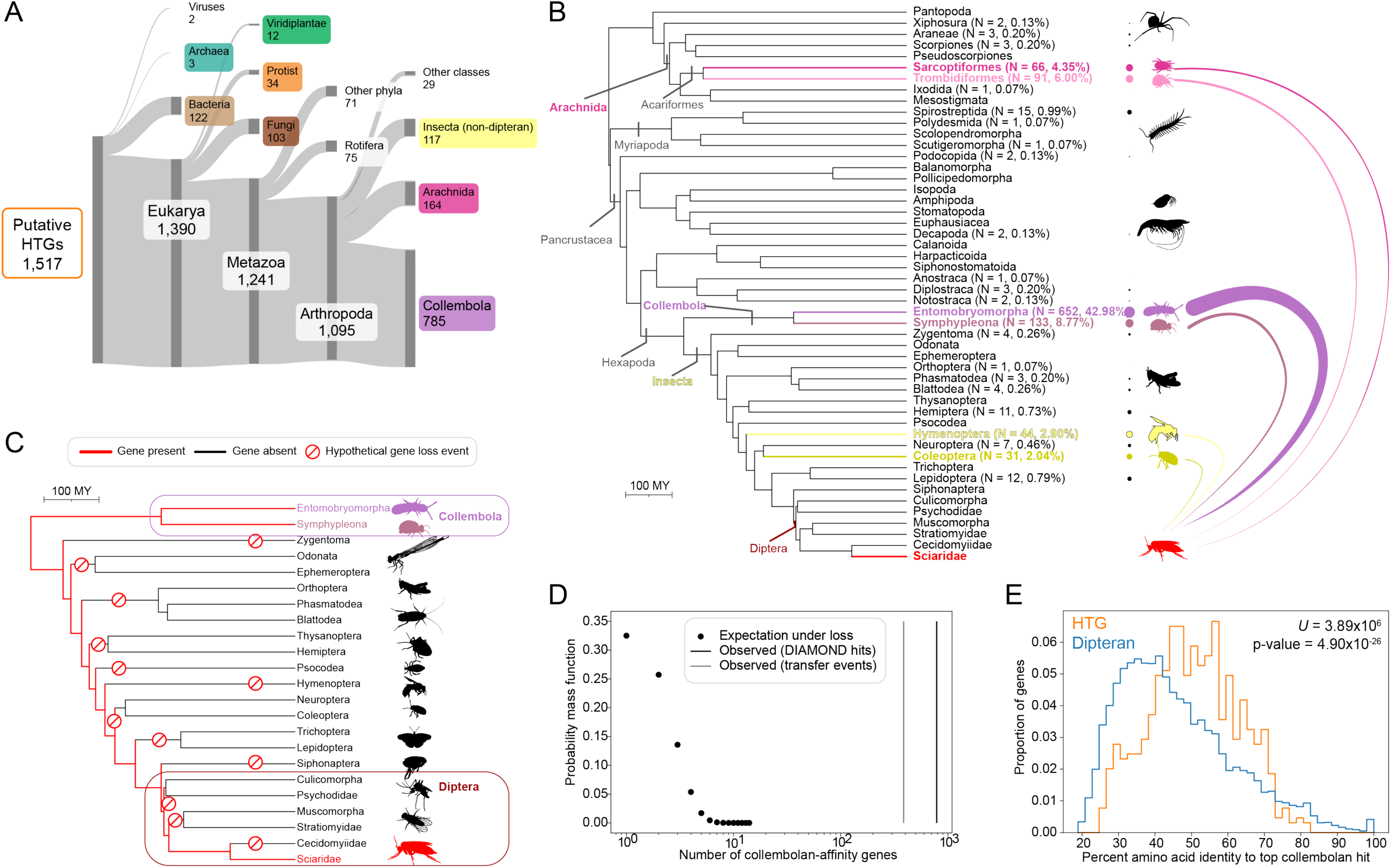
DIAMOND and phylogenetic inference reveal diverse origins of putative *B. coprophila* HTGs. **(A)** Sankey diagram of the taxonomic origin of the top non-sciarid DIAMOND hit for HTGs^116^. Branch height is proportional to the number of HTGs. **(B)** Phylogenetic relationships among arthropod lineages with annotated genomes represented in GenBank or RefSeq at the time of analysis. Tree tips correspond to orders, except for intra-order relationships within Diptera. The number and percentage of total HTGs with a top DIAMOND hit in each order are indicated to the right of each text label in parentheses; absence of numbers adjacent to a clade indicates no evidence of contributions to HGT in the *B. coprophila genome*. Circle sizes are proportional to the logarithm of HTG number. The top six putative donor orders for *B. coprophila* HTGs (family Sciaridae) by top DIAMOND hit are highlighted in colored text/silhouettes (from www.phylopic.org). **(C)** Phylogenetic tree of Hexapoda showing the minimum number of independent losses (red symbols) required to explain the presence of a gene in Collembola and Sciaridae but no other examined hexapod genome without HGT. Terminal leaf nodes are at the order taxonomic level except for Diptera, which is expanded to the infraorder and family level. Red lines indicate conservation of the gene under this hypothesis required since the hexapod common ancestor. **(D)** The expected number of collembolan-affinity genes (μ = 1.58, σ = 1.25) was computed under a null model assuming vertical descent with gene loss. Vertical lines indicate the observed number of putative collembolan-affinity HTGs via DIAMOND (black, N = 785) or collembolan origin transfer events via manual tree inspection (gray, N = 386). **(E)** Comparison of the percent identity to the top collembolan hit for *B. coprophila* proteins of dipteran ancestry and *B. coprophila* HTGs of putative collembolan origin via tree inspection, with one-sided Mann-Whitney test statistics indicated.

The preponderance of HTGs with collembolan affinity raised the alternative hypothesis that these putative HTGs are instead the products of vertical inheritance with gene loss^44^. To evaluate this possibility, we considered genes present in Collembola and Sciaridae but in no other available annotated insect genome, a scenario we have thus far interpreted as evidence of HGT. A consensus phylogeny of hexapods^43^ suggests a minimum of 12 independent losses would be required to explain this sparse gene distribution assuming no HGT (Figure 2C). We adopted a previously derived probabilistic model that assumes strict vertical inheritance and a constant rate of gene loss^45^, and found that the observed number of collembolan-affinity *B. coprophila* genes is more than two orders of magnitude greater than the expectation without HGT (Figure 2D).

We further reasoned that, if these putative HTGs were instead vertically inherited genes, their percent identity to top collembolan hits should be similar to the same statistic computed for dipteran-ancestry genes, which are of vertical descent^44^. Instead, HTGs are significantly more similar to their top collembolan hits than are dipteran sequences (median percent identity 50.8 vs. 40.3; one-sided Mann-Whitney *U* statistic = 3.89×10^6^; p-value = 4.90×10^-26^; Figure 2E). Under the simplifying assumption of a uniform substitution rate across genes, this result suggests that less time has elapsed since the common ancestor of *B. coprophila* HTGs and their collembolan homologs than since the common ancestor of *B. coprophila* and Collembola. This scenario is inconsistent with a hypothesis of strict vertical inheritance. Collectively, the data are most consistent with the hypothesis that massive inter-metazoan HGT has shaped the *B. coprophila* genome.

### HTG origins may reflect shared ecology

We noted a striking commonality among the top 20 species ranked by the number of top DIAMOND hits to HTGs: 18 occupy a soil-dwelling niche, accounting for 66.1% (1,002/1,517) of all HTGs (Table S7). Putative soil-dwelling donors include collembolans (785 HTGs), bdelloid rotifers (75 HTGs), and oribatid mites (53 HTGs), all three of which are detritivores^46–48^. Although homology with sequences from living organisms cannot definitively determine the original donor species, these data suggest a bias toward acquisition from soil-dwellers in the past. Similarly to collembolans, oribatid mites, and rotifers, the larvae of *B. coprophila* and other sciarids are soil-dwelling and feed on fungal and plant matter^22,49^. These findings accord with prior reports of ecological proximity as a facilitator of HGT^2,6,9^. However, we found no evidence of widespread HGT from the assembled genome of a *B. coprophila* rickettsial endosymbiont (Figure S2B)^27,36^, which is proposed to have contributed to a germline-restricted chromosome not included in the reference annotation analyzed here^27^.

### Some HTGs may have been transferred multiple times

To characterize the deeper evolutionary history of sciarid HGT, we next examined the context around each clade of sciarid sequences in its respective gene tree (Figures 3A, S2C-J; Tables S3, S8), beyond their closest phylogenetic relatives. In 60.5% (565/934) of inferred transfer events, the sciarid clade nested within an otherwise monophyletic clade of non-sciarid sequences, a topology supporting ancient ancestry within a single donor taxon (Figure 3A1-A2, Figure S3). In contrast, 10.9% (102/934) of sciarid clades were “doubly nested,” first within a clade of non-sciarid metazoan sequences, and then within a clade of non-metazoan sequences (Figure 3A3, Figure S4). We hypothesize that these topologies may reflect two serial transfers.

**Figure 3.**
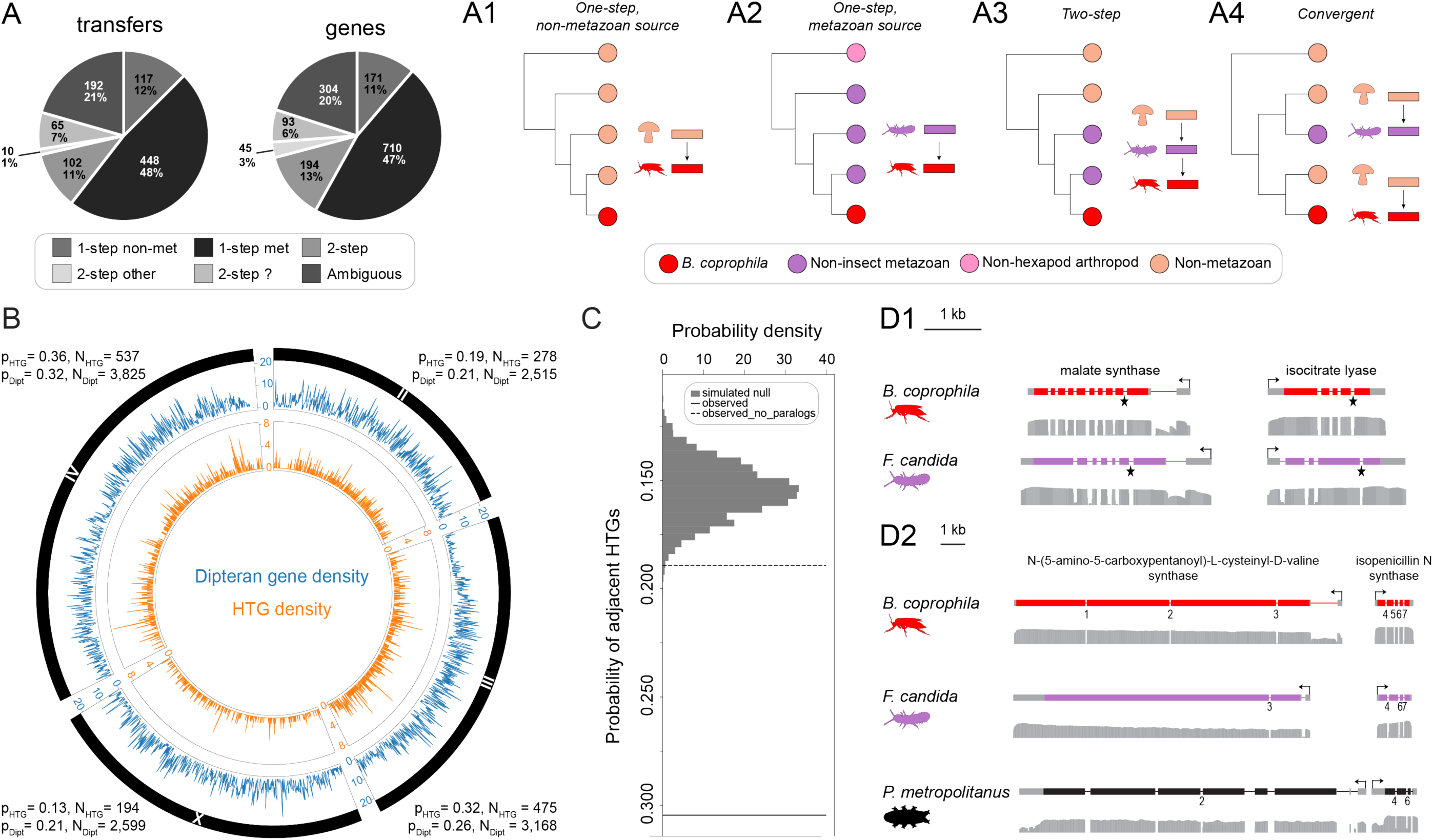
HTG genomic and evolutionary characteristics. **(A)** Schematic gene tree topologies consistent with four evolutionary scenarios: one-step transfers (A1-A2; Table S8A), two-step transfers (A3, Table S8B-C), and convergent transfers to sciarids and another metazoan taxon (A4; Table S10; 128 transfers (13.7% of all transfers) and 248 genes (16.3% of all HTGs)). Pie charts show relative contributions of transfer types to the putative HTGs detected in the *B. coprophila* genome. Categories not illustrated in this figure are shown in Figure S2C-E. “met” = metazoan; “non-met” = non-metazoan; “?” indicates unclear direction. **(B)** Circular genome diagram showing the number of HTGs (inner ring) and dipteran genes (outer ring) per 100 kb on each of the four *B. coprophila* chromosomes. Numbers outside each sector show the proportion and number of HTGs and dipteran protein-coding genes found on each chromosome, excluding 33 HTGs and 274 dipteran genes not assigned to a chromosome. **(C)** Observed proportions of HTGs immediately adjacent to other HTGs including (solid line) and excluding adjacent paralogs (dashed line), compared to a randomized null distribution (gray shaded region). **(D)** NCBI genome data tracks (not to scale) for adjacent genes putatively encoding consecutive enzymes of the glyoxylate shunt (D1) and the β-lactam biosynthesis pathway (D2) in *B. coprophila* and other animal recipients or donors of these HTGs (accessions in Table S25). Thick colored bars show CDS exons; thick gray bars show untranslated exon regions; thin lines indicate introns; arrows indicate the direction of transcription. Histograms below each transcript model indicate aggregate RNA-Seq exon coverage. Numbers below each transcript model in D2 indicate coding region-spanning introns for *B. coprophila* loci; conserved intron positions with respect to the homologous *B. coprophila* locus for *F. candida* and *P. metropolitanus* loci. The single conserved intron for each gene in the glyoxylate shunt cluster (D1) is indicated with a star. For isocitrate lyase, the intron position differs by one nucleotide position between the two homologous loci.

We interpret 80.4% (82/102) of doubly nested topologies (8.8% of all transfer events) as transfers from non-metazoans to collembolans, followed by transfers from collembolans to sciarids. Further supporting this inference, 15 of 34 *B. coprophila* HTGs with top matches in the collembolan *Folsomia candida*, as well as phylogenies consistent with an ultimate origin in bacteria, have at least one shared intron position with their collembolan homolog^48^ (Table S9). Since most bacterial genes lack introns^50^, we hypothesize that these conserved introns were acquired following a non-metazoan-to-metazoan transfer event. This intron conservation is consistent with homology between the *F. candida* and *B. coprophila* sequences, supporting a two-step transfer.

We also observed that 43.5% (128/294) of HGT events from non-metazoans to sciarids (13.7% of all transfers) co-occur in the same gene tree as a putative independent transfer to a different animal taxon (Figure 3A4; Figure S5; Table S10), most commonly rotifers (39.1%; 50/128 events). Further, 72/1,517 *B. coprophila* HTGs share predicted functions with genes that have been previously reported to have undergone HGT to diverse non-sciarid eukaryotes (Table S11). These observations suggest that some types of genes have been transferred to both sciarids and other eukaryotes.

### HTGs have non-random genomic localization

HTGs and dipteran genes are differentially distributed across chromosomes (contingency test *χ*^2^(3) = 73.5; p-value = 7.42×10^-16^), with HTGs most strongly underrepresented on the X chromosome and overrepresented on chromosome three (Figure 3B). We found that 30.5% (462/1,517) of HTGs are adjacent to other HTGs (Table S12), which exceeds the expectation under a randomized distribution of HTGs across the genome (randomization test p-value < 10^-4^; Figure 3C), even after exclusion of adjacent paralogs (p-value 2.8×10^-3^). This suggests that some genomic regions may have a higher propensity to acquire or retain HTGs, or that HGT sometimes involves the acquisition of multi-gene segments.

We wondered whether, analogously to operons and fungal gene clusters^51,52^, *B. coprophila* HTG clusters might contain functionally linked genes. Four (of 199) HTG clusters encode enzymes predicted to participate in the same biochemical pathway (Table S13). Two of these clusters, putatively encoding successive enzymes in the glyoxylate shunt (SI Note 2) and β-lactam antibiotic biosynthesis (Figure S6), shared intron-exon architecture and synteny with previously characterized homologs from the collembolan *F. candida*^53^ (Figure 3D1) as well as, in the latter case, the soil-dwelling tardigrade *Paramacrobiotus metropolitanus* (Figure 3D2)^54^. Such shared genic architecture and phylogenetic affinity (Figures S4F-G; S7A-B) among these clusters may reflect the acquisition and “domestication” of bacterial operons^51,52^ or fungal clusters followed by inter-animal exchange. We speculate that such clustering may facilitate the coordinated acquisition or regulation of these functionally linked HTGs^51,52^.

### HTGs are transcribed

We processed available poly(A)-selected RNA-Seq datasets^36,55,56^ for *B. coprophila* (Table S14) and found that 75.2% (1,141/1,517), 36.7% (557/1,517), and 6.9% (105/1,517) of HTGs are expressed at average levels greater than one, ten, and 100 transcripts per million (TPM), respectively (Figure 4A-B). As previously reported in other organisms^2,57^, the median TPM level for HTGs (5.1) is significantly lower than that of dipteran genes (18.5; two-sided Mann-Whitney *U* = 1.19×10^7^; p-value = 7.85×10^-64^).

**Figure 4.**
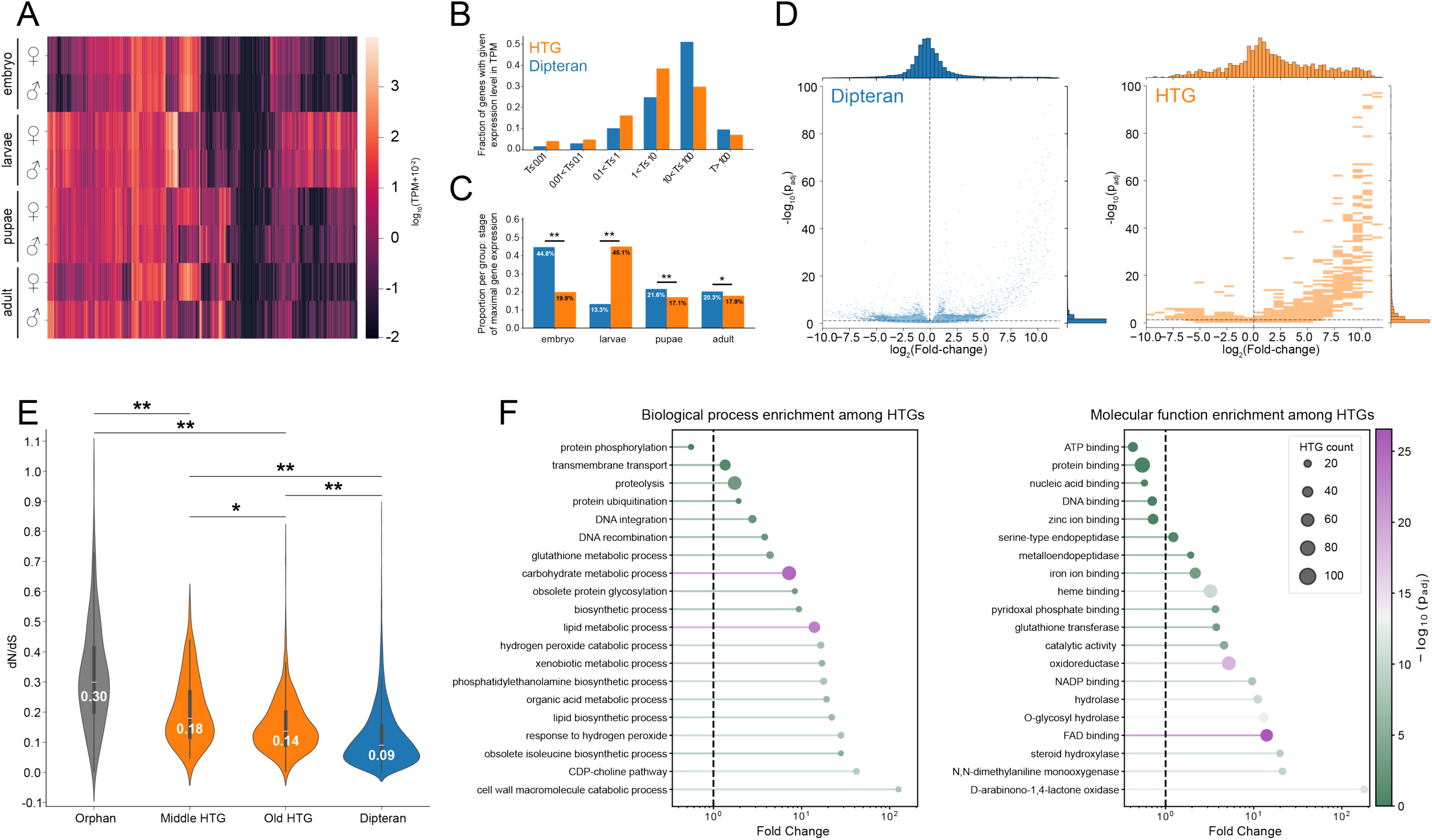
HTGs exhibit signatures consistent with biological functionality. **(A)** Heatmap of log-transformed mean RNA-Seq expression level in transcripts per million (TPM) by sex and stage for HTGs. **(B)** Mean expression level in TPM for dipteran genes and HTGs. **(C)** Stage of maximal mean expression level by gene ancestry. *χ*^2^(1) contingency test p-values from left to right are 5.97×10^-74^, 4.79×10^-208^, 7.06×10^-5^, and 0.03; *χ*^2^ values are 331.0, 947.0, 15.8, and 4.5. ** p<10^-4^, * p<0.05. **(D)** Volcano plots of differential expression between larval and non-larval samples, for dipteran-ancestry genes (left) and HTGs (right). Y-axes indicate −log₁₀ Benjamini-Hochberg-adjusted p-values; x-axes indicate log₂ fold change (larval/non-larval). **(E)** Pairwise dN/dS estimates (median values indicated under white median line) obtained for pairs of orthologous *B. coprophila* and *B. impatiens* genes separated by ancestry, with HTGs assigned to “middle” or “old” phylostratigraphic age. “Orphan” is as in Figure 1A. ** p<10^-4^, * p<0.05 under two-sided Mann-Whitney test; exact p-values left to right are 1.31×10^-11^, 1.07×10^-89^; 3.66×10^-4^, 1.63×10^-15^, 1.42×10^-72^; *U* statistics left to right are 9.14×10³, 7.19×10⁴, 2.61×10⁴, 1.88×10⁵, and 3.07×10⁶. . **(F)** Gene Ontology (GO) biological process (left) and molecular function (right) enrichment among HTGs. Circle size indicates the number of HTGs in a given gene set. Circle color indicates the Benjamini-Hochberg-adjusted p-value from Fisher’s exact test. X-axis position indicates fold-change enrichment over the genome-wide non-HTG background. See Table S18 for full data.

HTGs are expressed in multiple developmental stages and in both sexes (Figure 4A), as well as across multiple tissues and organs (Figure S8C). However, HTGs differ from dipteran genes in their spatiotemporal expression patterns (Figures 4C, S8D). Notably, 45.1% (661/1,466) of HTGs expressed with > 0 TPM in at least one stage attain their highest expression level in larvae, compared to 13.3% (1,624/12,205) of dipteran genes (Figure 4C; contingency test *χ*^2^(1): 947.0; p-value 4.79×10^-208^). Similarly, 39.6% (600/1,517) of HTGs were significantly upregulated (DESeq2 log_2_ fold change > 0, p_adj_ < 0.05) in larval vs. non-larval stages, compared to 14.2% (1,753/12,381) of dipteran genes (Figure 4D; contingency test *χ*^2^(1) = 617.8; p-value 2.27×10^-136^). These data suggest that many HTGs in *B. coprophila* are transcribed at a level consistent with biological function, and that HTGs may contribute to larval physiology.

### HTGs have been conserved under purifying selection

To characterize the temporal dynamics of *B. coprophila* horizontal transfer, we built and fossil-calibrated a sciarid species tree (Figure 1H) and assigned the minimum age of each HGT event via phylostratigraphy^58^ (Tables S3, S15). This analysis suggested that 57.1% (533/934) of transfers occurred prior to the divergence of *Bradysia* from the sciarid *Pseudolycoriella hygida*, which we estimate via a fossil-calibrated species tree (SI Methods) to have occurred 73.2 million years ago (MYA) (95% highest posterior density 50.0-98.5 MYA). Next, we analyzed the dN/dS ratio for pairwise codon alignments of *B. coprophila* HTGs with their one-to-one orthologs in *B. impatiens*^38^. We found that 99.9% (989/990) of analyzed ortholog pairs have evolved under purifying selection (dN/dS < 1; likelihood ratio test p-value < 0.05; Table S16). Combined with our inference of HTG persistence over tens of millions of years, this result is consistent with the hypothesis that HTG expression yields functional proteins.

HTGs also showed a significantly elevated rate of evolution relative to dipteran genes (median dN/dS 0.14 vs. 0.09; two-sided Mann-Whitney *U* statistic = 3.27×10^6^; p-value = 3.58×10^-84^), even when controlling for differences in RNA expression level, subcellular localization, and coding region length (Table S17). Younger HTGs have a higher median dN/dS than older HTGs (0.18 vs. 0.14; two-sided Mann-Whitney p-value = 3.66×10^-4^; Figure 4E), which may reflect transient adaptation or relaxation of constraint following acquisition, although accelerated evolution may bias our age estimates.

### HTGs have diverse predicted functions

We next turned to the prediction of HTG function. Gene Ontology (GO) assignments^36^ revealed that, relative to all other *B. coprophila* genes, HTGs are enriched in terms related to macromolecule catabolism, redox metabolism, and biosynthesis (Figure 4F). Notably, GO terms associated with transposons, a widespread source of inter-animal HGT^8,9^, collectively constitute only 3.1% (18/584) of HTGs with at least one GO biological process assignment. HTGs are predicted to localize to all subcellular compartments but, relative to all other *B. coprophila* genes, are enriched (fold change > 1.71; Fisher’s exact p_adj_ < 10^-10^) in predicted extracellular and endoplasmic reticulum localizations (Figure S8B; Table S18).

To further characterize HTG function, we manually assigned HTGs to putative biological pathways using domain annotations with InterProScan^59^, with additional evidence from existing expression data and functional assays as available (SI Note 2; Tables S18-19). Function predictions ranged from the catabolism of carbohydrates (Figure S9) to cell signaling, secondary metabolite biosynthesis, transmembrane transport, and epigenetic regulation (126, 63, 56, 68, and 27 HTGs, respectively). Consistent with prior reports that HTGs may contribute to physiological stress responses^1,3,7,31,60–64^, their domain architectures predict that *B. coprophila* HTGs participate in DNA repair, proteostasis, redox homeostasis, cofactor and antioxidant biosynthesis (Figure S10), xenobiotic metabolism/toxin resistance, and immune defenses (16, 58, 67, 34, 115, and 23 HTGs, respectively).

From these manually annotated HTGs, we selected eight HTGs of metazoan phylogenetic affinity lacking homologs in non-sciarid dipterans (Table S20; Figure S4D) for detailed examination and functional analysis. We hypothesized that these genes might confer novel functionality in *B. coprophila.* For each candidate, we detected expression in *B. coprophila* larvae using RT-PCR and used Sanger sequencing to confirm that all amplicon sequences matched the sequence of the annotated HTG (Table S14). We also assessed conservation of catalytic residues by alignment with crystal structures of homologs with experimentally demonstrated functionality (Figure S11). We then interrogated the functions of each HTG when introduced into the yeast *Saccharomyces cerevisiae* and the fly *Drosophila melanogaster*. In the following sections, we provide *in vivo* evidence that these HTGs have enzymatic functionality that we predict cannot be conferred by extant vertically inherited dipteran genes, consistent with the hypothesis that horizontal acquisition of new genes from other animals may have endowed *B. coprophila* with novel physiological abilities.

### HTGs restore NAD^+^ biosynthetic activity

We found HTGs with predicted function in the biosynthesis of four antioxidants or antioxidant cofactors (Figures 5A and S10; SI Note 2; Tables S21 and S22), including NAD^+^. From these pathways, we selected the kynurenine pathway, which functions in tryptophan catabolism and *de novo* synthesis of NAD^+^ across eukaryotes^65^, for further characterization. In most insects, only the tryptophan catabolic steps are active due to inferred loss of the three lower pathway enzymes in the last common ancestor of insects; thus, NAD^+^ must be synthesized from dietary niacin^66,67^. We predicted that *B. coprophila* HTGs encode enzymes that catalyze eight reactions in tryptophan and NAD^+^ metabolism (Figure 5A), including all three ancestrally lost enzymes in the biosynthetic pathway. The closest non-sciarid relatives of the latter three HTGs are from mites (arachnids) for 3-hydroxyanthranilate 3,4-dioxygenase (XP_037051528.1; HAAO; Figure S3C) and quinolinate phosphoribosyltransferase (XP_037047821.1; QPRT; Figure S3D), and from a collembolan for kynureninase (XP_037041147.1; KYNU; Figure S7D)^65^. Single-gene null mutants in the *S. cerevisiae* orthologs of all three genes (*bna1*, *bna6*, and *bna5*, respectively) produced pronounced growth defects on niacin-deficient growth medium (Figure 5E-G). We introduced the *B. coprophila* HTGs encoding these enzymes into *S. cerevisiae* deletion strains and performed genetic complementation assays to assess their *in vivo* functionality. We found that expression of all three HTG-encoded enzymes rescued growth of each corresponding null mutant (Figure 5E-G). This result demonstrates that the biochemical functionality of these enzymes is intact, suggesting that HTGs acquired from multiple animal sources may restore NAD^+^ biosynthesis in *B. coprophila*.

**Figure 5.**
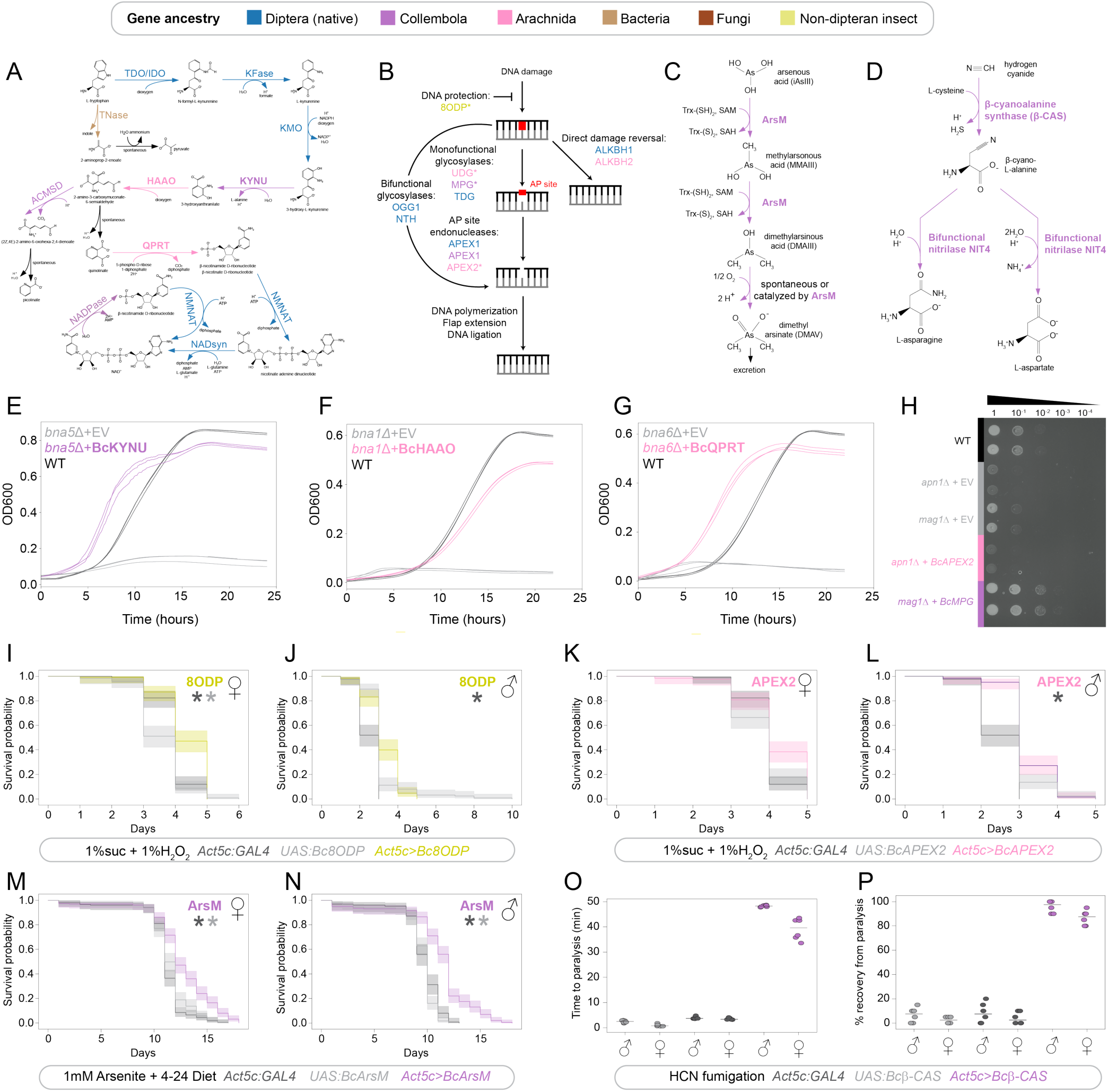
Transgenic assays demonstrate physiological activity of *B. coprophila* HTGs. **(A)** Predicted activity of HTGs and dipteran-ancestry genes involved in tryptophan catabolism and/or NAD^+^ biosynthesis (Table S21). Text and arrow colors indicate predicted enzyme-coding gene ancestry and reactions predicted to be solely catalyzed or facilitated by HTGs. Not shown for simplicity: an additional ACMSD HTG of arachnid affinity; an IDO HTG of fungal affinity; a Kfase HTG of collembolan affinity. **(B)** Schematic of putative HTG and dipteran gene activity in repair of non-bulky DNA damage (for example by oxidation, alkylation or uracil substitution; Table S24). Asterisks indicate HTGs inferred to be absent in non-sciarid dipterans. **(C)** Reactions predicted to be catalyzed by arsenite methyltransferase (ArsM). Trx-(SH)_2_ and Trx-(S)_2_ indicate reduced and oxidized thioredoxin, respectively; SAM and SAH indicate S-adenosylmethionine and S-adenosylhomocysteine, respectively. **(D)** Reactions predicted to be catalyzed by β-cyanoalanine synthase and a downstream nitrilase, both predicted HTGs. **(E)-(G)** Savitzky-Golay filter smoothed optical density (OD600) growth curves of *S. cerevisiae* colored by genotype, for KYNU ((E), *S. cerevisiae* ortholog *bna5*); HAAO ((F), *S. cerevisiae* ortholog *bna1*); and QPRT ((G), *S. cerevisiae* ortholog *bna6*). WT and EV indicate wild type and empty vector controls. **(H)** *S. cerevisiae* spot assay for *B. coprophila* APEX2 and MPG complementation of null mutants of their respective *S. cerevisiae* orthologs (*apn1Δ* and *mag1Δ*). Cell concentrations decrease from left to right across four 10-fold serial dilutions on 0.02% MMS (two replicates per genotype). **(I)-(J)** Survival curves for *D. melanogaster* exposed to hydrogen peroxide, colored by genotype for the 8ODP transgene for females (I) and males (J). Colored curves indicate constitutive, ubiquitous transgene expression; dark gray and light gray indicate driver-only and UAS-only controls. Asterisks: p < 0.05 and hazard ratio <1 for the protective effect of the ubiquitous expression compared to control genotypes (indicated by asterisk color) via Cox proportional hazards regression. **(K)-(L)** Survival curves as in (I)-(J) for the APEX2 transgene in females and males, respectively. **(M)**-**(N)** Survival curves of *D. melanogaster* exposed to arsenite, for females (M) and males (N). Colors and symbols are as in (I)-(L). Shaded regions in I-N indicate 95% confidence intervals. **(O)** Time to paralysis of all flies per replicate following HCN fumigation. **(P)** Percentage of flies recovering overnight following HCN fumigation.

### HTGs contribute to DNA repair

InterProScan annotations predicted that 16 *B. coprophila* HTGs participate in DNA repair. We selected three of six such genes predicted to participate in the repair of oxidized or alkylated nucleotides (Figure 5B) for *in vivo* assessment of DNA repair function. XP_037036837.1 encodes a methylpurine DNA glycosylase (MPG)^68^ of collembolan affinity (Figure S7E); XP_037042473.1 encodes a member of apurinic/apyrimidinic (AP) site endonuclease family 2 (APEX2)^69^ of mite (arachnid) affinity (Figure S3E); and XP_037049870.1 encodes an oxidized purine nucleoside triphosphate hydrolase (8ODP)^70^ of butterfly and moth or lacewing (insect) affinity (Figure S7C). All three enzymes are found across the tree of life but were likely lost in or before the dipteran common ancestor (Table S20).

First, we performed complementation assays with MPG and APEX2 in *S. cerevisiae* (Figure 5H). Null mutations in the *S. cerevisiae* orthologs of either gene (*MAG1*^71^ and *APN1*^72^, respectively) caused a pronounced growth defect on exposure to the methylating agent methyl methanesulfonate. While *B. coprophila* APEX2 did not rescue the *APN1* growth defect, the *MAG1* growth defect was rescued by *B. coprophila* MPG. This suggests that at least one HTG of collembolan affinity retains a function in DNA repair.

Next, we created transgenic *D. melanogaster* lines carrying APEX2 or 8ODP under UAS control^73^. Inducing expression of either *B. coprophila* gene using a constitutive, ubiquitous GAL4 driver reduced hydrogen peroxide-induced mortality (hazard ratio 0.28-0.83). The statistical significance of the effect varied by sex and choice of parental control (Figure 5I-L; Table S23). APEX2 and 8ODP expression also promoted female survival without hydrogen peroxide exposure (hazard ratio 0.48-0.72; Figure S12A, S12C), as has been observed for overexpression of some endogenous DNA repair genes in *D. melanogaster*^74^. Collectively, these results are consistent with the hypothesis that HGT from animals contributes to DNA repair in *B. coprophila*.

### An HTG enables arsenite tolerance

We predicted that 115 HTGs participate in detoxification or toxin resistance. Among these, XP_037042500.1 is a predicted arsenite methyltransferase (ArsM) of collembolan affinity (Figure S4E). ArsM detoxifies inorganic arsenite via successive methylation reactions (Figure 5C)^75^. Thus, ArsM acquisition could have endowed *B. coprophila* with the ability to withstand arsenite toxicity. To test this hypothesis, we generated transgenic *D. melanogaster* carrying the *B. coprophila* ArsM gene under UAS control. Constitutive ubiquitous expression of the transgene did not significantly improve survival on standard *D. melanogaster* diet (Figure S13, Table S23). Adding 1 mM sodium arsenite to standard diet killed all male control flies within 13 days (Figure 5N) and all female control flies within 17 days (Figure 5M). However, driving expression of *B. coprophila* ArsM with a ubiquitous constitutive GAL4 driver delayed complete mortality on sodium arsenite by between one (females) and five (males) additional days (hazard ratio 0.20-0.49; p-value < 1×10^-5^; Figure 5M-N, Table S23).

### An HTG enables cyanide tolerance

Five HTGs of collembolan phylogenetic affinity were predicted to enable tolerance of cyanide. We ubiquitously expressed one of these, which we predicted to be a β-cyanoalanine synthase (XP_037050347.1; Figure S4D)^7^, in *D. melanogaster*, and tested its predicted detoxification activity (Figure 5D) using a cyanide fumigation assay^76^. Following fumigation, we found that transgene expression with a ubiquitous constitutive GAL4 driver increased the median time to paralysis by at least 11.8-fold (Figure 5O; Video S1) and the probability of overnight recovery by at least 13.0-fold (Figure 5P) in both sexes.

## Discussion

We provide evidence for an unprecedented level of HGT within an animal genome, even more notable because at least 78.3% of the HTGs likely originate from other animals (Figure 2A-B; Figure S2A). The observed proportion of protein-coding HTGs (9.2%) is comparable to the levels observed in prokaryotic genomes^4^, while other eukaryotes have on average no more than 5% of their protein-coding genome derived from HGT^15,77,78^. To our knowledge, the *B. coprophila* genome contains a greater proportion and number of HTGs than any previously analyzed animal genome^2,15,79^. Although we cannot completely rule out phylogenetic artifacts, contamination, or gene loss as alternative explanations^1^, our conservative detection and validation approach (Figure 1; Figure 2C-E) suggests that these phenomena are unlikely to fully account for the signature of massive HGT in this genome.

We have further shown that multiple *B. coprophila* HTGs are biochemically active when expressed in yeast and *Drosophila*, supporting the hypothesis that they have contributed to novel stress-tolerance and biosynthetic physiology. Our findings that HGT restored ancestral losses of the DNA repair enzyme MPG (Figure 5H) as well as NAD^+^ biosynthesis in *B. coprophila* (Figure 5A, E-G) show that inter-metazoan HGT may circumvent the irreversibility of gene and complex-phenotype loss in animals, as has previously been reported for fungi^51,80^. In sum, inter-animal HGT, previously only rarely documented^3,12–14^, appears to be a major force of genomic and physiological change in sciarids.

We speculate that sciarid ecology may have favored the fixation and retention of HTGs (Figure 6). HTGs are disproportionately transcribed in *B. coprophila* larvae (Figure 4C-D), which is the soil-dwelling stage and the primary feeding stage of the sciarid life cycle^49^. We predict that HTGs participate in carbohydrate catabolism (126 HTGs; Figure S9), antibiotic biosynthesis (Figure S6), and toxin resistance (115 HTGs), all of which may be adaptive functions in detritivorous and herbivorous niches (SI Note 2)^7,23,25,40,48,53,61,81^. In particular, sciarids are likely to encounter arsenite as a natural component of soil^82^ and cyanide as a plant toxin^7^, and our *Drosophila* transgenic assays show that single HTGs are sufficient to confer resistance to both (Figures 5C, D, M-P). Recent work also suggests that an HTG confers cyanide-resistant cellular respiration in the sciarid *P. hygida*^22^.

**Figure 6.**
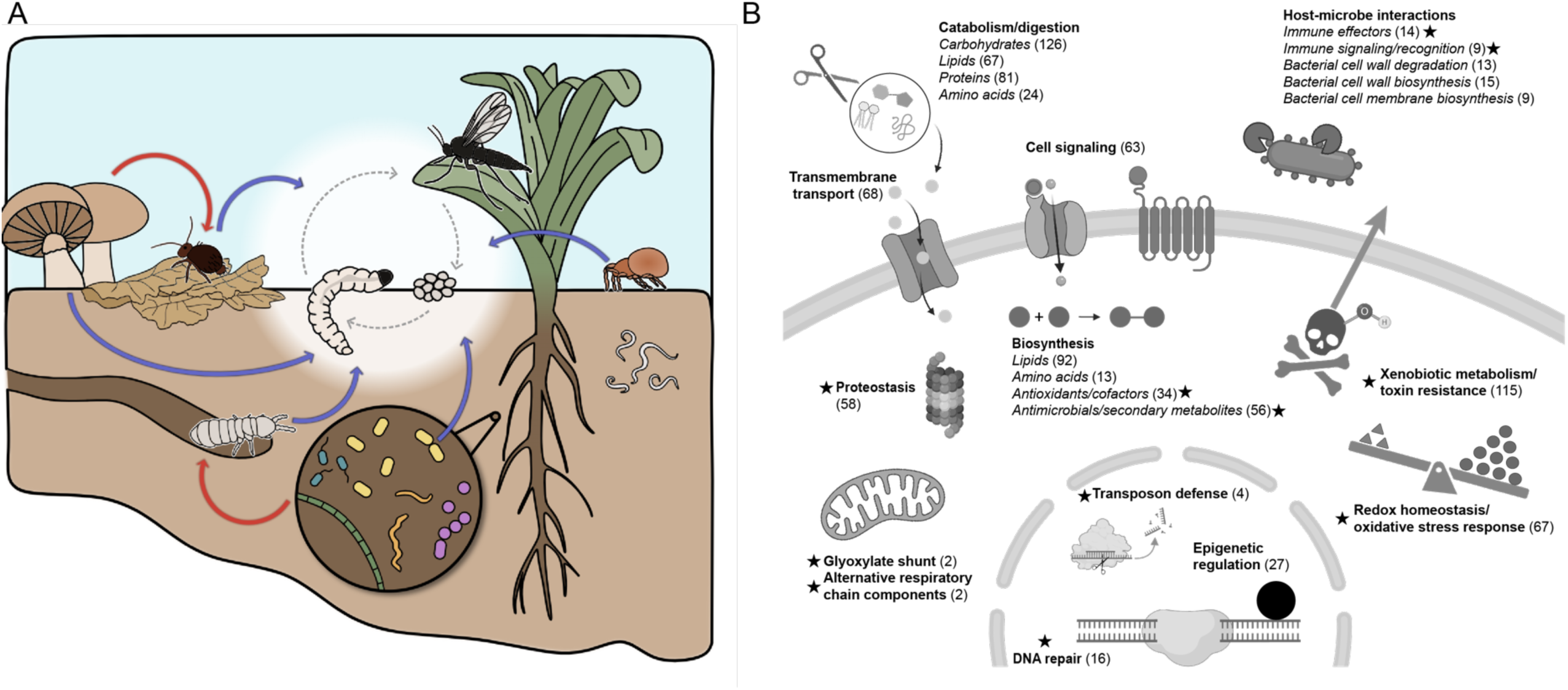
Conceptual model for the facilitators and consequences of HGT in *B. coprophila*. **(A**) Sciarids (white circle) occupy a soil-dwelling niche where they acquire HTGs from other soil-dwelling microbes, fungi, and invertebrates (blue arrows). Red arrows indicate putative transfers from non-animals to other invertebrates, which may be followed by transfer to sciarids. **(B)** Illustration of subcellular localizations and predicted biological functions of manually annotated HTGs (SI Note 2). Stars indicate functions putatively associated with stress responses; the number of HTGs associated with each process (Table S19) is indicated in parentheses; italicized text indicates subprocesses within the broader process in bold text.

The soil environment could also subject sciarids to frequent oxidative and/or DNA damage from desiccation^83^, transient hypoxia^22,84^, or redox chemistry^85^. *Bradysia* tolerates a higher level of hypoxia^86^ and ionizing radiation^55^ than other arthropods, and HTGs constitute 11.8% (40/338) of *B. coprophila* genes upregulated in response to ionizing radiation^55^. Thus, HGT may have enhanced DNA repair and/or redox homeostasis in this animal. Consistent with this hypothesis, we experimentally validated biochemical activity of HTGs in the biosynthesis of NAD^+^ (Figure 5A, E-G), a cofactor that supports oxidative stress and DNA-damage responses^87^, and the responses to DNA-alkylating as well as oxidative stressors (Figure 5B, H-L).

Notably, carbohydrate catabolism^25,40^, detoxification^88,89^, and responses to oxidative^81^ and DNA damage^60^ have putatively been enhanced by HGT in other transitions to herbivorous or soil-dwelling niches^6,7^. Thus, repeated horizontal acquisition of genes conferring niche-associated functions (Table S11) may facilitate ecological convergence across diverse animals^6,7,42^. Taken together, these results add to the emerging paradigm that HGT plays an important role in animal innovations including niche adaptation^3,6,7^. Our findings expand the functional relevance of HGT beyond prokaryotes and show that animal-to-animal HGT can be both prevalent and biologically relevant.

What factors might have enabled massive HGT in sciarids? Proximity to soil-dwelling donor lineages likely contributed (Table S4; Figure 6A). Collembolans, mites, and rotifers are thought to possess genomic adaptations to life in soil, including HTGs from microbes^16,23,47,48,53,60,61,81^. Our phylogenetic analyses suggest that multi-step networks of genetic exchange among these organisms and sciarids may also contribute to adaptation to this shared niche (Figures 6A, S4, S7; Tables S8B-H). We speculate that ecological factors including the stabilization of extracellular DNA by adsorption to soil particles^90^, horizontal transmission of cargo-carrying transposons or viruses^10^, or frequent DNA damage caused by soil-related stressors (see above)^17,83^ might contribute to the apparent frequency of HGT in the soil.

To conclude, our results suggest that inter-animal HGT cannot be overlooked as a source of genomic and physiological novelty in animal evolution. We anticipate that future work will uncover more examples of genetic exchange among animals, especially those that inhabit stress-inducing environments.

## Methods

Bioinformatic and experimental analyses are described in brief here, with additional detail provided in SI Methods.

### HGT inference

#### DIAMOND-based HGT inference

The longest isoform for each *B. coprophila* protein-coding gene was queried against the clustered non-redundant (clusteredNR) database (https://ftp.ncbi.nlm.nih.gov/blast/db/experimental/, access date 5/26/2026) with DIAMOND (v.2.2.0 --evalue 0.01, very-sensitive mode). Protein-coding genes were assigned to one of five ancestry annotations (Table S6) depending upon the taxonomic distribution of the top 2,000 DIAMOND hits (Figure 1A) as follows:

- *orphan*: no significant (E-value < 0.01) hits outside of the taxonomic family of the query.
- *non_dipt* (HTG 1 in Figure 1A): significant hits (E-value < 10^-5^) in non-dipteran lineages and no significant (E-value < 0.01) hits in Diptera outside the family of the query sequence.
- *alien* (HTG 2 in Figure 1A): significant hits in other dipteran families that are more than ten orders of magnitude higher in E-value (i.e., weaker similarity) than the best non-dipteran hit, such that the “Alien Index”^16^ (AI) > 10.
- *dipt*: top hit outside of the taxonomic family of the query is in a different dipteran family. These genes are inferred to be of vertical descent.
- *none*: all others; i.e., the top hit outside of the taxonomic family of the query is non-dipteran, with AI < 10 and/or E-value > 10^-5^.

Hits to synthetic sequences (NCBI taxid 32630) and the genome of *Rhagoletis zephyria* (NCBI taxid 28612), previously shown to be contaminated with reads from mites^91^, were excluded from the above calculations and subsequent phylogenetic inference. We modified the previously derived Alien Index (AI)^16^ to allow for HTG candidates of non-dipteran origin, as follows:

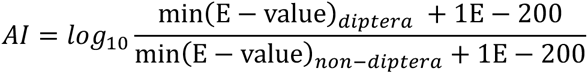

Here, *diptera* refers to hits in Diptera outside of the family of the query sequence, and *non* − *diptera* refers to any hit outside of Diptera, including both metazoan and non-metazoan clades.

Genes assigned to the *non_dipt* or *alien* categories were considered to be of putative HGT ancestry. We note that our AI cutoff of ten is conservative and may undercount the number of HTGs, especially from other arthropods. Accordingly, one cytochrome P450 sequence previously identified via phylogenetic analysis as an HTG of arthropod origin (XP_037052058.1)^31^ failed to pass our threshold (Table S1). Comparisons to dipteran-ancestry genes in all downstream analyses refer to the *dipt* category. The above criteria were also applied to the longest predicted protein isoform per protein-coding gene for 53 non-*Bradysia* dipteran species with RefSeq annotations of chromosome-level or scaffold-level assemblies (genome accessions in Table S2).

#### Phylogenetic inference

DIAMOND hits for putative HTGs were similarity-clustered prior to gene tree inference. Phylogenetic datasets were constructed to include protein sequences from *B. coprophila*, other members of the same taxonomic family (Sciaridae), and non-sciarid homologs retrieved via DIAMOND. Genome annotations for other sciarids were obtained from GenBank if available or generated via BRAKER3^92^ if no annotation was available. Sequences were aligned with MUSCLE (v.5.1)^93^, followed by alignment trimming with trimAl (v.1.4.1)^94^, and maximum-likelihood gene tree inference with IQ-TREE (v.2.2.0.3)^95^. Trees were subjected both to automated analysis of closest phylogenetic relatives on midpoint-rooted trees (Figure S2A), and manual inspection of the broader phylogenetic context surrounding each clade of sciarid sequences (Figure 3A1-A4; Figure S2C-J), including manual re-rooting to reflect species relationships when applicable (55 cases). Each (potentially multi-sequence) clade of sciarid sequences was considered to reflect an HGT event if it phylogenetically grouped with sequences from non-dipteran sources and was separated from non-sciarid dipteran sequences by at least one node with > 70% bootstrap support. We additionally excluded topologies suggestive of one-step transfer from sciarids to non-sciarids. Only HTGs supported by both DIAMOND and tree inspection were included in the final set of 1,517 HTGs.

Closest relatives of HTGs in Figures 5 and S6, S9, and S10, as well as Tables S18, S21, S22, S24, and S25 were determined from the closest non-sciarid clade on gene trees, or top non-sciarid DIAMOND hits if this clade was polyphyletic. Sciarid homologs of HTGs were defined as all sciarid sequences more closely related to a *B. coprophila* HTG than any non-sciarid sequence. Thus, a *B. coprophila* HTG and its homologs in other species were inferred to descend from a single transfer event in a sciarid ancestor. Phylostratigraphic age estimates (Figure 1H; Tables S3 and S15) were assigned according to the distribution of homologs across sciarid species.

#### Contamination analysis

Previously published Oxford Nanopore reads^36^ were mapped to the *B. coprophila* reference annotation with Minimap2 (v.2.28)^96^, SAMtools (v.1.21)^97^, and BEDTools (v.2.31.1)^98^ to compute read depth, length and co-occurrence data for HTGs and dipteran genes. Null distributions of the proportion of HTGs found in chimeric and pure HTG-containing reads in Figure 1G were produced by 1,000 iterations of a simulation in which annotated ‘dipteran’ genes were randomly relabeled as ‘dipteran’ or ‘HTG’ such that the proportion of simulated HTGs was equal to the observed proportion. Microsynteny between loci in the genomes of *B. coprophila* and other sciarids (Figure S1E-F) was assessed via reciprocal best DIAMOND BLASTp hits.

#### In silico functional analysis

Previously published *B. coprophila* RNA-Seq TPM and raw count data^36,55,56^ were accessed from NCBI. Differential expression analysis was performed in DESeq2^99^. dN/dS was computed for reciprocal best DIAMOND BLASTp hits between *B. coprophila* and *B. impatiens*, the most closely related species with a publicly available chromosome-level assembly, using PAL2NAL^100^-generated codon alignments and PAML^101^. Analysis was restricted to pairs with dS < 2, and selective constraint was measured using a likelihood ratio test (*χ*^2^(1) = 2 Δlog(*L*)) of the null hypothesis that dN/dS = 1, with false discovery-rate adjustment according to the Benjamini-Hochberg procedure implemented in statsmodels^102^ (version 0.14.0). Figure 4E is restricted to HTGs shared by *B. coprophila* and *B. impatiens*. “Middle-age” HTGs were defined as those with tree-inferred orthologs in *B. impatiens* but not in more distantly related sciarids, and “old” HTGs have tree-inferred orthologs in *Lycoriella ingenua* or *P. hygida*.

Subcellular localization was predicted for the longest protein isoform per *B. coprophila* gene using DeepLoc2.0^103^ (“accurate” model). Plastid predictions were excluded for enrichment analyses. Gene Ontology (GO) term annotations pre-computed with InterProScan^59^ version 5.72-103.0 were obtained from the NCBI FTP site for the *B. coprophila* reference genome annotation (“GCF_014529535.2-RS_2026_02_gene_ontology.gaf”, access date 7/7/2026) and mapped to their respective names using the “go-basic.obo” file obtained from the GO website (release 2026-06-15). Functional enrichments for DeepLoc2-predicted subcellular localizations, GO molecular function, and GO biological process were computed with Fisher’s exact test in SciPy^104^ (version 1.8.1), with “alternative” set to two-sided for subcellular localization and “greater” for GO terms. Enrichment computations were restricted to the longest isoform per gene, excluding all genes without at least one assigned GO term. P-values were adjusted with the Benjamini-Hochberg procedure.

The following biological process terms were considered to be potentially associated with transposable elements: GO:0006310 (DNA recombination); GO:0015074 (DNA integration); GO:0006313 (DNA transposition); GO:0098039 (replicative DNA transposition); GO:0098038 (non-replicative DNA transposition); GO:0006317 (DNA-mediated transposition); GO:0006318 (DNA-mediated transposition); GO:0032197 (retrotransposition).

All *B. coprophila* HTGs were submitted to InterProScan^59^ (version 5.77-108.0) for protein domain annotation. Biological pathways (Table S18, sheet “manual_annotations”) were manually assigned from inspection of InterPro domain descriptions (access date 7/14/2026) or the sources indicated in Table S18. Author-assigned pathways were reviewed and revised for internal consistency using assistance from ChatGPT (version 5.5, “Pro” intelligence). Hercynylcysteine sulfoxide lyase domain assignments (Figure S10A) were validated with the KofamKOALA^105^ web server (version 2026-06-01, KEGG release 118.0).

Carbohydrate-active enzyme (CAZyme) annotations were obtained for *B. coprophila* from a previous publication^40^ and mapped to the current genome annotation. We found that 258/270 (95.6%) of CAZymes identified by Trinca and colleagues^40^ from annotation version GCF_014529535.1 successfully mapped to GCF_014529535.2, of which 89 were identified as HTGs. Carbohydrate catabolism HTGs were tabulated by combining previously reported^40^ Pfam^106^ and dbCAN2^107^–based annotations (Table S18, “carbohydrate_catabolism”; Table S26). When InterProScan annotations differed from previously reported annotations^40^, priority was given to the previously reported annotation.

Protein structures were predicted with AlphaFold2^108^ using ColabFold^109^ (v.1.5.5) and aligned to solved crystal structures from the Protein Data Bank (PDB) with PyMOL^110^ (v.3.0.3).

### RT-PCR confirmation of B. coprophila HTG expression

cDNA was synthesized from mRNA extracted from seven-day-old *B. coprophila* larvae and used as a template for a PCR reaction to assess endogenous expression of putative HTGs selected for trans-species experimental analysis. Gene-specific primers were designed using the NCBI Primer-BLAST tool^111^ to span multiple introns when possible. PCR products were subjected to Sanger sequencing and compared to corresponding RefSeq annotations. Primer sequences and GenBank accessions of Sanger sequencing results are available in Table S14 (sheet “Sanger_results”).

### Transgenic assays in D. melanogaster

#### Design and synthesis of transgenic constructs

The coding sequences (CDSs) of *B. coprophila* ArsM, β-CAS, APEX2, and 8ODP (referenced with Bc prefixes) were codon-optimized for *D. melanogaster* expression using the Twist Biosciences Codon Optimization tool^112^. Each codon-optimized sequence had a (Gly4Ser)3 linker and hemagglutinin (HA) tag added to the 3’ end of the sequence and a Kozak sequence (ATCAAC ATG) at the 5’ end just upstream of the start site. These sequences were synthesized by Twist Biosciences and cloned between the XbaI and XhoI restriction sites after the removal of the myristoylated GFP from the original plasmid (pJFRC12-10XUAS-IVS-myr::GFP; Addgene #26222). Both XbaI and XhoI restriction sites were retained in the final plasmid.

#### Generation of transgenic flies

Each plasmid was integrated into the *D. melanogaster* genome via phiC31-mediated recombination into an attP site on the second chromosome (attP40)^113^. Plasmids were injected by BestGene Inc. and transgenic flies were identified by eye color.

#### Drosophila maintenance and stocks

Flies were raised at 25 °C and 60% relative humidity with standard *Drosophila* food prepared as previously described^114^. All flies used in this study were either generated herein or obtained from the Bloomington Drosophila Stock Center (BDSC) and are listed in the Supporting Information Materials “Experimental Materials” table.

#### Test for resistance to hydrogen peroxide

We adapted our protocol from methods reported by Vrailas-Mortimer and colleagues^115^. Virgin male and female flies of BcAPEX2 and Bc8ODP experimental (UAS+GAL4) and parental control (UAS-only and GAL4 driver-only) genotypes were collected by crossing virgin females of the UAS line to males of the GAL4 lines (for experimental flies), or virgin females of the UAS or GAL4 lines to Oregon R males (for control flies). Flies were aged for two days and then starved for six hours. 25 flies of the same sex were transferred into plastic vials with 1.3% agarose and 1% sucrose in reverse osmosis (RO) water (controls) or 1.3% agarose, 1% sucrose and 1% hydrogen peroxide in RO water (experimental), with five replicate vials per each combination of genotype, sex, and treatment. Survivors were counted daily and flies were transferred every four days into new vials until all the flies had died.

#### Test for resistance to arsenite

We adapted our protocol from methods reported by Ortiz and colleagues^75^. Virgin male and female flies of all BcArsM genotypes (UAS-only or GAL4 driver-only parental controls, and UAS+GAL4 experimental flies) were collected by crossing as described for hydrogen peroxide assays above. 25 one-day-old flies of the same sex were placed in vials with Formula 4-24 instant medium (controls) or 1 mM sodium arsenite-supplemented Formula 4-24 instant medium (experimental). Between five and ten independent replicate vials were analyzed per combination of sex, genotype, and condition (Table S23). Survivors were counted daily and flies were transferred every five days into new vials until all the flies had died.

#### Test for resistance to cyanide

We adapted our protocol from methods reported by Fernandez-Ayala and colleagues^76^. Virgin male and female flies of Bcop β-CAS genotypes (UAS-only or GAL4 driver-only parental controls, and UAS+GAL4 experimental flies) were collected by crossing as described for hydrogen peroxide assays above. 20 one-day-old flies of the same sex were flipped into plugged vials with 1% agarose in RO water containing 10 mM KCN, 50 mM Tris-HCl (pH 7.5) inside a fume hood at room temperature. Six replicate vials were assayed per combination of sex and genotype. Flies were kept in the vials until the last fly in all three conditions stopped moving (58 minutes; see Video S1). Flies were then transferred to vials with 1% agarose in RO water and kept overnight at 25 °C and 60% relative humidity to measure recovery. The time taken for the paralysis of all flies in each vial was measured, as well as the number of flies that recovered overnight.

#### Survival analysis

Experimental recordings of cumulative deaths per day were transformed to long-format tables in which each row corresponds to a single death or censoring. All flies alive on the last day were recorded as a separate censoring event. In this table, we enforced monotonicity in cumulative deaths to account for inadvertent mis-recording of deaths; Cox proportional hazards regression results were robust to removal of two replicates in the arsenite experiment that exhibited apparent mis-recording on a single day in each replicate.

These long-format tables were visualized as Kaplan-Meier survival curves and analyzed using Cox proportional hazards regression with lifelines^116^ (version 0.30.0). Cox regressions (Table S23) were fit with biological replicates as clusters to account for potential non-independence within vials. Comparisons of 1% H₂O₂ or arsenite-supplemented diets to unsupplemented diets were performed on pooled datasets using sex and genotype as covariates. Pairwise comparisons of UAS+GAL4 experimental flies to GAL4 driver-only or UAS-only control flies were performed on subsets of the data separated according to dietary medium and sex.

### Saccharomyces cerevisiae complementation assays

#### Plasmid construction

Putative *B. coprophila* kynurenine pathway coding sequences (CDSs) for KYNU, HAAO, and QPRT were initially codon optimized for expression in *Nicotiana tabacum*. Due to difficulties driving expression in this system (not shown), we then cloned the *B. coprophila* CDSs from the *N. tabacum* expression plasmids into a *S. cerevisiae* expression vector. The yeast expression vector pPN352^117^, with an Actin promoter (∼1,000 base pairs upstream of the start codon of the Actin gene) and an ADH1 terminator, was linearized by NotI digestion to introduce the CDS. Gibson assembly reactions were performed using pPN352-NotI as the vector and individual *B. coprophila* coding sequences as inserts to generate pPN352-BcKYNU, pPN352-BcHAAO, and pPN352-BcQPRT. Representative correct clones were confirmed by whole plasmid sequencing (Plasmidsaurus) and used for subsequent yeast transformations.

*B. coprophila* DNA repair genes (BcMPG, BcAPEX2) were amplified based on the gBlocks sequence containing yeast-codon-optimized CDSs using primers to add 2X SV40 nuclear localization sequences to the C-termini of the predicted proteins. PCR products were purified and assembled into NotI-linearized pPN352 by Gibson assembly.

#### Yeast strains and growth conditions

*S. cerevisiae* BY4741 deletion strains lacking the yeast equivalents of five *B. coprophila* HTGs were obtained from the yeast deletion collection (Research Genetics; Catalog No. 95401.H2)^118,119^ and revived from frozen stocks onto appropriate synthetic medium (CSM^120,121^) or yeast extract, peptone, dextrose (YPD^122^) plates. CSM Medium comprised of 6.7 g/L Yeast Nitrogen Base (Research Products International), 0.74 g/L Complete Supplement Mixture (CSM; Sunrise Science Products) and 2% (w/v) glucose. For CSM-URA, Complete Supplement Mixture (CSM) without Uracil (Sunrise Science Products) was used. The wild-type MATa strain RDY68 (S288C background) was used as a positive control for growth and DNA damage resistance. Cells were routinely cultured at 30 °C in synthetic medium lacking uracil (SC-URA) with 2% glucose unless otherwise noted.

#### Yeast transformation and selection

Deletion strains were transformed with the empty vector pPN352 or the indicated *B. coprophila* CDS expression plasmids using a standard high-efficiency LiAc/PEG transformation protocol^123^. Transformants were selected on CSM-URA plates and re-streaked to ensure clonality. For each genotype (empty vector vs. *B. coprophila* rescue construct), two independent transformants were chosen for downstream growth and DNA damage assays.

#### Nicotinic acid auxotrophy complementation (kynurenine pathway)

Complementation of kynurenine pathway deletions by *B. coprophila* HTGs was assayed on nicotinic-acid–free medium. Overnight cultures of empty vector, BY4741 *bna1Δ*, *bna5Δ*, and *bna6Δ* transformants grown in SC-URA medium containing nicotinic acid were harvested. 10 µL of each culture was transferred to a 96-well filter plate. Cells were pelleted at 1,800 rpm for three minutes (twice) and washed with 200 µL of post-shift medium to remove residual nicotinic acid. Washed cells were resuspended in 200 µL of yeast nitrogen base (YNB) medium lacking nicotinic acid (YNB-NA, Formedium) supplemented with SC-URA and 2% glucose, transferred to a 96-well plate, and incubated at 30 °C with orbital shaking in a plate reader. Optical density at 600 nm was recorded every six minutes for 24 hours, and growth curves were compared between empty-vector and *B. coprophila* gene expression constructs in the deletion strains, with RDY68 included as a wild-type positive control. Growth curves were plotted with a Savitzky-Golay smoothing filter in SciPy (savgol_filter window_length: 31; polyorder: 1).

#### MMS sensitivity assay (DNA alkylation repair)

To assess complementation of DNA repair defects, *apn1Δ* and *mag1Δ* strains transformed with empty vector or with BcAPEX2 or BcMPG expression constructs were tested on methyl methanesulfonate (MMS). SC-URA plates containing 2% glucose and 0.02% MMS were prepared by adding MMS to cooled SC-URA agar just before pouring (35 mL per omniplate). Overnight cultures were adjusted to the same OD600, and four 10-fold serial dilutions were prepared in sterile medium. Five µL of each dilution was spotted onto control and MMS-containing plates, which had been dried in a laminar-flow hood for five minutes before spotting. Plates were incubated at 30 °C and imaged after 48 hours to assess the relative survival of each strain–plasmid combination.

#### Software access and use of artificial intelligence

Except where otherwise noted in “Supporting Information Materials: Software and algorithms,” all bioinformatics software was accessed via Singularity images distributed by the BioContainers project^124^ on the Harvard Faculty of Arts and Sciences Research Computing cluster. ChatGPT (versions 4-5.5) was used to assist with scripting from human-generated pseudocode; review, commenting, and organization of human-generated code; and proofreading of the manuscript for spelling, grammar, and consistency. All AI-assisted code was reviewed, revised, and interpreted by R.R.K.. Figures were assembled with https://BioRender.com/ (RRID:SCR_018361) and Adobe Illustrator v30.1. Sankey diagrams were created using SankeyMATIC^125^.

## Data availability

Input data as well as intermediate and final data outputs have been deposited as a Dryad repository (10.5061/dryad.5dv41nsmx). Sanger sequencing results are available on GenBank (accessions provided in Table S14). Gene trees for selected HTGs are provided as supplementary Figures S3, S4, S5, S7 and S14, and all gene trees are available on iTOL (https://itol.embl.de/shared/20NWy5j2znqY5). All code required to reproduce these analyses is available at https://github.com/rishabhrajkapoor/Bradysia_coprophila_HGT (commit ID: 594a074).

## Acknowledgments

We are grateful to Susan Gerbi (Brown University) for providing us with *B. coprophila* cultures, knowledge and feedback on the manuscript; Sean Eddy (Harvard University), Gonzalo Giribet (Harvard University), Nadia Monesi (University of São Paulo), and Thomas Niehaus (University of Minnesota) for stimulating discussion and feedback; Ryan Nett and Fabiola Muro Villanueva (Harvard University) for kynurenine pathway plasmid design and initial heterologous expression attempts using *N. tabacum*; Extavour lab members Sarah Frail, Evelyn Schwager, Olakunle Jayesimi, and Yu Shirai for assistance with experiments and discussion; Sarah Frail for the illustration in Figure 6A; and Robert Baird and the Yukiko Yamashita lab (Whitehead Institute) for *B. coprophila* culture assistance and discussion. Computational resources were provided and maintained by Harvard Faculty of Arts and Sciences Research Computing. This work was supported by an NSF Graduate Research Fellowship Program award 2023356057 to R.R.K.; award RGP0041/2022 from the Human Frontier Science Program (S.P.) to C.G.E.; grant NIH/NIGMS R01 GM043987 to A.W.M.; a Simmons Award to P.N.; a grant from the NSF-Simons Center for the Mathematical and Statistical Analysis of Biology (NSF #1764269, Simons #594596) to A.W.M.; the Howard Hughes Medical Institute; and funds from Harvard University. C.G.E. is an Investigator of the Howard Hughes Medical Institute.

## Author contributions

Project ideation: R.R.K. and C.G.E.; computational analyses: R.R.K.; wet lab experiments: T.K., P.N., S.P.; writing: R.R.K., C.G.E., T.K., P.N.; project supervision: C.G.E. and A.W.M. (yeast experiments).

