## Supplementary Information File for "Massive inter-animal horizontal gene transfer contributes to stress tolerance in the fungus gnat *Bradysia coprophila*"

This file contains:

- SI Table of Contents
- SI Figures S1-S14
- SI Video S1 and Legend
- SI Notes 1-2
- SI Materials: Software and Algorithms
- SI Materials: Experimental Materials
- SI Methods
- SI Tables S1-S30 and SI Table Legends
- SI References

### SI Table of Contents

| | |
|---|---|
| **SI Figures S1-S14** | |
| Figure S1. Bioinformatic characteristics of HTGs. | 4 |
| Figure S2. Supporting data for DIAMOND and phylogenetic inference of HTG origins. | 5 |
| Figure S3. Examples of gene trees supporting one-step transfers. | 6 |
| Figure S4. Examples of gene trees supporting two-step transfers. | 7 |
| Figure S5. Examples of gene trees supporting convergent HGT events to sciarids and other metazoan genomes. | 9 |
| Figure S6. HTGs may enable the synthesis of β-lactam antibiotics. | 10 |
| Figure S7. Examples of gene trees of “ambiguous” topology. | 11 |
| Figure S8. Additional data related to HTG conservation and predicted functions. | 12 |
| Figure S9. HTGs may contribute to the catabolism of plant, fungal, and bacterial cell wall-derived polysaccharides. | 13 |
| Figure S10. HTGs may contribute to multi-step antioxidant and cofactor biosynthetic pathways. | 14 |
| Figure S11. Alignment of AlphaFold2 predictions for HTGs and solved homolog structures. | 15 |
| Figure S12. *D. melanogaster* survival curves related to APEX2 and 8ODP transgenic expression. | 23 |
| Figure S13. *D. melanogaster* survival curves related to ArsM transgenic expression. | 24 |
| Figure S14. Examples of gene trees supporting gene family expansion via HGT. | 25 |
| **SI Video S1** *D. melanogaster* cyanide fumigation assay of β-CAS activity. | 26 |
| **SI Notes** | |
| SI Note 1. HGT contributes to the expansion of gene families. | 27 |
| SI Note 2. HTGs have diverse predicted functions in *B. coprophila* stress responses, metabolism, and ecology. | 28 |
| *Macromolecule catabolism and digestion* | *28* |
| *Transmembrane transport* | *29* |
| *Detoxification/xenobiotic metabolism* | *29* |
| *Proteostasis* | *30* |
| *Epigenetic regulation and transposon defense* | *30* |
| *Redox homeostasis/oxidative stress responses* | *30* |
| *DNA repair* | *31* |
| *Biosynthesis* | *31* |
| *Glyoxylate shunt* | *32* |
| *Cell signaling* | *33* |
| *Immunological defenses, protein toxins, and host-microbe interaction* | *33* |
| **SI Materials: Software and Algorithms** | **35** |
| **SI Materials: Experimental Materials** | **37** |
| **SI Methods** | **39** |
| Phylogenetic inference | 39 |
| *Sciarid database construction* | *39* |
| *Maximum likelihood gene tree construction* | *39* |
| *Automated gene tree analysis* | *40* |
| *Manual gene tree inspection and HGT inference* | *41* |
| *Analysis of intron position conservation for two-step transfers* | *42* |
| *Analysis of putative convergent transfers to other metazoans* | *43* |
| *Gene family expansion analysis: post-acquisition gene duplication* | *43* |
| *Gene family expansion analysis: multiple putative independent transfers to the B. coprophila genome* | *44* |
| *Expansions of gene families via HGT of genes with pseudoparalogs in the B. coprophila genome* | 44 |
| Analysis of HTG synteny between B. coprophila and other sciarid genomes | 44 |
| Search for possible HGT donor sequences in a putative *B. coprophila* endosymbiont genome | 45 |

| | |
|---|---|
| Liftoff annotation transfer | 45 |
| Analysis of Oxford Nanopore (ONT) reads | 45 |
| Nucleotide composition analyses | 45 |
| Arthropod lineage tree construction | 46 |
| Analysis of alternative hypothesis of vertical descent with differential loss | 46 |
| *Probabilistic expectation of collembolan-affinity genes under vertical inheritance and loss* | *46* |
| *Collembolan percent identity analysis* | *47* |
| Genome localization analysis | 47 |
| Transcriptomic analysis | 48 |
| Sciarid species tree inference | 48 |
| Phylostratigraphic dating | 49 |
| Selective pressure (dN/dS) analysis | 49 |
| Mapping of accessions from previous B. coprophila genome annotation to current annotation | 50 |

| | |
|---|---|
| **SI Tables S1-S30** | |
| Table S1. Previously published B. coprophila HTGs. | 51 |
| Table S2. Per-genome results of DIAMOND screen for HGT across 54 dipteran genomes. (attached file) | 53 |
| Table S3. HGT gene tree analysis. (attached file) | 53 |
| Table S4. Distribution of the numbers of inferred B. coprophila genes per HGT clade. | 54 |
| Table S5. HTG presence across other genomes. (attached file) | 55 |
| Table S6. Per-gene results of DIAMOND screen for HGT across the *B. coprophila* genome. (attached file) | 55 |
| Table S7. Top 20 species by top non-sciarid DIAMOND hits to HTGs. | 56 |
| Table S8. Taxonomic origins of HTGs as determined via manual inspection of gene trees. | 57 |
| Table S9. Data on intron conservation with respect to *Folsomia candida*. (attached file) | 59 |
| Table S10. Non-sciarid recipients of convergent HGT. | 60 |
| Table S11. *B. coprophila* HTGs with literature-derived evidence consistent with transfers of functionally similar HTGs to other eukaryotes. | 61 |
| Table S12. Genomic clustering of HTGs. (attached file) | 63 |
| Table S13. Clusters of adjacent HTGs predicted to catalyze consecutive biochemical reactions. | 64 |
| Table S14. HTG expression data. (attached file) | 65 |
| Table S15. Phylostratigraphic age assignments of B. coprophila HTGs and transfer events. | 66 |
| Table S16. Pairwise dN/dS estimates. (attached file) | 67 |
| Table S17. Ordinary least squares statistics for dN/dS. | 68 |
| Table S18. Functional predictions for B. coprophila HTGs. (attached file) | 69 |
| Table S19. Total numbers of HTGs assigned via manual annotation. | 71 |
| Table S20. Selected HTGs absent or rare in non-sciarid dipterans. | 83 |
| Table S21. Selected dipteran genes and HTGs with predicted function in tryptophan catabolism and de novo $NAD^+$ synthesis. | 85 |
| Table S22. Selected dipteran genes and HTGs with predicted function in biosynthetic pathways. | 86 |
| Table S23. Results of Cox proportional hazards regression for D. melanogaster transgenic experiments. | 88 |
| Table S24. Selected dipteran genes and HTGs with predicted function in repair of damaged DNA bases. | 90 |
| Table S25. RefSeq gene and protein accessions for glyoxylate shunt and β-lactam biosynthesis. | 92 |
| Table S26. Putative HTGs predicted to function in carbohydrate catabolism. | 93 |
| Table S27. Expression of putative glyoxylate shunt HTGs. | 94 |
| Table S28. Genomes used for species tree inference. | 95 |
| Table S29. Dipteran fossils used for soft minimum and maximum constraints in MCMCtree. | 96 |
| Table S30. Parameters used for MCMCtree inference. | 97 |

| | |
|---|---|
| **SI References** | **98** |

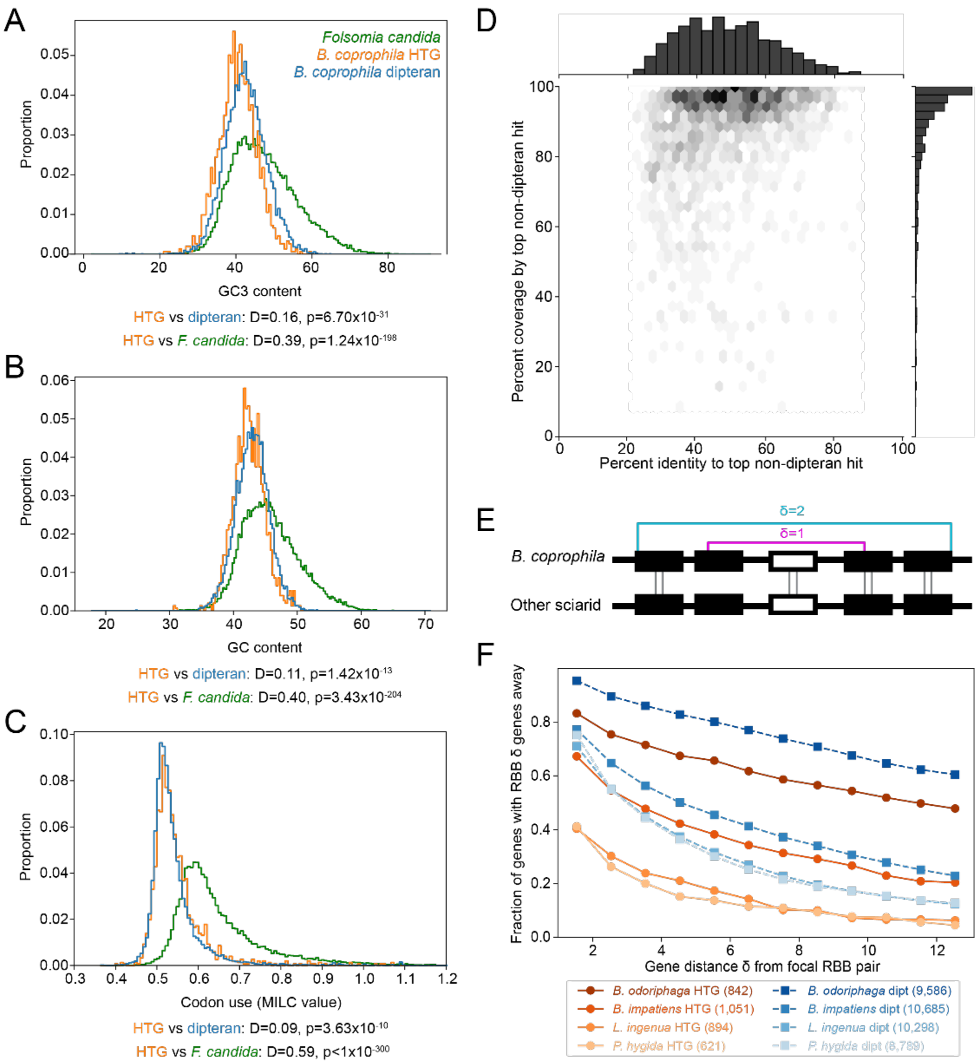

**Figure S1**. **Bioinformatic characteristics of HTGs. (A)** Coding-region GC3 content and **(B)** GC content, respectively, separated by gene origin. **(C)** MILC (Measure Independent of Length and Composition)[1] values for *Bradysia coprophila* coding sequences (CDSs), separated by ancestry. Higher MILC values correspond to greater deviation from global codon use frequencies. Gene ancestry legend in (A) applies also to (B) and (C). In (A)-(C), green curves indicate statistics computed for coding regions across the reference genome annotation of the collembolan *Folsomia candida*, the species containing the most top hits to *B. coprophila*. Two-sample, two-sided Kolmogorov-Smirnov test-statistics (D) and p-values are indicated below each plot. The HTG distributions for all three statistics are more similar to the distributions for dipteran genes than to genes from the top putative donor lineage (lower value of D). **(D)** DIAMOND BLASTp summary statistics for all 1,517 *B. coprophila* putative HTGs. Hexagons show the joint distribution of percent identity and percent query length coverage for the top non-dipteran DIAMOND hit. Histograms along the axes show marginal distributions. **(E)** Schematic illustrating the procedure for microsynteny analysis between genomes of *B. coprophila* and other sciarids. For a given focal reciprocal best BLASTp (RBB) pair (white), we determined whether at least one RBB (connected with gray lines) was present on either side at a distance of $\delta$ genes away, for increasing values of $\delta$. **(F)** The fraction of RBB pairs with at least one RBB pair $\delta$ genes away plotted for varying values of $\delta$, and pairwise combinations of *B. coprophila* and other sciarid genomes. Sciarids in the legend are listed from top to bottom in increasing order of evolutionary divergence from *B. coprophila*, with the total number of dipteran genes and HTGs with RBBs indicated in parentheses. 83.3% (701/842) of *B. coprophila* HTGs with an RBB in *Bradysia odoriphaga* were one gene away from another RBB pair, compared with 95.4% (9,142/9,586) of *B. coprophila* dipteran-ancestry genes.

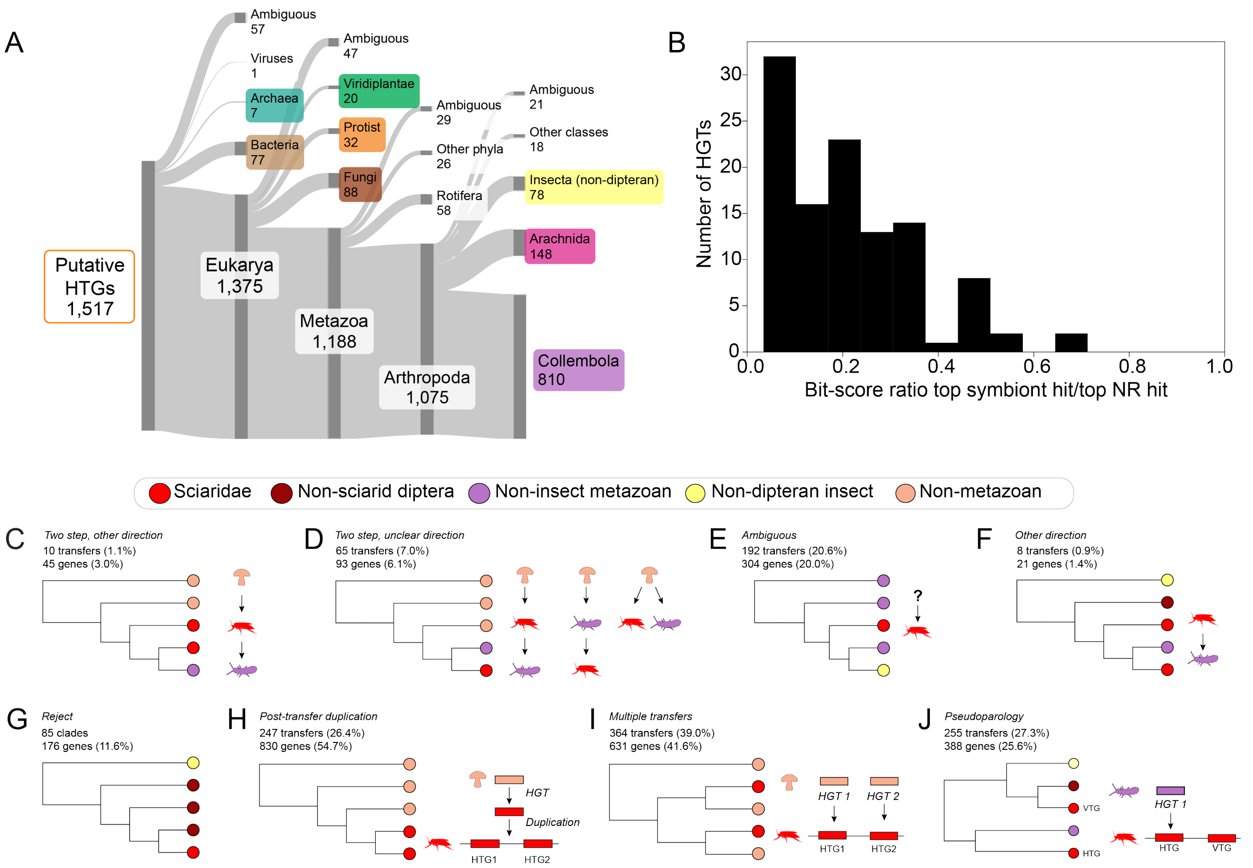

**Figure S2. Supporting data for DIAMOND and phylogenetic inference of HTG origins. (A)** Sankey diagram of the closest non-sciarid sequences ("sisters") to HTGs on midpoint-rooted gene trees. "Ambiguous" indicates that the closest non-sciarid branch of the gene tree contained sequences assigned to multiple taxonomic labels. Branch height is proportional to the number of HTGs. **(B)** The distribution of the ratio of DIAMOND bitscores for top hits in the putative *B. coprophila* rickettsial endosymbiont genome (GenBank accession JAHXDM000000000.1) to top non-sciarid hits in the NCBI NR database for 111/1,517 HTGs with at least one hit in the *B. coprophila* endosymbiont genome annotation. Since no HTGs have a higher bitscore hit in the endosymbiont genome than in their top hit in NR, we conclude that endosymbiotic gene transfer is an unlikely explanation for most putative *B. coprophila* HTGs. **(C-J)** Schematic gene tree topologies supporting the following inferences: **(C)** a two-step other direction transfer, from non-metazoans to Sciaridae, then to non-sciarid metazoans (Table S8G, H); **(D)** two-step transfer of unclear direction (could be non-metazoans to Sciaridae to non-sciarid metazoans, or non-metazoans to non-sciarid metazoans to Sciaridae) or convergent transfers (Table S8D, E); **(E)** an "ambiguous" topology consistent with HGT but of unclear source; **(F)** an "other direction" transfer, from sciarids to non-dipterans, rejected as a transfer to sciarids (Table S8F); **(G)** standard dipteran vertical inheritance, rejected as a transfer to sciarids; **(H)** gene family expansion via post-transfer gene duplication; **(I)** gene family expansion via multiple independent transfers to the *B. coprophila* genome; **(J)** HGT of a gene with a pre-existing dipteran ancestry vertically transferred gene (VTG) homolog (pseudoparalog[2]) in the *B. coprophila* genome. Numbers and percentages of inferred transfer events (of 934) and *B. coprophila* HTGs (of 1,517) are indicated per scenario.

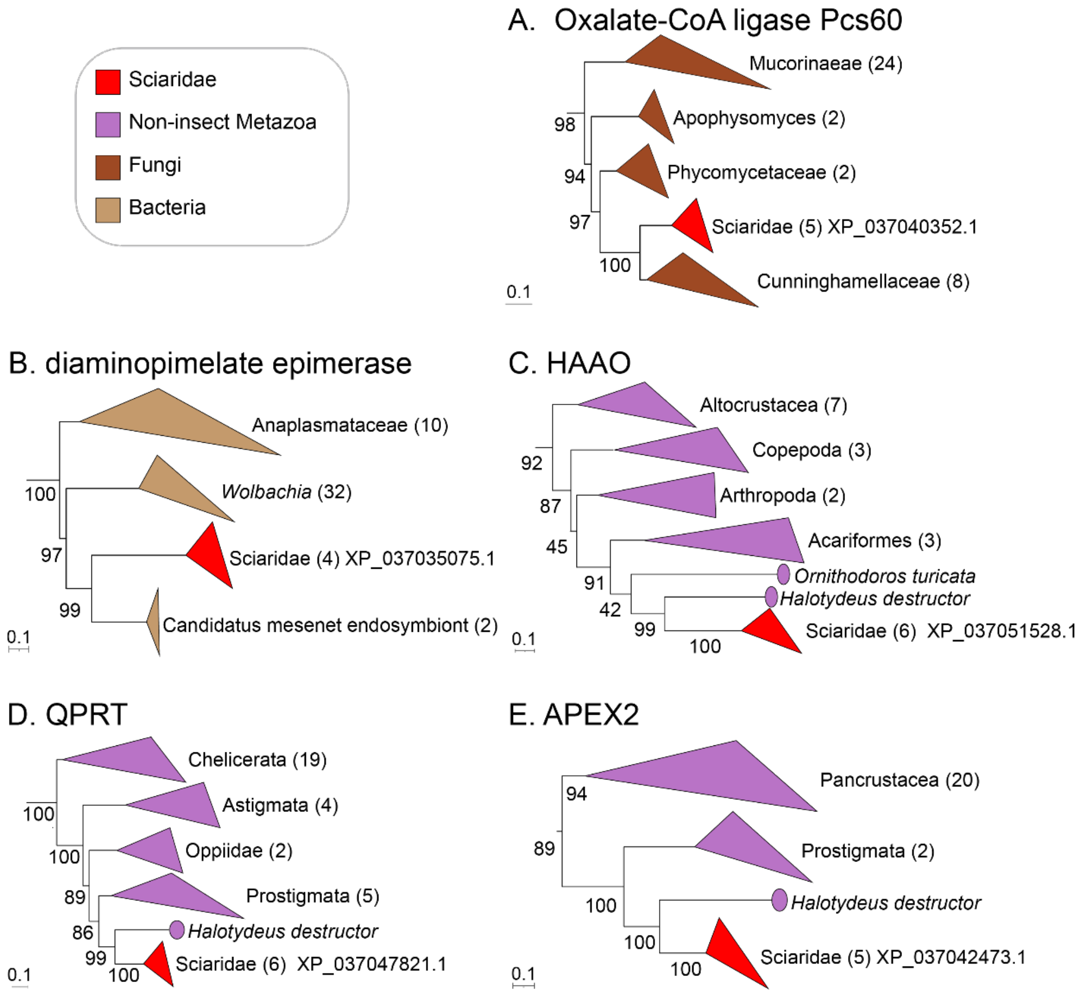

**Figure S3. Examples of gene trees supporting one-step transfers.** Trees are pruned to show only the relevant portion of the tree; full topologies are available on iTOL (https://itol.embl.de/shared/20NWy5j2znqY5). Scale bars show substitutions per site. Internal node numbers indicate ultrafast bootstrap support. Tree clades are labeled with the lowest common taxonomic rank. Circles indicate individual sequences; triangles represent collapsed multi-sequence clades, with edge lengths representing shortest and longest distances from the node to its leaves, and number in parentheses indicating the number of sequences in the clade. Text adjacent to sciarid clades indicates RefSeq accession(s) of *B. coprophila* sequences. **(A)** Tree supporting a one-step transfer from fungi to sciarids (iTOL identifier: tree_454). The sciarid clade contains a single *B. coprophila* sequence we annotate as a predicted Oxalate-CoA ligase Pcs60. The sister clade of the sciarid clade consists entirely of sequences from Cunninghamellaceae (phylum Mucoromycota), a family of predominantly soil-dwelling fungi[3]. **(B)** Tree supporting a one-step transfer from bacteria to sciarids (iTOL identifier: tree_321). The sciarid clade contains a single *B. coprophila* sequence annotated by RefSeq as a diaminopimelate epimerase. The sister clade of the sciarid clade contains multiple members of the endosymbiotic and pathogenic order *Rickettsiales*[4]. **(C)** Tree supporting a one-step HGT event from mites (class Arachnida, superorder Acariformes) to sciarids (iTOL identifier: ML_tree_723). The sciarid clade contains a single *B. coprophila* sequence that we experimentally verified as a 3-hydroxyanthranilate 3,4-dioxygenase (HAAO; Figure 5A, F). **(D)** Tree supporting a one-step HGT event from mites (class Arachnida, superorder Acariformes) to sciarids (iTOL identifier: ML_tree_648). The sciarid clade contains a single *B. coprophila* sequence that we experimentally verified as a quinolinate phosphoribosyltransferase (QPRT; synonym nicotinate-nucleotide pyrophosphorylase; Figure 5A, G). **(E)** Tree supporting a one-step HGT event from mites (class Arachnida, superorder Acariformes) to sciarids (iTOL identifier: ML_tree_520). The sciarid clade contains a single *B. coprophila* sequence we annotated as an apurinic/apyrimidinic-site endonuclease family 2 member (APEX2; synonym putative endonuclease 4), and experimentally analyzed for resistance to alkylating and oxidative stress (Figure 5B, H, K, L).

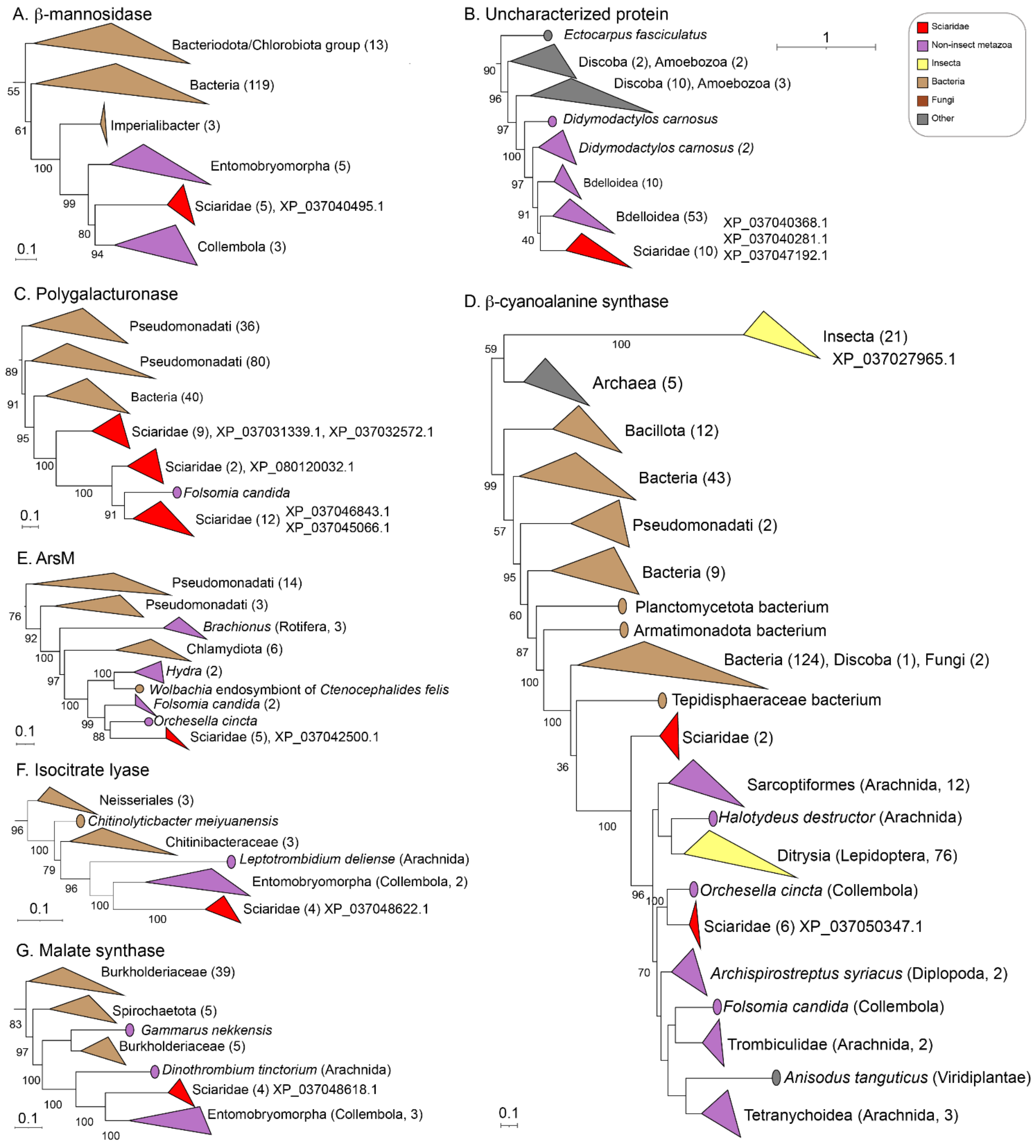
A. β-mannosidase
Bacteriodota/Chlorobiota group (13)
Bacteria (119)
Imperialibacter (3)
Entomobryomorpha (5)
Sciaridae (5), XP_037040495.1
Collembola (3)
55
61
100
99
80
94
0.1
B. Uncharacterized protein
Ectocarpus fasciculatus
Discoba (2), Amoebozoa (2)
Discoba (10), Amoebozoa (3)
Didymodactylos carnosus
Didymodactylos carnosus (2)
Bdelloidea (10)
Bdelloidea (53)
Sciaridae (10)
XP_037040368.1
XP_037040281.1
XP_037047192.1
90
96
97
100
97
91
40
1
Sciaridae
Non-insect metazoa
Insecta
Bacteria
Fungi
Other
C. Polygalacturonase
Pseudomonadati (36)
Pseudomonadati (80)
Bacteria (40)
Sciaridae (9), XP_037031339.1, XP_037032572.1
Sciaridae (2), XP_080120032.1
Folsomia candida
Sciaridae (12)
XP_037046843.1
XP_037045066.1
89
91
95
100
100
91
0.1
D. β-cyanoalanine synthase
Insecta (21)
XP_037027965.1
Archaea (5)
Bacillota (12)
Bacteria (43)
Pseudomonadati (2)
Bacteria (9)
Planctomycetota bacterium
Armatimonadota bacterium
Bacteria (124), Discoba (1), Fungi (2)
Tepidisphaeraceae bacterium
Sciaridae (2)
Sarcoptiformes (Arachnida, 12)
Halotydeus destructor (Arachnida)
Ditrysia (Lepidoptera, 76)
Orchesella cincta (Collembola)
Sciaridae (6) XP_037050347.1
Archispirostreptus syriacus (Diplopoda, 2)
Folsomia candida (Collembola)
Trombiculidae (Arachnida, 2)
Anisodus tanguticus (Viridiplantae)
Tetranychoidea (Arachnida, 3)
59
100
99
57
95
60
87
100
100
36
100
96
100
70
0.1
E. ArsM
Pseudomonadati (14)
Pseudomonadati (3)
Brachionus (Rotifera, 3)
Chlamydiota (6)
Hydra (2)
Wolbachia endosymbiont of Ctenocephalides felis
Folsomia candida (2)
Orchesella cincta
Sciaridae (5), XP_037042500.1
76
92
100
97
100
100
99
88
0.1
F. Isocitrate lyase
Neisseriales (3)
Chitinolyticbacter meiyuanensis
Chitinibacteraceae (3)
Leptotrombidium deliense (Arachnida)
Entomobryomorpha (Collembola, 2)
Sciaridae (4) XP_037048622.1
96
100
79
96
100
100
0.1
G. Malate synthase
Burkholderiaceae (39)
Spirochaetota (5)
Gammarus nekkensis
Burkholderiaceae (5)
Dinothrombium tinctorium (Arachnida)
Sciaridae (4) XP_037048618.1
Entomobryomorpha (Collembola, 3)
83
97
100
100
100
100
0.1

**Figure S4 (previous page). Examples of gene trees supporting two-step transfers.** All except (D) are pruned to show only the relevant portions of the tree topology; full topologies are available on iTOL (https://itol.embl.de/shared/20NWy5j2znqY5). Legend and symbols as in Figure S3. **(A)** Tree supporting a two-step transfer, from bacteria to Collembola, followed by Collembola to Sciaridae (iTOL identifier: ML_tree_464). The sciarid clade contains a single *B. coprophila* sequence that we annotate as a predicted β-mannosidase (Table S18, sheet "carbohydrate_catabolism"). **(B)** Tree supporting a two-step transfer, from protists of the groups Amoebozoa or Discoba to Rotifera, followed by Rotifera to Sciaridae (iTOL identifier: ML_tree_448). The sciarid clade contains three *B. coprophila* sequences annotated by RefSeq as uncharacterized proteins. **(C)** Tree supporting a "two-step, other direction" transfer from bacteria to Sciaridae, followed by Sciaridae to the collembolan *F. candida* (iTOL identifier: ML_tree_197). The three sciarid clades contain predicted polygalacturonase proteins (Table S18, sheet "carbohydrate_catabolism"). **(D)** Tree supporting HGT of an experimentally validated (Figure 5D, O, P) β-cyanoalanine synthase (iTOL identifier: ML_tree_704). Topology is consistent with a two-step transfer, from bacteria to non-sciarid arthropods, then from non-sciarid arthropods to sciarids. The closest non-sciarid arthropod sequence is from the collembolan *Orchesella cincta*, but inference of the sequence and direction of inter-arthropod transfer(s) is complicated by the complex interleaving of collembolan, arachnid, millipede (Diplopoda) and lepidopteran sequences within the arthropod clade. The top insect clade contains insect cystathionine β-synthase enzymes, members of the same enzyme family as β-cyanoalanine synthase[5] that are conserved across eukaryotes but participate in the synthesis of cysteine[6] rather than cyanide detoxification. The tree topology supports the absence of β-cyanoalanine synthase enzymes from non-sciarid dipterans. **(E)** Tree supporting HGT of an experimentally validated (Figure 5C, M, N) arsenite methyltransferase (iTOL identifier: ML_tree_523). Topology is consistent with a two-step transfer, from bacteria to Collembola, then from Collembola to sciarids. Tree also exhibits possible independent transfers to *Hydra* (phylum Cnidaria) and *Brachionus* (phylum Rotifera). **(F)** Tree supporting HGT of a predicted isocitrate lyase (iTOL identifier: ML_tree_661). Topology is consistent with a two-step transfer, from bacteria to non-sciarid arthropods, then from non-sciarid arthropods to sciarids. The closest relatives of the sciarid clade are sequences from the collembolan order Entomobryomorpha. **(G)** Tree supporting HGT of a predicted malate synthase (iTOL identifier: ML_tree_660). Topology is consistent with a two-step transfer, from bacteria to non-sciarid arthropods, then from non-sciarid arthropods to sciarids. The closest relatives to the sciarid clade are sequences from the collembolan order Entomobryomorpha.

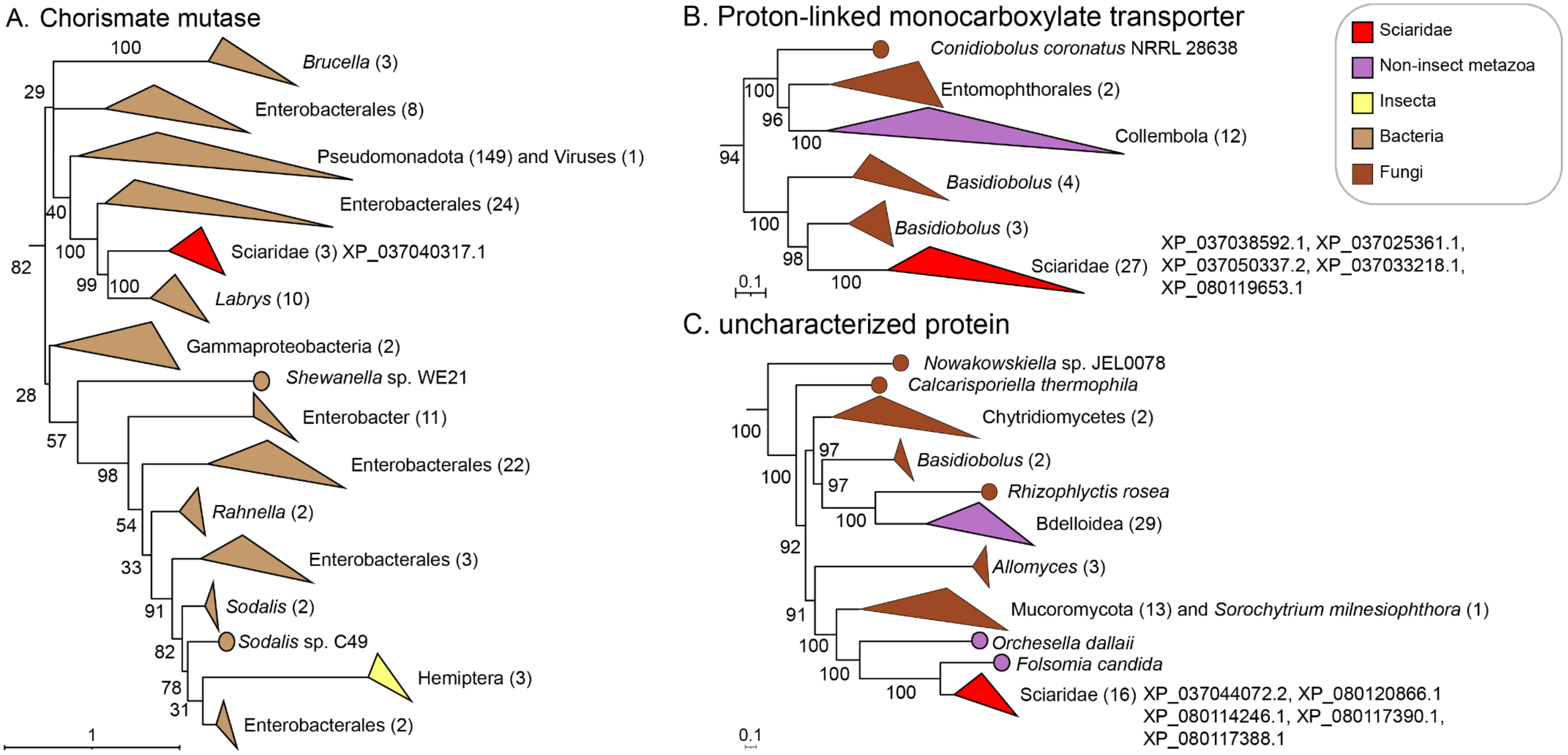

**Figure S5. Examples of gene trees supporting convergent HGT events to sciarids and other metazoan genomes.** Trees are pruned to show only the relevant portions of the tree topology; full topologies are available on iTOL (https://itol.embl.de/shared/20NWy5j2znqY5). Legend and symbols as in Figure S3. **(A)** Tree (iTOL identifier: ML_tree_451) supporting an independent transfer to *B. coprophila* and Hemiptera (class: Insecta) from bacteria. The single *B. coprophila* sequence in the sciarid clade is annotated by RefSeq as a secreted chorismate mutase. **(B)** Tree (iTOL identifier: ML_tree_47) supporting independent HGT events to sciarids and collembolans from fungi. All five *B. coprophila* sequences in the sciarid clade are predicted proton-linked monocarboxylate transporters (Table S18, sheet "manual_annotations"). **(C)** Tree topology (iTOL identifier: ML_tree_562) supporting two independent transfer events from fungi to: (1) bdelloid rotifers and (2) collembolans and sciarids. The sciarid sequences likely originated from a two-step transfer (fungi to collembolans to sciarids), with the sciarid clade containing five *B. coprophila* sequences annotated by RefSeq as "uncharacterized proteins".

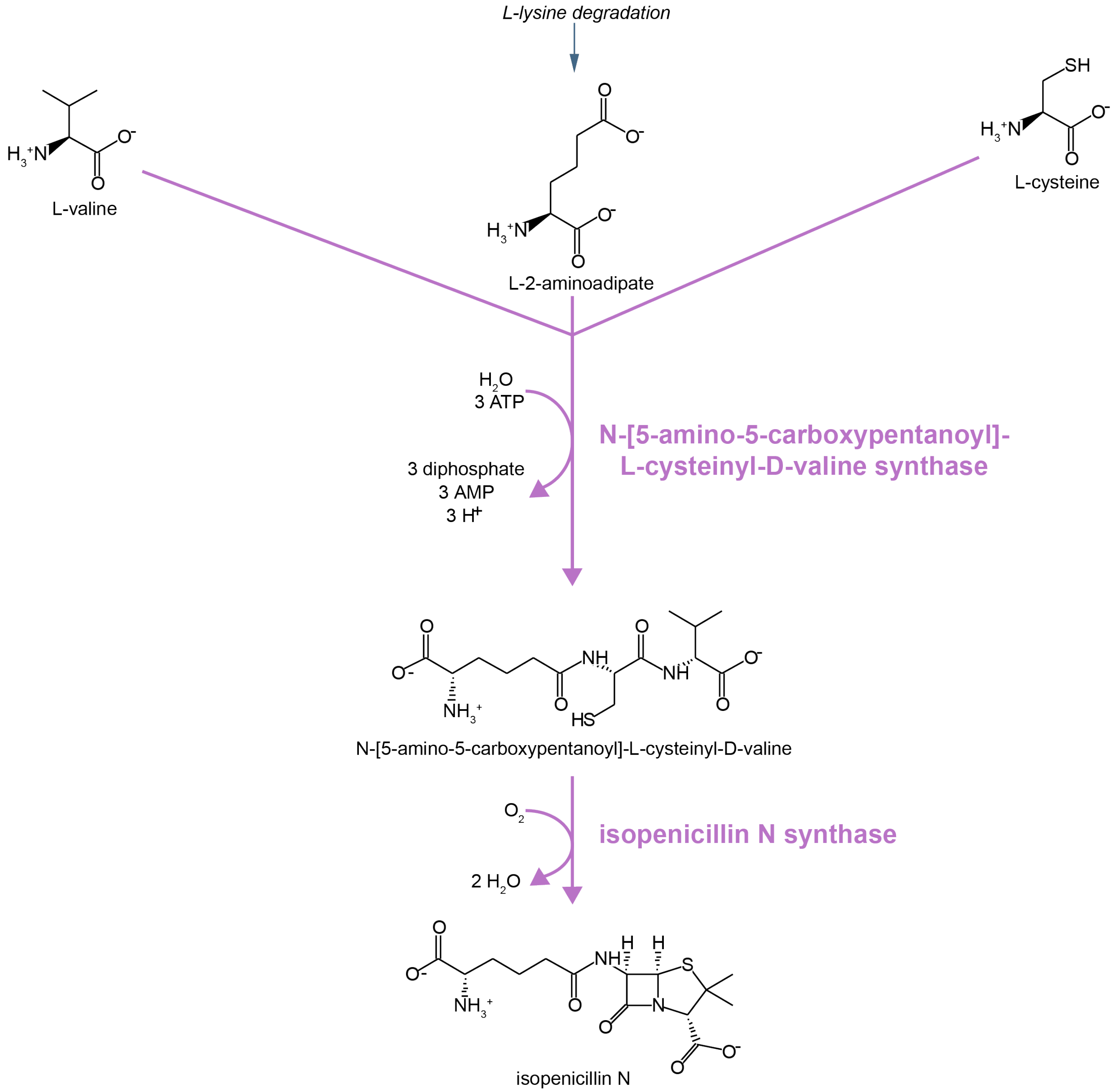

**Figure S6. HTGs may enable the synthesis of β-lactam antibiotics.** Accessions are available in Table S22. Purple arrows and enzyme names indicate reactions predicted to be solely catalyzed or facilitated by HTGs derived from collembolans. Enzymes catalyzing two consecutive steps in synthesis of isopenicillin from amino acid precursors are predicted to be encoded by adjacent HTGs (Figure 3D2). Isopenicillin is the common precursor to all β-lactam antibiotics[28]. Pathway diagram is modified from MetaCyc[24].

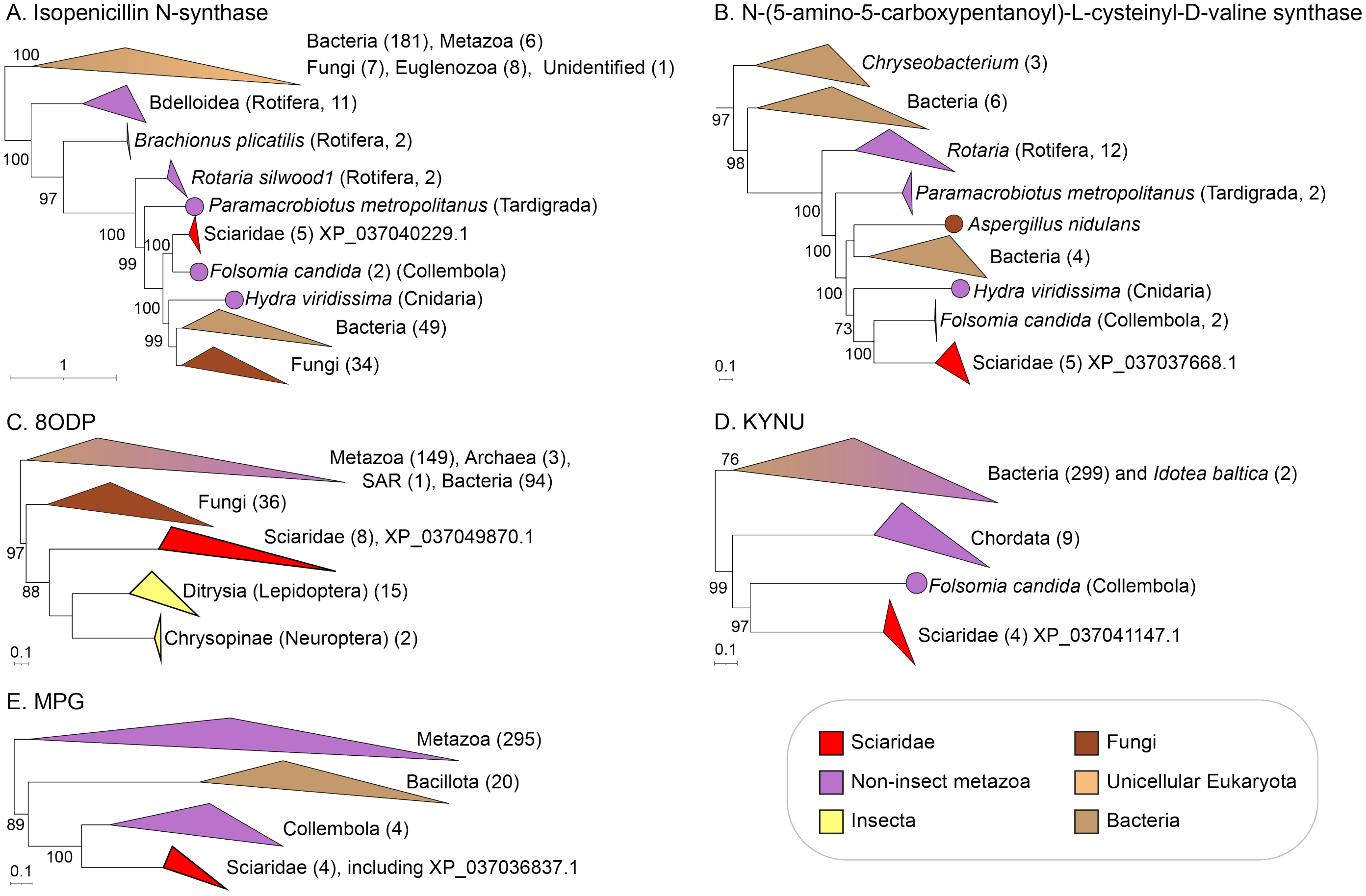

**Figure S7. Examples of gene trees of "ambiguous" topology,** in which the source of the HGT cannot be unambiguously inferred due to a lack of nesting of sciarid sequences in a clade of sequences of a single taxonomic source. **(A)** Tree (iTOL identifier: ML_tree_443) supporting HGT of a predicted *B. coprophila* isopenicillin-N synthase (Table S18, sheet "manual_annotations"). **(B)** Tree supporting HGT of a *B. coprophila* N-(5-amino-5-carboxypentanoyl)-L-cysteinyl-D-valine synthase (iTOL identifier: ML_tree_375), trimmed to show the portion of the tree relevant to HGT inference. In (A)-(B), gene annotations are supported by domain annotation and a sister relationship between the *B. coprophila* sequences and sequences from *F. candida*, in which β-lactam synthesis is biochemically active[7]. Topologies show a complex interleaving of bacterial, fungal, tardigrade, collembolan, and rotifer sequences, which precludes inference of the exact sequence of transfers, although ultimate fungal or bacterial origin appears likely. **(C)** Tree (iTOL identifier: ML_tree_691) supporting HGT of a predicted oxidized purine nucleoside triphosphate hydrolase (8ODP; Figure 5B, I, J). Sciarid sequences are most closely related to sequences from non-dipteran insects (orders Lepidoptera and Neuroptera). **(D)** Tree (iTOL identifier: ML_tree_485) supporting HGT of an experimentally verified *B. coprophila* kynureninase (KYNU; Figure 5A, E). Sciarid sequences are most closely related to a sequence from the collembolan *F. candida*. **(E)** Tree (iTOL identifier: ML_tree_350) supporting HGT of an experimentally verified *B. coprophila* methylpurine glycosylase (MPG; Figure 5B, H). Sciarid sequences are most closely related to sequences from collembolans.

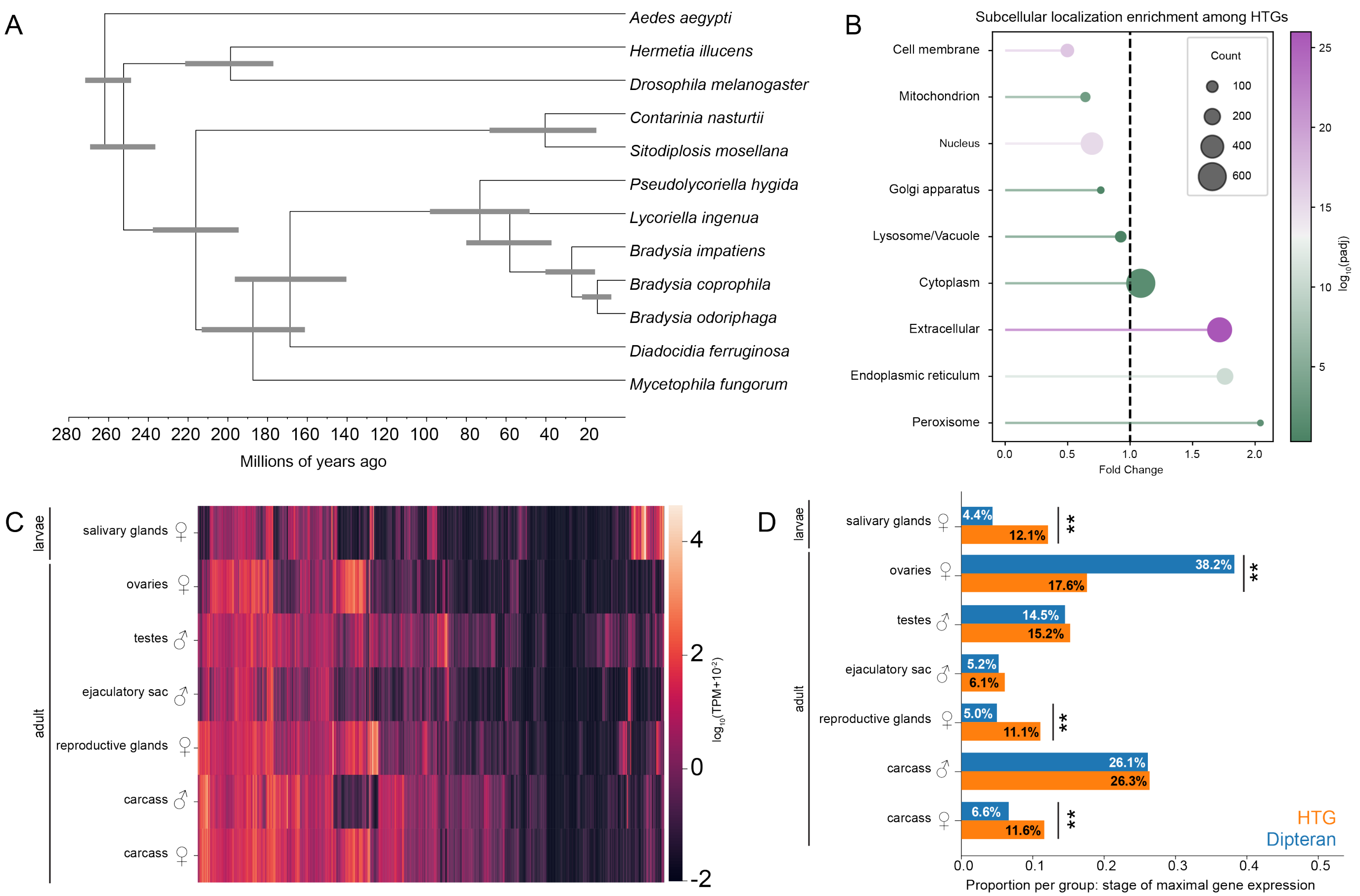

**Figure S8. Additional data related to HTG conservation and predicted functions. (A)** The full time-calibrated species tree that was pruned for Figure 1H. Gray bars around nodes indicate 95% Bayesian credible intervals. *D. ferruginosa* and *M. fungorum* were excluded from Figure 1H because the genomes used for these species were not annotated. Therefore, the gall midges *C. nasturtii* and *S. mosellana* were shown as the closest dipterans with genomes that could be searched via DIAMOND. **(B)** Enrichment of DeepLoc2[8] subcellular localization predictions among HTGs. Circle size indicates the number of HTGs in each gene set; circle color indicates the Benjamini-Hochberg corrected p-value from Fisher's exact test; the position along the x-axis indicates fold-change enrichment over the genome-wide non-HTG background. Full data are available in Table S18, sheet "localization_enrichment". **(C)** Heatmap of mean tissue- or organ-specific expression values of HTGs (horizontal axis) in log transcripts per million from NCBI BioProjects PRJNA1109384 and PRJNA291918. Carcass refers to whole adult bodies without gonads. **(D)** Tissue or organ of maximal mean RNA-Seq expression by gene ancestry. $\chi^2(1)$ contingency test exact p-values from top to bottom are: $6.88 \times 10^{-37}$, $5.42 \times 10^{-55}$, 0.48, 0.19, $1.26 \times 10^{-21}$, 0.87, $3.37 \times 10^{-12}$; $\chi^2$ statistics are: 161.0, 244.0, 0.5, 1.8, 91.3, 0.03, 48.5.

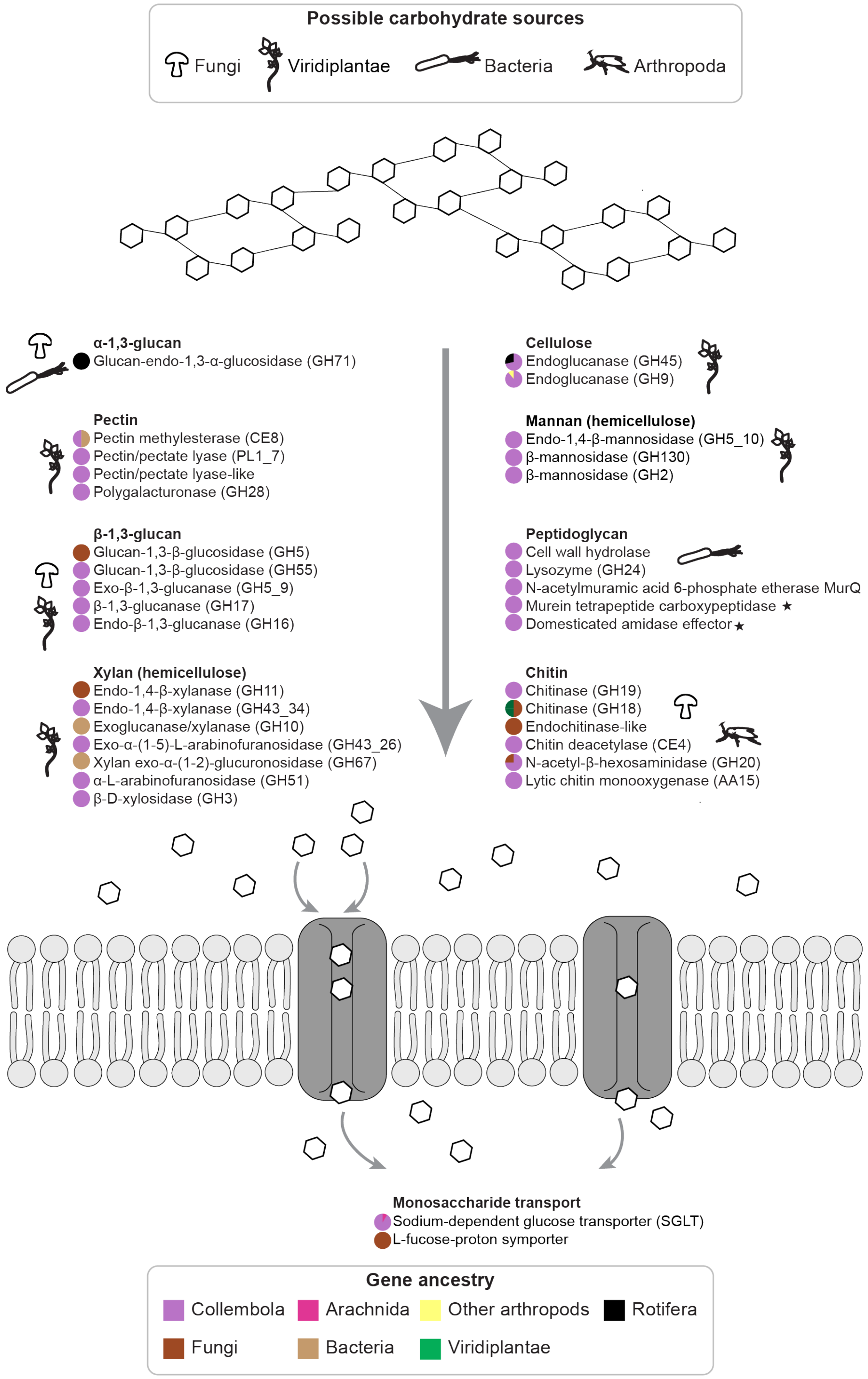

**Figure S9. HTGs may contribute to the catabolism of plant, fungal, and bacterial cell wall-derived polysaccharides.** Pie charts indicate the inferred ancestry of HTGs assigned to each function via manual annotation (Table S18, sheet "carbohydrate_catabolism"). Ancestry for putative HTGs was inferred from the closest non-sciarid phylogenetic relative (sister clade) except when the sister clade was polyphyletic, in which case origins were assigned according to the top non-sciarid DIAMOND hit. Associated subcellular localization predictions and numbers of HTGs in each category are in Table S26. Plausible ecological sources of each carbohydrate[20–23] are indicated with phylopic silhouettes (www.phylopic.org). Stars indicate enzymes predicted to act on non-carbohydrate components of peptidoglycan.

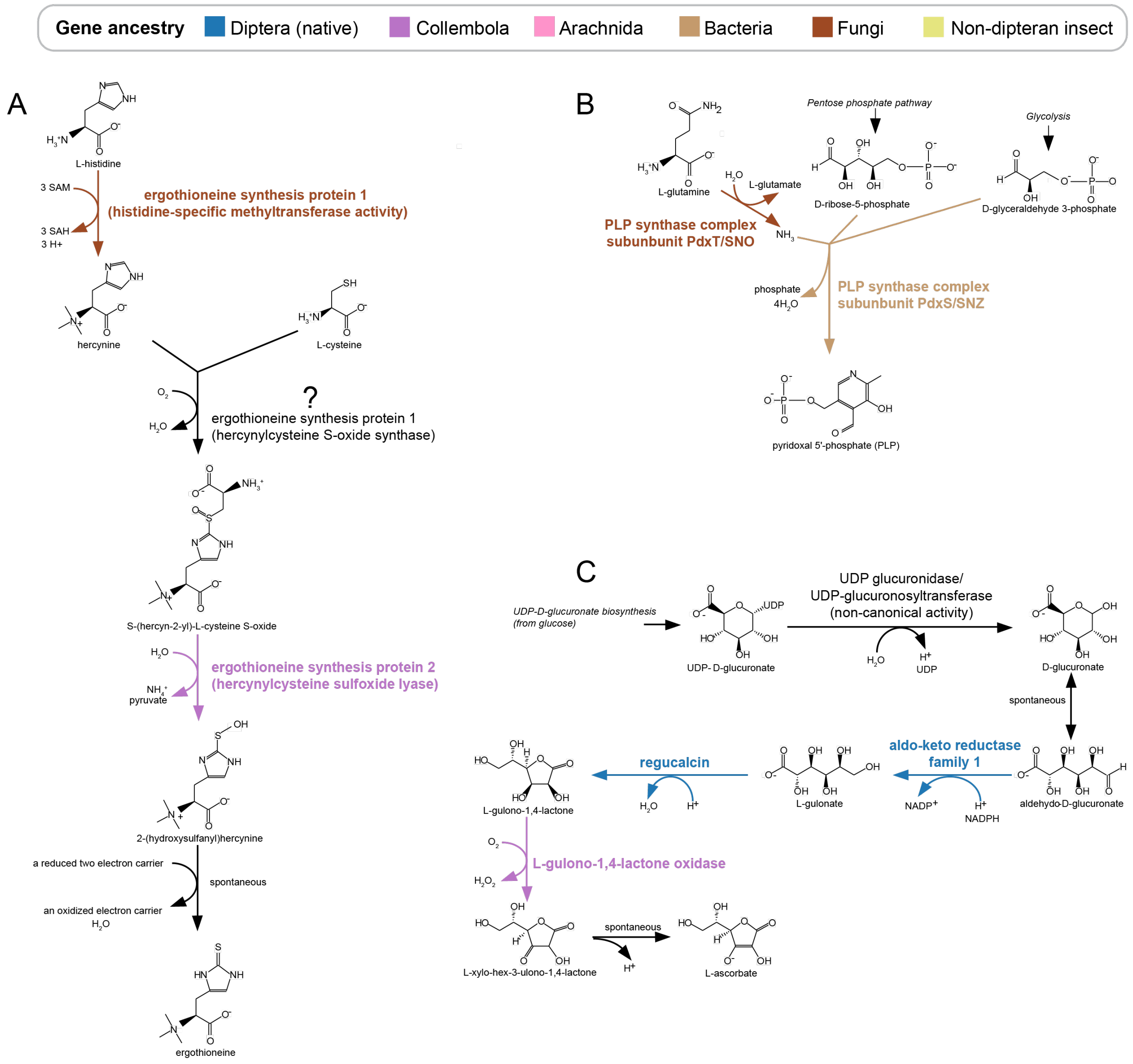

**Figure S10. HTGs may contribute to multi-step antioxidant and cofactor biosynthetic pathways.** Accessions are available in Table S22. Text and arrow colors indicate predicted ancestry, with non-dipteran ancestry shown for reactions predicted to be solely catalyzed or facilitated by HTGs. Pathway diagrams are modified from MetaCyc[24]. "Spontaneous" indicates steps that may occur non-enzymatically. **(A)** Predicted biosynthetic pathway for the antioxidant and non-canonical amino acid ergothioneine. In fungi, the first and second steps are characterized by the same bifunctional protein (ergothioneine synthesis protein 1)[25]. However, the *B. coprophila* protein is lacking the domain predicted to catalyze the second step (indicated by a question mark). SAM and SAH indicate S-adenosylmethionine and S-adenosylhomocysteine, respectively. **(B)** Predicted biosynthetic pathway for coenzyme pyridoxal 5'-phosphate, the biologically active form of vitamin B6. **(C)** Predicted biosynthetic pathway for ascorbic acid (vitamin C). The first step is thought to be catalyzed as a byproduct of UDP-glucuronosyltransferase catalysis[26]. UDP-glucuronosyl and UDP-glucosyl transferase family members (CAZy family GT1) were previously predicted in the *B. coprophila* genome[27], including five HTGs detected here (Table S18, sheet "predicted_CAZymes"). Not shown for simplicity are three collembolan and one arachnid-affinity regucalcin HTGs.

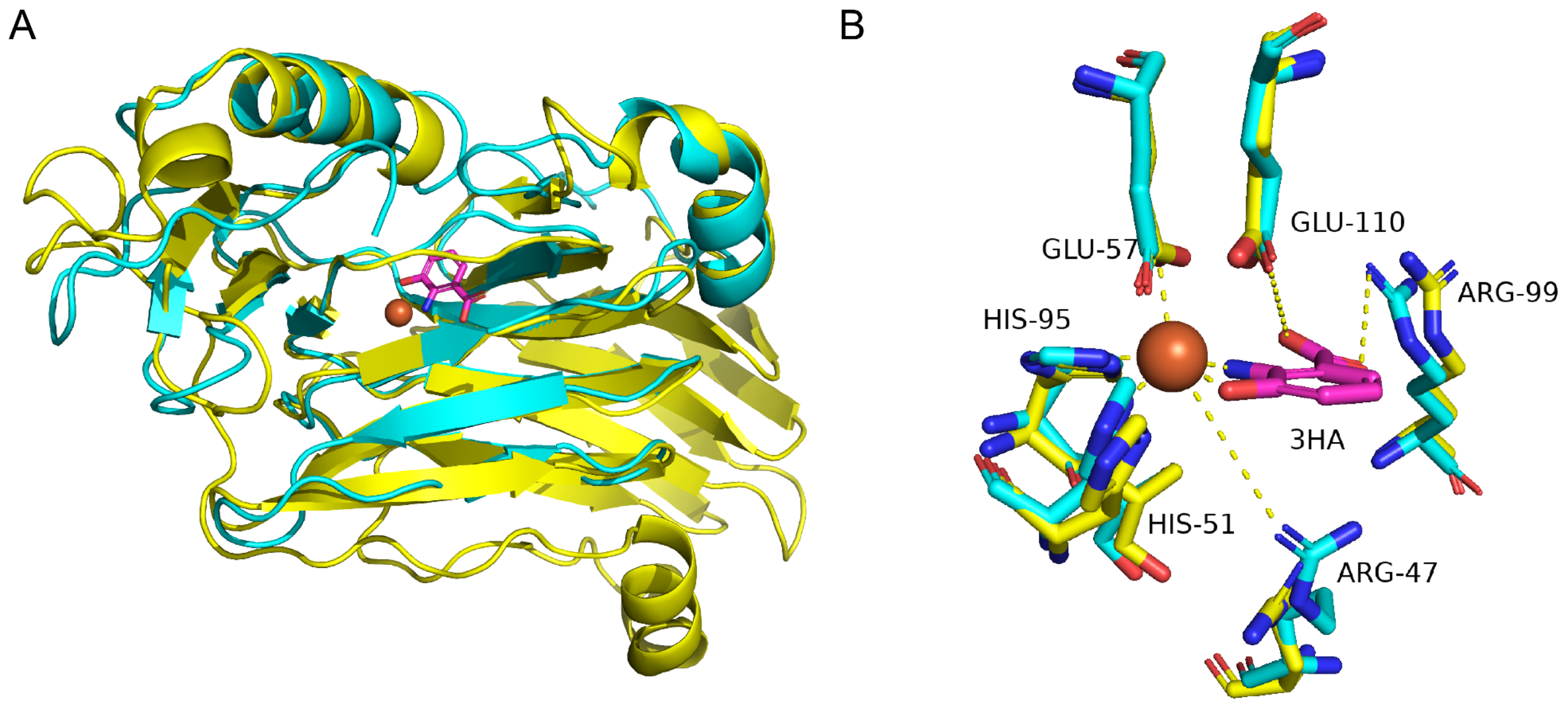

**Figure S11. Alignment of AlphaFold2 predictions for HTGs and solved homolog structures. (A)** Whole-protein alignment (root-mean-square deviation (RMSD) = 0.79) of AlphaFold2 predictions for *B. coprophila* 3-hydroxyanthranilate-3,4-dioxygenase (HAAO; yellow, RefSeq accession XP_037051528.1) to a crystal structure of *Ralstonia metallidurans* 3-hydroxyanthranilate-3,4-dioxygenase (cyan, PDB accession 1YFY) complexed with the Fe(III) cofactor (red sphere) and the 3-hydroxyanthranilate (3HA, magenta) substrate. **(B)** Active site residues from 1YFY (HAAO), with the following functions[9]: coordination of the catalytic Fe(III) ion (His-51, Glu-57, and His-95); stabilization of molecular oxygen as an oxidizing agent (not shown, Arg-47 and Fe(III)); polar contact with the 3HA substrate (Arg-99). All residues shown are conserved in the *B. coprophila* enzyme. Figure 5A, F.

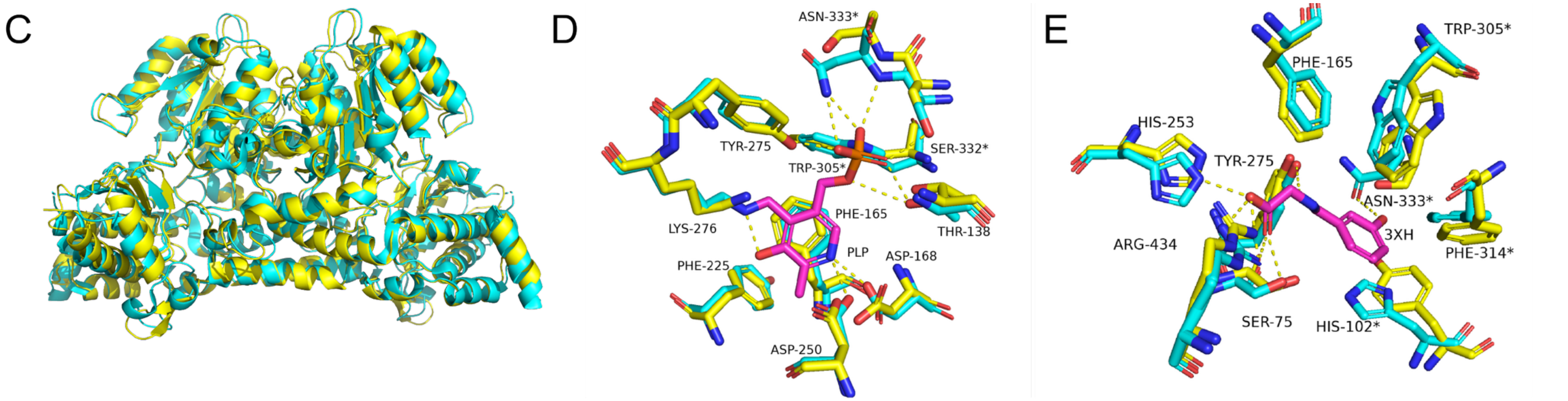

**Figure S11. (C)** Whole-protein dimer alignment (RMSD = 0.93). of AlphaFold2 predictions for a dimer of *B. coprophila* kynureninase (KYNU, color: yellow, RefSeq accession XP_037041147.1) to a crystal structure of *Homo sapiens* kynureninase (cyan, PDB accession 3E9K) complexed with the pyridoxal 5'-phosphate (PLP) coenzyme, and 3-hydroxyhippuric acid (3XH), a structural analog of the substrate 3-hydroxykynurenine. In D-E, asterisks indicate residues in the opposite (symmetric) monomer. **(D)** 3E9K active site residues interacting with PLP (magenta)[10], with the following functions: covalent attachment to PLP (Lys-276); polar contacts to PLP (Tyr-275, Ser-332*, Asn-333*, Asp-168, Asp-250); direct or indirect pi-stacking with PLP (Phe-165, Phe-225, Trp-305*). **(E)** 3E9K active site residues interacting with 3XH (magenta)[10], with the following functions: pi-stacking with the substrate (Phe-165, Trp-305*, and Phe-314*, all universally conserved among kynureninases); polar contacts with the substrate (Arg-434, Tyr-275, Asn-333*, and Ser-75); proposed catalytic residues (found across both **(D)** and **(E)**, Asn-333*, Tyr-275, Arg-434, Lys-276). All residues are conserved in the *B. coprophila* enzyme, except for His-102* and Asn-333*, which have been replaced by physicochemically similar residues (Tyr and Ser respectively). Figure 5A, E.

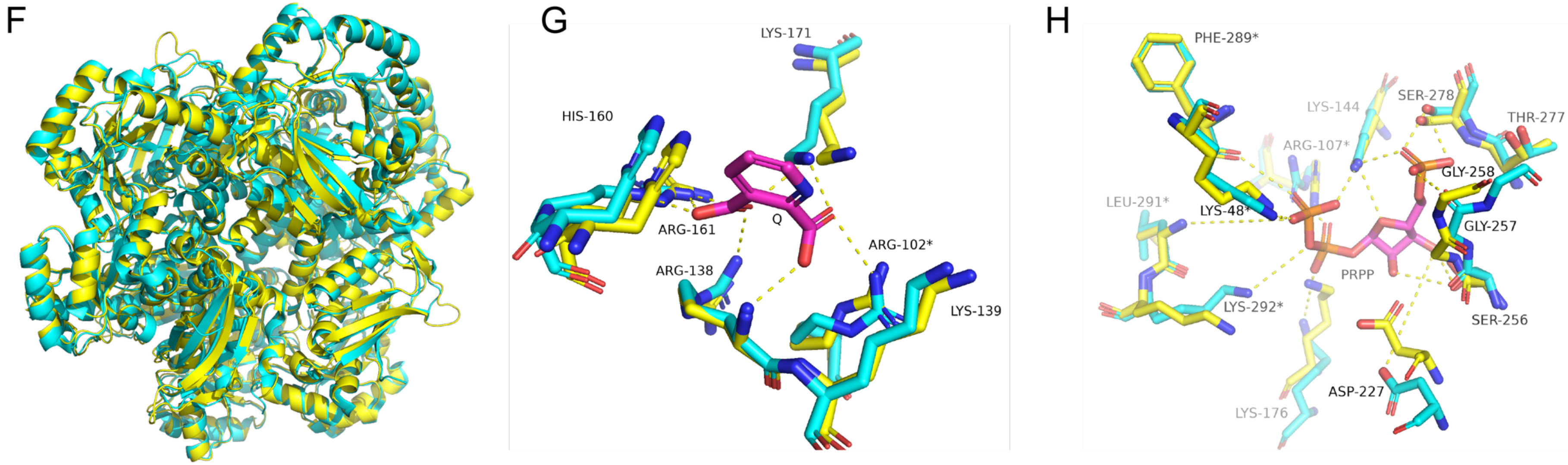

**Figure S11. (F)** Alignment of AlphaFold2 predictions for a *B. coprophila* quinolinate phosphoribosyltransferase (QPRT; yellow, RefSeq accession XP_037047821.1) hexamer aligned to a crystal structure of hexameric *H. sapiens* QPRT (cyan, PDB accession 5AYY, RMSD = 0.85) complexed with quinolinate (Q, magenta). **(G)** Alignment to quinolinate-interacting active site residues from two adjacent monomeric subunits in 5AYY (numbered). All quinolinate-interacting residues are conserved, including residues with the following functions[11,12]: stabilization of the SN1 oxocarbenium intermediate (Arg-138); catalysis (Arg-138* and Lys-171); quinolinate recognition (Arg-102* and Arg-161). Polar contacts to quinolinate are shown as dashed lines. **(H)** Alignment to phosphoribosyl pyrophosphate (PRPP)-interacting active site residues from two adjacent subunits in *S. cerevisiae* quinolinate phosphoribosyltransferase (PDB 3C2F, RMSD = 0.79 for *S. cerevisiae* monomer to *B. coprophila* hexamer). Numbers refer to 3C2F residues. Key conserved residues that interact with the substrate phosphate groups and ribose sugar[13] are shown, along with the predicted catalytic residues. Lys-144, Lys-48* and Lys-176 engage in water-mediated hydrogen bonds (water molecule not shown). All residues are conserved except for Leu-291*, which is replaced by a methionine that maintains the backbone amide NH in the appropriate orientation for hydrogen bonding. In (G-H), dashed lines indicate polar contacts with substrate; asterisks indicate a residue from an adjacent monomer, distinct from the monomer containing the remaining residues. Figure 5A,G.

I

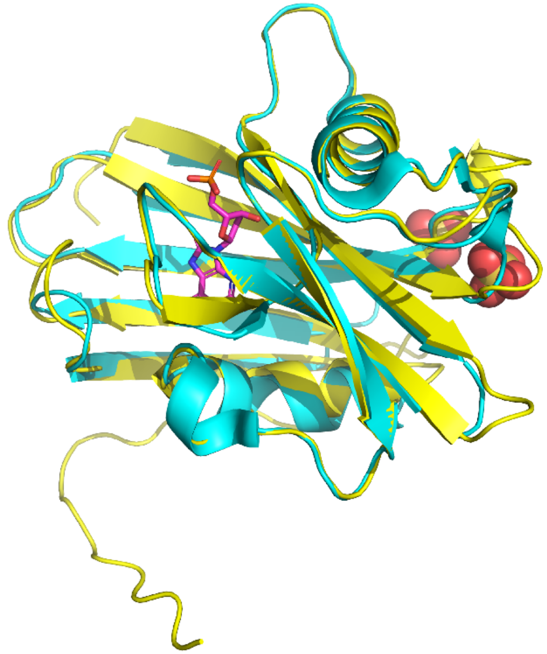

J

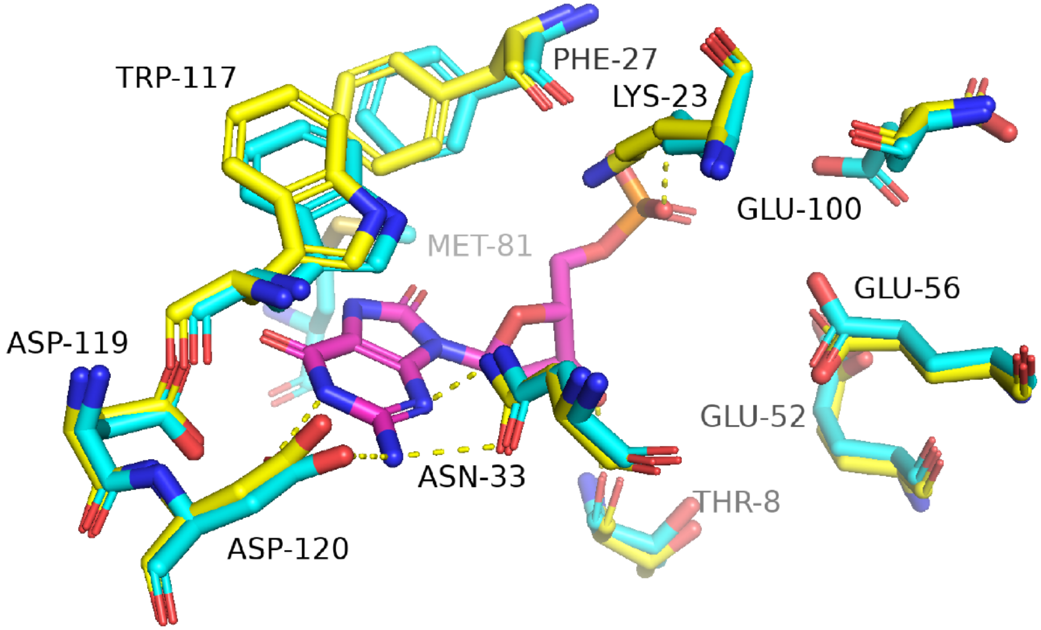

**Figure S11. (I)** Whole-protein alignment (RMSD = 0.46) of AlphaFold2 prediction for *B. coprophila* oxidized purine nucleoside triphosphate hydrolase (8ODP; yellow, RefSeq accession XP_037049870.1) to a crystal structure of its *H. sapiens* homolog, MTH1 (cyan, PDB accession 3ZR0), complexed with sulfate ions (spheres) and its product, 8-oxo-dGMP (magenta). **(J)** 3ZR0 active site residues inferred to participate in the following roles[14] are numbered: substrate hydrolysis (Glu-100, Glu-56, Glu-52); base recognition (Asn-33, Asp-119, Asp-120); non-polar interactions with the product base (Trp-117, Phe-27); substrate polar contacts and binding specificity (Asn-33, Asp-119, Asp-120, Thr-8, Lys-23); recognition of oxidized nucleotides (Met-81 and Phe-27). Lys-23 may instead form a polar contact with an adjacent water molecule (water molecule not shown). All indicated residues except for Met-81 are conserved. Polar contacts with the product are shown as dashed yellow lines. Figure 5B, **I**, J.

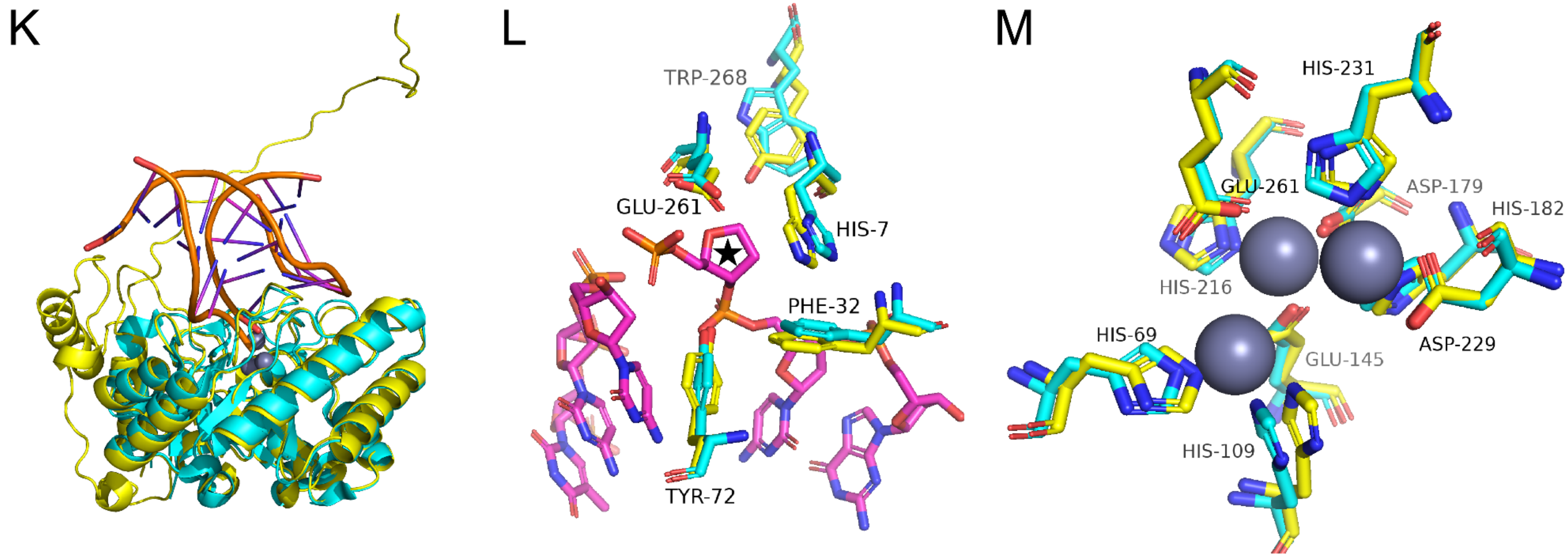

**Figure S11. (K)** Whole-protein alignment (RMSD = 0.73) of AlphaFold2 prediction for *B. coprophila* AP endonuclease family 2 (APEX2; yellow, RefSeq accession XP_037042473.1) to a crystal structure of a member of the same homologous family in *Escherichia coli*, endonuclease IV (cyan, PDB accession 1QUM) complexed with catalytic zinc ions (gray spheres) and double-stranded DNA (orange and magenta) containing a synthetic abasic site. We note that *E. coli* endonuclease IV is a member of the AP endonuclease family 2[15]. **(L)** Alignment of active site residues that interact with the abasic site[16]. A single strand of DNA containing tetrahydrofuran (indicated with a black star) as a synthetic abasic site is shown in magenta. All indicated residues are conserved, except for Trp-268, which has been replaced by a tyrosine. **(M)** Alignment of residues that coordinate three catalytic zinc ions in the active site. All indicated residues are conserved. Residues from 1QUM are numbered in (L-M). Figure 5B, H, K, L.

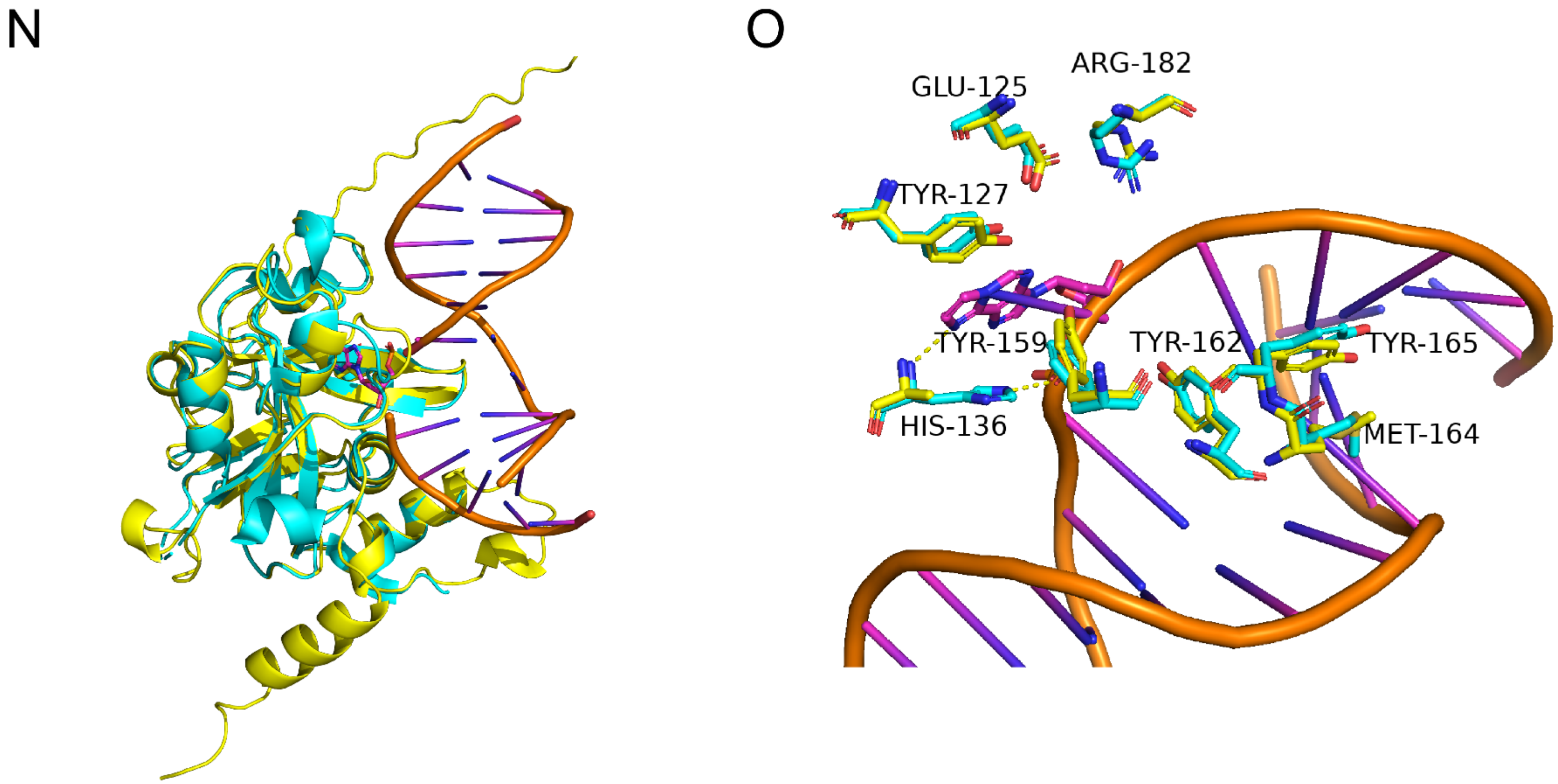

**Figure S11. (N)** Whole-protein alignment (RMSD = 0.57) of AlphaFold2 prediction for *B. coprophila* methylpurine DNA glycosylase (MPG; yellow, RefSeq accession XP_037036837.1) to a crystal structure of its *H. sapiens* homolog, methyladenine DNA glycosylase (cyan, PDB accession 1F4R), complexed with 1,$N^6$-ethenoadenine-DNA, with DNA bases in magenta and backbone in orange. **(O)** Alignment of active site residues identified via site-directed mutagenesis[17]. Polar contacts to the alkylated base are shown as dashed lines. All active site residues are conserved except for His-136, which was found to only partially abrogate catalytic activity when substituted[17]. Tyr-162, Met-164, and Tyr-165 participate in DNA base-pair intercalation; Tyr-127, His-136, and Tyr-159 participate in substrate stacking; Arg-182 and Glu-125 participate in catalytic water binding (water molecule not shown). Figure 5B, H.

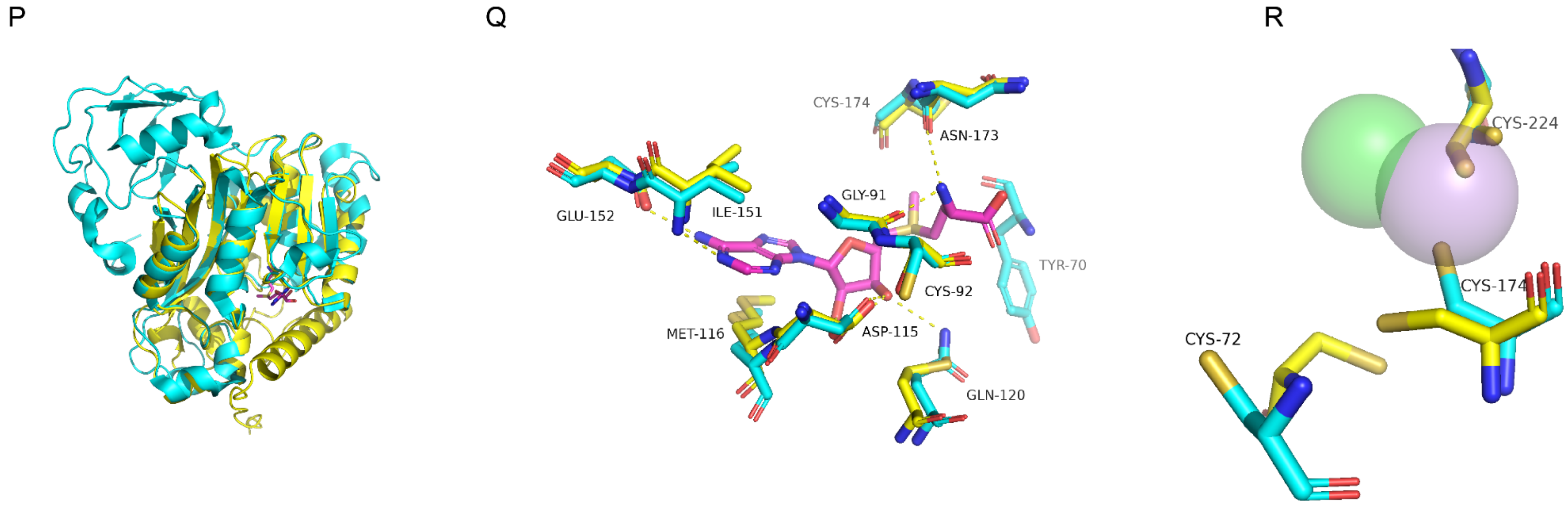

**Figure S11. (P)** Whole-protein alignment of AlphaFold2 prediction for *B. coprophila* arsenite methyltransferase (ArsM; yellow, RefSeq accession XP_037042500.1) to a crystal structure of *Cyanidioschyzon merolae* arsenite methyltransferase (cyan, PDB accession 4FR0; RMSD = 0.70) complexed with an S-adenosylmethionine (SAM) coenzyme or As(III) ion[18]. 4FR0 is missing residues 1 through 48 at the N-terminus (corresponding to the unaligned unstructured region and alpha-helix in yellow only, bottom right). *B. coprophila* arsenite methyltransferase contains the conserved catalytic core, but is missing the C-terminal domain of unknown function (unaligned cyan region, top left) found in its *C. merolae* homolog. **(Q)** Alignment of SAM-binding residues from PDB accession 4FR0[18] to *B. coprophila* arsenite methyltransferase, with residues from 4FR0 numbered and SAM shown in magenta. All residues are conserved, except Gln-120 (replaced by Met) and Tyr-70. Polar contacts to SAM are shown as dashed lines, including a putative contact to Asn-173 not previously reported to our knowledge. **(R)** Alignment of catalytic and As(III)-binding residues from *C. merolae* ArsM PDB accession 4FSD to *B. coprophila* arsenite methyltransferase, with residues from 4FSD numbered, an As(III) ion in purple, and a chloride ion in green. Labeled cysteine residues are conserved across arsenite methyltransferases, with site-directed mutagenesis of any of the three leading to the loss of catalytic activity[19]. Figure 5C, M, N.

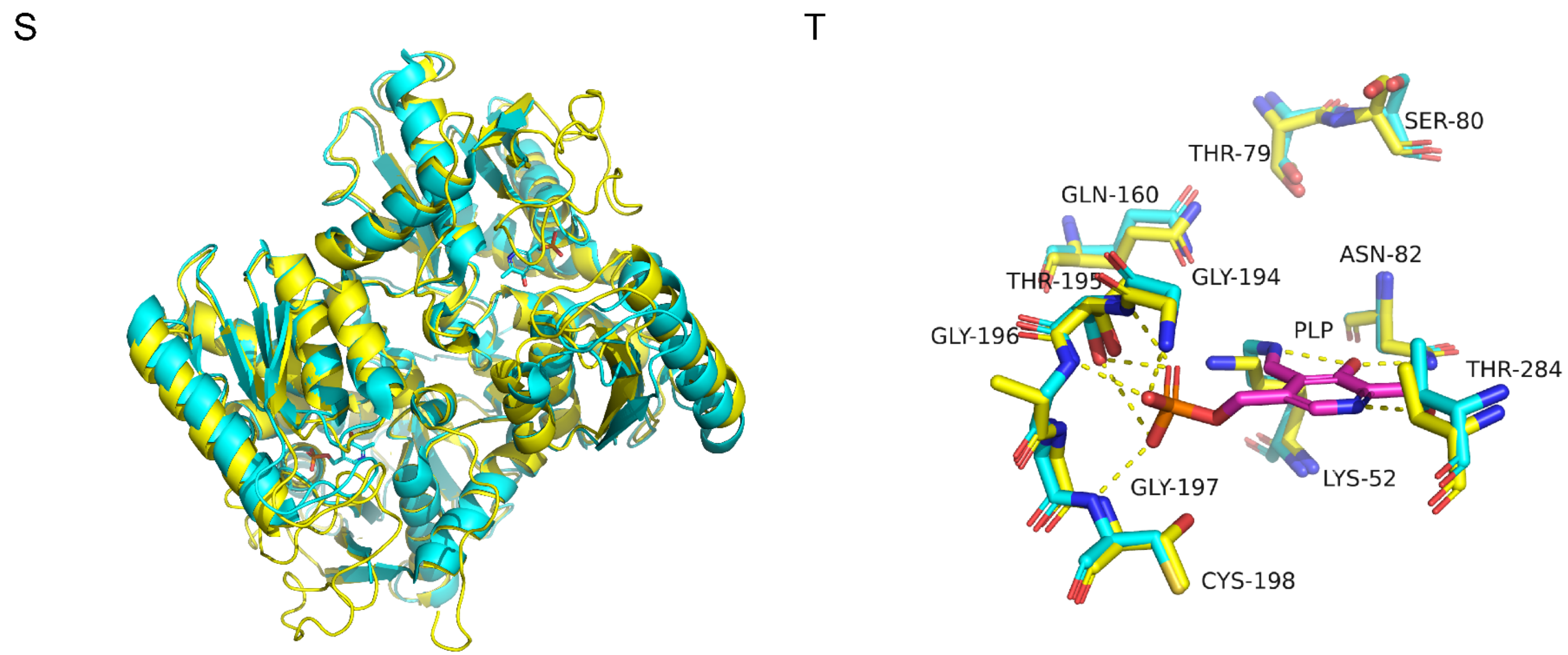

**Figure S11. (S)** Whole-protein dimer alignment (RMSD = 0.93) of *B. coprophila* β-cyanoalanine synthase (β-CAS; yellow, RefSeq accession XP_037050347.1) to a crystal structure of *Tetranychus urticae* β-cyanoalanine synthase (cyan, PDB accession 6PMU) complexed with a pyridoxal 5'-phosphate (PLP) coenzyme (magenta in T). 6PMU is missing residues 221 through 257 (unstructured loops in yellow only) due to the absence of interpretable electron density at these likely unstructured residues[5]. **(T)** Active site residues[5] from 6PMU are numbered and polar contacts to PLP shown as dashed lines. All indicated active site residues are conserved or replaced by physicochemically similar residues (Gly196Ala), including residues responsible for the covalent (Lys-52) and non-covalent (Asn-82, Thr-284, Gly-194, Cys-198, Thr-195) attachment of PLP, and the catalytic residues Thr-79, Ser-80, Asn-82, Gln-160. Figure 5D, O, P.

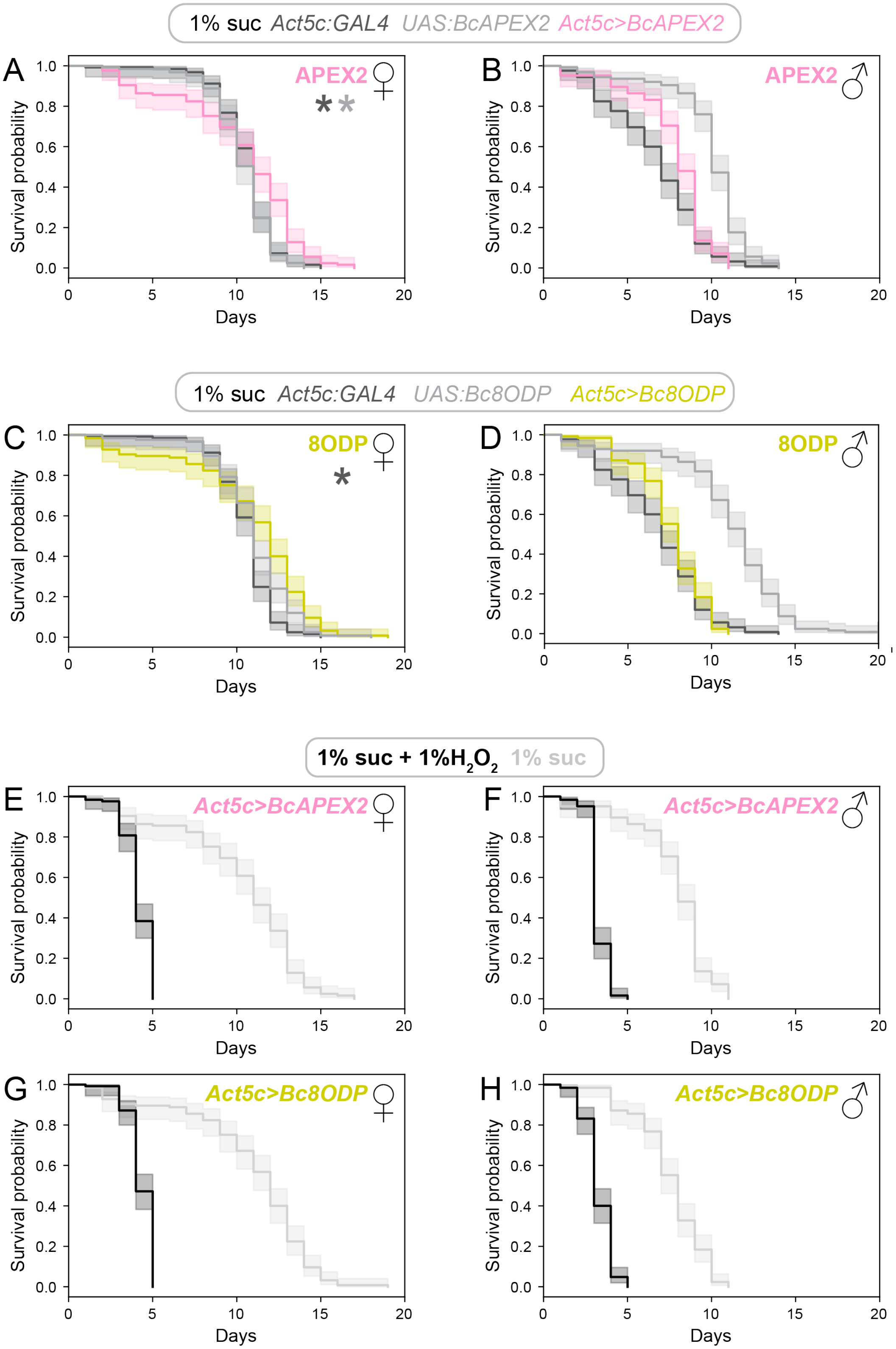

**Figure S12. *D. melanogaster* survival curves related to APEX2 and 8ODP transgenic expression. (A)-(D)** show effects of genotype on survival on a control diet (1% sucrose), for APEX2-expressing and 8ODP-expressing genotypes separated by sex. Asterisks: $p<0.05$ Hazard ratio <1 for the protective effect of the ubiquitous expression compared to control genotypes (indicated by asterisk color) via Cox proportional hazards regression. **(E)-(H)** show the effects of hydrogen peroxide on survival of the genotype expressing the transgene. Shaded regions indicate 95% confidence intervals.

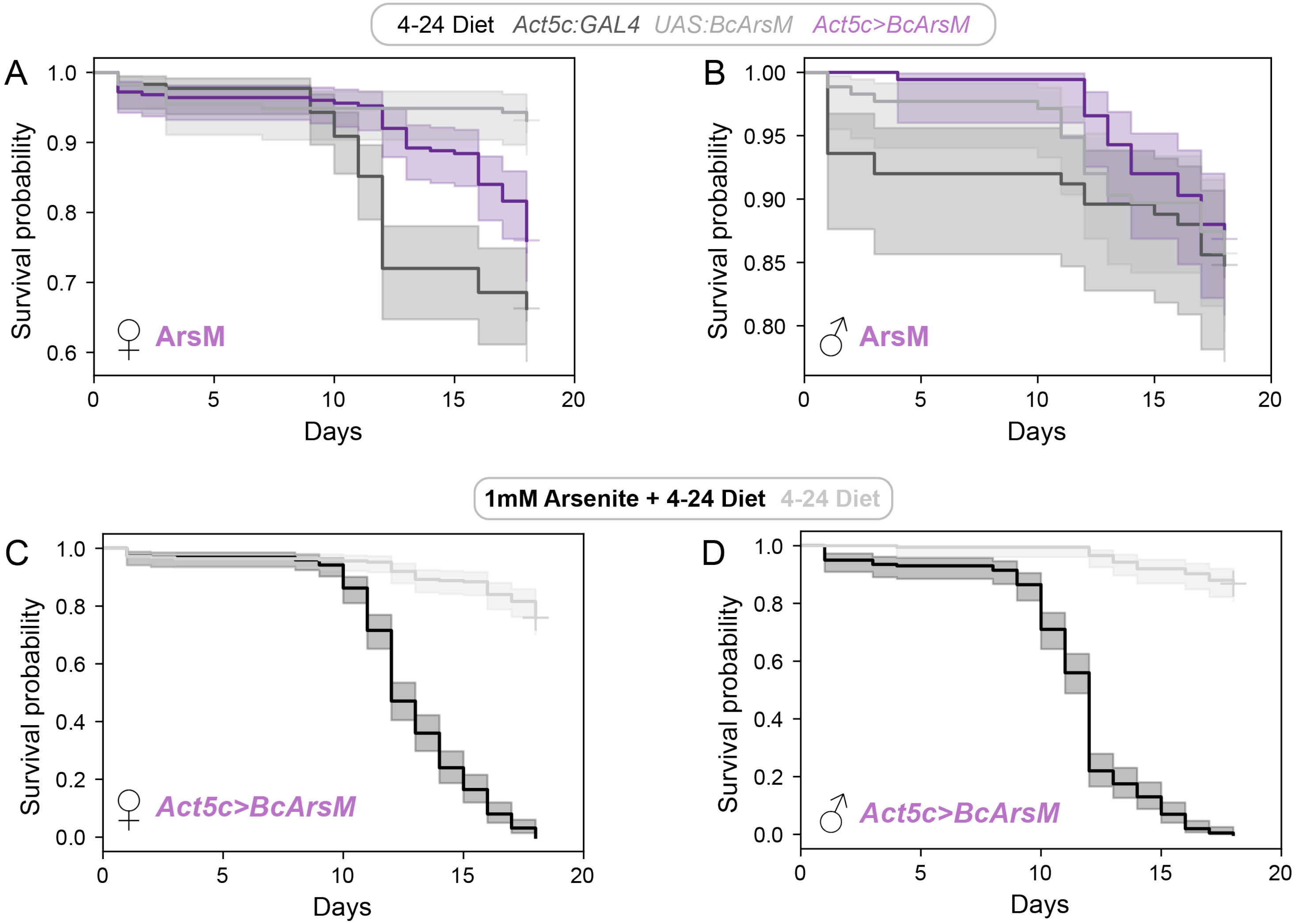

**Figure S13.** ***D. melanogaster*** **survival curves related to ArsM transgenic expression. (A)-(B)** show effects of genotype on survival on a control diet. **(C)-(D)** show the effects of dietary arsenite supplementation on survival of transgenic flies ubiquitously expressing the ArsM transgene. Figure 5C, M, N.

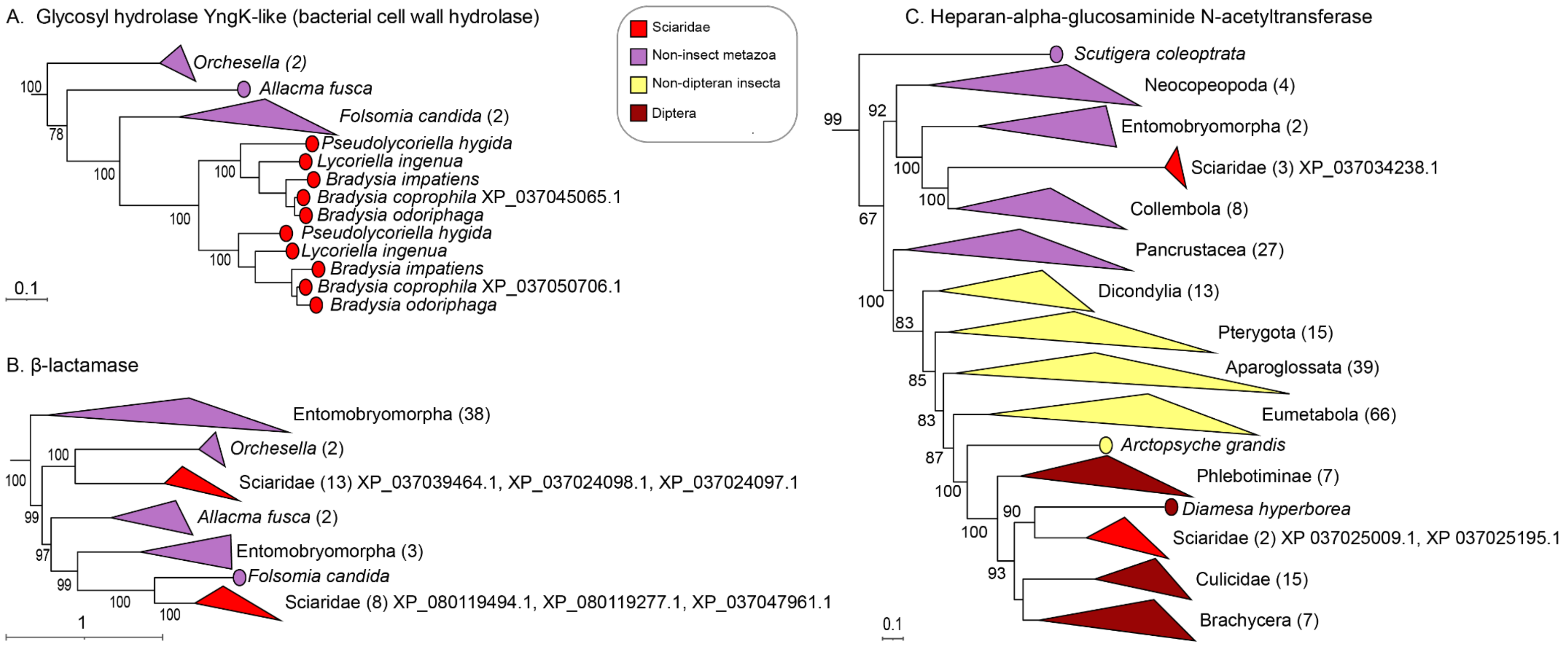

**Figure S14. Examples of gene trees supporting gene family expansion via HGT.** Legend and symbols as in Figure S3. Trees are partial, with full topologies available on iTOL (https://itol.embl.de/shared/20NWy5j2znqY5). **(A)** Tree supporting a one-step transfer from collembolans to sciarids, followed by gene family expansion via post-transfer duplication (iTOL identifier: ML_tree_598). In this case, both the transfer and duplication likely occurred prior to the divergence of *Pseudolycoriella* and *Bradysia*;. The *B. coprophila* genome has two paralogs, both of which we annotate as glycosyl hydrolase YngK-like bacterial cell wall hydrolase proteins (Table S18, sheet "carbohydrate_catabolism"). **(B)** Tree consistent with a gene family expansion via two independent HGT events into sciarid genomes from collembolan donors (iTOL identifier: ML_tree_1). The top and bottom sciarid clades each contain three *B. coprophila* sequences, all of which we annotate as predicted β-lactamases (Table S18, sheet "manual_annotations"). **(C)** Tree supporting a gene family expansion via HGT of a gene with a pre-existing vertically inherited homolog ('pseudoparalog'[2]) in the *B. coprophila* genome (iTOL identifier: ML_tree_291). The top clade of sciarid sequences is of inferred collembolan (HGT) ancestry, and the bottom clade of sciarid sequences is of inferred dipteran (vertical inheritance) ancestry. *B. coprophila* sequences in both clades are annotated by RefSeq as heparan-alpha-glucosaminide N-acetyltransferases.

**SI Video S1 (below & attached video file). Video of *D. melanogaster* cyanide fumigation assay of β-CAS activity.** 20 individuals of each indicated sex and genotype were placed into each vial with 1% agarose containing 10 mM KCN, 50 mM Tris-HCl (pH 7.5) inside a fume hood at room temperature and imaged for 58 minutes. This 30 second video is sped up by approximately 116x. Figure 5O.

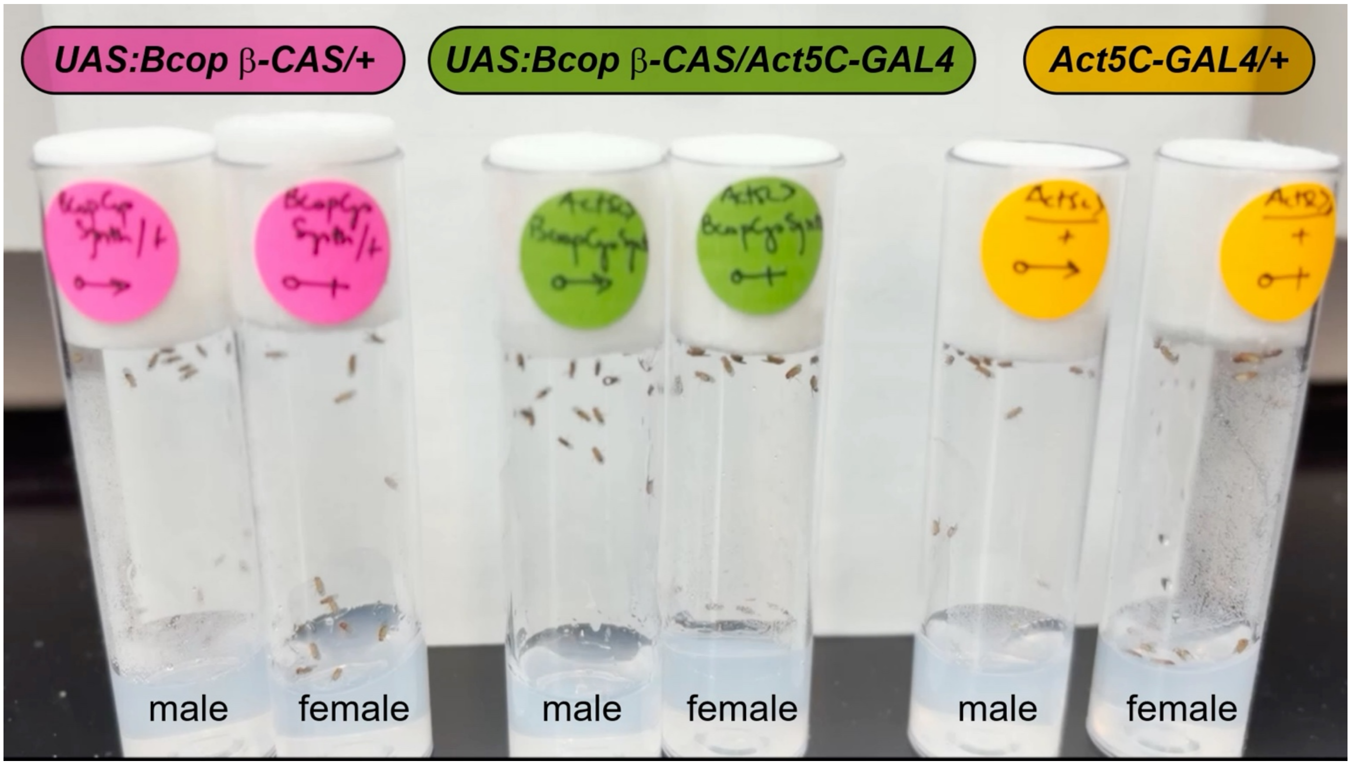

**SI Note 1. HGT contributes to the expansion of gene families.**

Prior studies of horizontal transfer across multiple systems have revealed that HGT frequently contributes to gene family expansions[29–32]. We found that 26.4% (247/934) of sciarid HTG clades contain sequences of more than one *B. coprophila* gene, a topology consistent with the origination of paralogs via gene duplication following a single ancestral HGT event (Figure S2H; Figure S14A; Table S4). We infer that 687 HTGs are single-copy and 830 are members of multigene paralogous families. We further observed that 39.0% (364/934) of sciarid HTG clades co-occur in the same gene tree as a distinct sciarid clade containing sequences from other *B. coprophila* HTG(s) (Figure S2I, Figure S14B). This topology suggests that multiple closely related HTGs might have appeared in the *B. coprophila* genome via independent HGT events, rather than post-transfer duplication. Finally, 27.3% (255/934) of transfer events were inferred in the same gene tree as a dipteran-ancestry homolog (Figure S2J; Figure S14C), suggesting that HGT not only confers novel gene families but may also expand pre-existing sciarid gene families through pseudoparology[2]. Future work should examine whether such cases of gene family expansion induced by the above three HGT-associated mechanisms have resulted in increases in gene dosage, or sub- or neo-functional specialization of the copies, as has been extensively documented for expansions by gene duplication[2,33,34].

**SI Note 2. HTGs have diverse predicted functions in *B. coprophila* stress responses, metabolism, and ecology.**

In the following sections, we describe physiological and biochemical processes that we predict are impacted by HGT in *B. coprophila* based on amino acid sequence-based domain annotations. Following initial analysis of Gene Ontology terms (Figure 4F), we performed detailed manual curation of HTG functions as presented in Table S18 and Figure 6B. We based our inferences on InterProScan[15] domain annotations, with additional supporting data from subcellular localization predictions, published expression data, and published functional experimental studies as available. We hope that this discussion will provide a foundation for future functional investigations into the physiological contributions of HTGs.

*Macromolecule catabolism and digestion*

Gene Ontology analysis revealed that carbohydrate metabolism (GO:0005975) was the most common predicted biological process among HTGs (N = 61 HTGs), and was also strongly enriched among HTGs relative to non-HTGs (fold-change 7.22, FDR-adjusted p-value $1.20 \times 10^{-25}$). To further characterize the contribution of HGT to carbohydrate metabolism, we compared our set of HTGs to a previously published[27] annotation of *B. coprophila* carbohydrate-active enzymes (CAZymes)[27] and found that HTGs constitute 89 of 258 (34.5%) predicted *B. coprophila* CAZymes that mapped to the latest *B. coprophila* genome annotation (GCF_014529535.2; Table S18, sheet "predicted_CAZymes").

Combining these previous predictions[27] with our InterProScan-based manual annotations, we predicted that a total of 126 HTGs participate in carbohydrate catabolism, including 82 predicted to enable the breakdown of plant, fungal, and bacterial cell wall components (Figure S9; Table S18, sheet "carbohydrate_catabolism"). Consistent with a functional role for these enzymes, 12 of 18 *Pseudolycoriella hygida* salivary CAZymes with previous experimental validation of catabolic function[27] are inferred by phylogenetic analysis to be homologous to *B. coprophila* HTGs identified here (Table S18, sheet "confirmed_CAZymes"). Our genome-scale phylogenetic results corroborate and extend prior work showing HGT of two families of sciarid glycoside hydrolases[35,36], and a recent hypothesis based on sequence similarity (but not gene trees) that HGT has contributed to the plant-cell-wall-degrading enzyme repertoire of sciarids[37].

Beyond CAZymes, we predict that an additional 81 HTGs function in protein catabolism, 24 HTGs in amino acid catabolism, and 67 HTGs in lipid catabolism. Since larvae are the predominant feeding stage of the sciarid life cycle[27,38], we examined published data for evidence of expression of catabolism genes in larval samples. Of 600 HTGs with significantly higher RNA-Seq expression in larval than non-larval samples (DESeq2 adjusted p-value < 0.05, $\log_2$ fold change > 0), 49 are predicted to function in protein catabolism (8.2%), 37 in lipid catabolism (6.2%), 80 in carbohydrate catabolism (13.3%), and nine in amino acid catabolism (1.5%). Protein catabolism, lipid catabolism, and carbohydrate catabolism HTGs are significantly overrepresented among larval-upregulated HTGs (Fisher's exact odds ratio values 2.46, 1.94, 2.91, respectively; p-values $7.37 \times 10^{-5}$, $5.74 \times 10^{-3}$, $1.25 \times 10^{-8}$, respectively).

We next examined a published larval saliva proteome of the sciarid *P. hygida* derived via mass spectrometry[27], and found that 11.0% (126/1,149) of genes with expression evidence in this dataset are HTGs via tree-inferred homology to *B. coprophila* HTGs. Of these 126 HTGs detected in the *P. hygida* saliva proteome, 15 (11.9%) are predicted to participate in protein catabolism, 19 (15.1%) in carbohydrate catabolism, six in lipid catabolism (4.8%) and two in amino acid catabolism (1.6%). Protein catabolism and carbohydrate catabolism HTGs are significantly overrepresented among *P. hygida* larval salivary proteome HTGs (Fisher's exact odds ratio values 2.71 and 2.13, respectively; p-values $1.90 \times 10^{-3}$ and $5.79 \times 10^{-3}$, respectively).

Collectively, these data suggest that HGT may have functionally expanded the digestive enzyme repertoire of sciarids, facilitating their exploitation of a broad dietary niche encompassing fungi, leaf litter, living plant tissue, and even other sciarids[37–39]. We note that although endogenous plant cell wall-degrading enzymes are generally rare in animals[40], independent transitions to phytophagy in other lineages of insects have also been accompanied by horizontal acquisition of CAZymes[29,31,41,42].

*Transmembrane transport*

A total of 68 HTGs have domain annotations consistent with function in transmembrane transport, of which 35 (51.5%) have predicted cell membrane localization and 25 (36.8%) have predicted lysosome/vacuole localization. All five HTGs with domain annotations consistent with function in trafficking into or out of the mitochondrion also have predicted mitochondrial localization, including a predicted mitochondrial carnitine/acyl carnitine carrier, which imports lipids into the mitochondrion for catabolism via β-oxidation[43]. Transmembrane transporters include those that may function in nutrient import (e.g., 12 sodium-coupled glucose transporters)[44] as well as toxin export (see below).

*Detoxification/xenobiotic metabolism*

We observed 115 HTGs predicted to contribute to the detoxification, elimination, or tolerance of exogenously derived toxic molecules (xenobiotics). Aside from the cyanide- and arsenite-detoxification HTGs (Figures 5C, D, M-P), we predict that HTGs span all three phases of xenobiotic metabolism[45]. These include 60 putative phase I xenobiotic metabolism enzymes (e.g., 12 flavin monooxygenases and 27 cytochrome P450 enzymes)[45,46], 30 phase II conjugation enzymes (e.g., 17 glutathione S-transferases)[45], and three phase III efflux transporters[45,47]. The *Bradysia odoriphaga* homologs of two HGT-derived *B. coprophila* cytochrome P450 enzymes (published accessions XP_037049686.1[48] with updated accessions XP_080118275.1 and XP_037050080.1[49]) have been shown via RNA interference to promote insecticide resistance[48,49], providing direct experimental evidence of a physiological role for these HTGs. A distinct *B. coprophila* HTG encoding a predicted intradiol ring cleavage dioxygenase is phylogenetically related to an enzyme from the herbivorous spider mite *Tetranychus urticae* (iTOL identifier: ML_tree_724) that has been shown via *in vitro* and *in vivo* assays to metabolize plant toxins and promote survival on exposure to plant toxins[50].

Our results show that HGT of putative xenobiotic metabolism genes extends beyond previous findings of HGT of cytochrome P450s[51] in sciarids. We speculate that horizontally acquired xenobiotic metabolism genes may promote detoxification of naturally occurring defensive molecules produced by plants, fungi, and microbes encountered in the sciarid niche[41,52,53]. Consistent with this interpretation, HTGs with predicted function in xenobiotic metabolism or toxin resistance are enriched among HTGs we detect in a published *P. hygida* saliva proteome (21 HTGs; Fisher's exact test p-value $2.68 \times 10^{-4}$; odds ratio 2.76). Since sciarids secrete saliva into their feeding substrate prior to ingestion[27], we speculate that HTGs could enable environmental detoxification prior to uptake of toxin-bearing food. We note, however, that many enzymes we predict to participate in xenobiotic metabolism may also have unrelated roles in biosynthesis, defense, homeostasis, or other biological processes[45,46].

*Proteostasis*

We predicted that 58 HTGs participate in protein quality control and homeostasis of protein folding, including 11 chaperones and 16 E3 ubiquitin ligases predicted to contribute to protein degradation[21]. We speculate that these HTGs may contribute to protein quality control under homeostatic conditions and/or participate in responses to stresses imposed by environmental toxins, oxidative stress, heat, or cold[54].

*Epigenetic regulation and transposon defense*

A total of 27 HTGs may participate in epigenetic regulation, including four nuclear zinc finger proteins, and a total of ten predicted SET domain-containing histone methyltransferases. Consistent with their domain annotation-predicted functions, 88.9% (24/27) of these putative epigenetic regulators have predicted nuclear localization. Four of these HTGs are homologous to genes that have been shown in other organisms to defend against transposons, including two histone methyltransferases associated with transposon suppression (SUV39H2)[55], and two components of the RNA interference pathway (argonaute 2 and dicer endonuclease)[21,56]. Future work should examine whether these horizontally acquired regulatory genes primarily function in the regulation of other horizontally acquired genes or transposons, or have become integrated into the regulation of endogenous genes.

*Redox homeostasis/oxidative stress responses*

We predict that 67 HTGs contribute to oxidative stress tolerance or redox homeostasis. These include two alternative mitochondrial respiratory proteins (alternative NADH dehydrogenase and alternative oxidase) that may limit the production of reactive oxygen species[57], enzymes that directly detoxify reactive oxygen species (11 catalases and five peroxidases), antioxidant biosynthesis (see below), and antioxidant disulfide-reduction systems (three thioredoxins and 27 glutathione metabolism genes)[58]. An alternative oxidase was previously identified as an HTG and shown to be physiologically active in the sciarid *P. hygida*[39,59]. Oxidoreductase activity is strongly enriched among HTGs (Fisher's exact test odds ratio 5.23, adjusted p-value $5.42 \times 10^{-19}$), raising the possibility that, in addition to responding to possible environmental stressors, oxidative stress response HTGs may facilitate homeostasis in the face of endogenous sources of reactive oxygen species.

*DNA repair*

Domain annotations suggest participation of a total of 16 HTGs in DNA repair processes, including six HTGs in the repair of damaged base pairs (Figure 5B). We also recapitulated the prior finding of a DASH cryptochrome/photolyase, predicted to resolve pyrimidine dimers, as a *B. coprophila* HTG[60,61]. We found that 87.5% (14/16) of HTGs predicted to function in DNA repair are predicted to localize to the nucleus.

*B. coprophila* tolerates double the radiation dose of *D. melanogaster*, a finding previously speculated to be linked to a unique set of DNA repair genes that are transcriptionally upregulated on irradiation[62]. Upon reanalysis of these data, we found that 40 out of 338 (11.8%) genes upregulated upon irradiation are putative HTGs (Table S14, sheet “irradiated_upregulated”). Four of these upregulated HTGs are in the set of HTGs with predicted DNA repair function, including the most highly upregulated gene in this dataset (poly[ADP-ribose] polymerase tankyrase), MPG (validated here via complementation; Figure 5H), Argonaute-2, and a predicted repair-associated polymerase. Irradiation-upregulated HTGs also include seven HTGs predicted to function in proteostasis and three predicted glutathione S-transferases, which participate in the oxidative stress response and xenobiotic metabolism[45]. These findings are consistent with physiological overlap in the effects and responses to DNA damage, oxidative stress, toxins, and protein-folding stress[45,63–65]. Collectively, these HTGs may contribute to *B. coprophila*’s unique capacity to withstand stressors imposed by hypoxia[66] and ionizing radiation[62]. In the following section, we discuss HTGs that may contribute to stress tolerance through biosynthetic pathways.

*Biosynthesis*

We find 13 HTGs that may participate in amino acid biosynthesis, and 92 HTGs that may participate in lipid biosynthesis or modification. The latter category includes six isoprenoid/terpene synthase enzymes, of which four were identified in a recent study that also obtained *in vitro* evidence of function for three of these enzymes[67]. We additionally predict a total of 56 HTGs that may participate in the synthesis of antimicrobials or other secondary metabolites, including the two β-lactam antibiotic biosynthesis genes (Figures 3D2, S6) and 12 genes associated with polyketide or non-ribosomal peptide biosynthesis. *B. coprophila* isopenicillin-N synthase, predicted to participate in β-lactam antibiotic biosynthesis, is phylogenetically most closely related (Figure S7A) to a collembolan HTG whose enzymatic activity was previously experimentally verified[7]. Since biosynthesis of these secondary metabolites has primarily been documented in plants, microbes, and fungi to date[68], future work should explore the biochemical contributions of these HTGs through *in vitro* characterization and *in vivo* metabolomics in *B. coprophila*.

HGT may have additionally contributed to the *de novo* biosynthesis of four compounds in *B. coprophila* that are likely to participate in redox homeostasis (Figure 5A and Figure S10; Tables S21 and S22). Key enzymatic steps in all four pathways are absent in most insects and/or animals (Table S20), highlighting the biochemical novelty potentially conferred by these HTGs. Below, we describe the HTGs that we predict or experimentally demonstrated to participate in these biosynthetic pathways, along with relevant evidence of the antioxidant properties of their end products from existing studies.

- **$NAD^+$.** We provide genetic complementation evidence of the biochemical activity of three consecutive HTGs in the *de novo* synthesis of $NAD^+$ from tryptophan (Figure 5A, E-G). Additionally, we detect two putative horizontal acquisitions of 2-amino-3-carboxymuconate-6-semialdehyde decarboxylase (ACMSD), an enzyme that regulates concentrations of the toxic kynurenine pathway intermediate quinolinate[69]. NAD(H) is a cofactor for several redox homeostasis enzymes and is non-catalytically consumed by $NAD^+$ ADP-ribosyltransferases (e.g. poly(ADP-ribose) polymerases (PARPs)), key components of redox stress responses and DNA repair[65] represented by nine putative HTGs.
- **Ergothioneine**. We find HTGs predicted to encode both enzymatic steps of the fungal ergothioneine biosynthesis pathway (Figure S10A), although we note that the first enzyme catalyzes the first and second steps in fungi[25] but is missing a domain that catalyzes the second enzymatic step in *B. coprophila*. Ergothioneine is a redox-scavenging non-canonical amino acid[25], shown via dietary supplementation to promote redox homeostasis in *Drosophila melanogaster*[70].
- **Pyridoxal 5'-phosphate (PLP)**. Most mammals and insects are obligately dependent on dietary vitamin B6 for PLP owing to one or more ancestral gene losses[71]. We predict that both subunits of the heteromeric PLP synthase complex are encoded by adjacent HTGs in *B. coprophila* (Figure S10B). PLP, the active form of vitamin B6, has been shown via depletion studies to protect against oxidative stress[72] and DNA damage[64]. We note that independent horizontal transfers of PLP biosynthetic enzymes have been documented in plant parasitic nematodes, where they are also speculated to function in responses to oxidative stress[73].
- **Ascorbate (vitamin C)**. We predict that HTGs participate in two steps of the synthesis of ascorbate, an essential free radical scavenger and redox homeostasis regulator (Figure S10C)[74]. The terminal enzyme in this pathway (L-gulono-1,4-lactone oxidase) is putatively encoded by 17 *B. coprophila* HTGs but was lost in or before the last common ancestor of insects[75,76]. Other insects likely acquire ascorbate from diet or via alternative biosynthetic pathways[75,76].

Interestingly, proteins with 'pyridoxal phosphate binding' and 'NADP binding' GO molecular function predictions are enriched in the set of HTGs (fold-change: 3.68, adjusted p-value: $6.28 \times 10^{-4}$ and fold-change: 9.61, adjusted p-value: $8.74 \times 10^{-9}$, respectively), suggesting PLP synthesis and kynurenine pathway HTGs may support the function of other *B. coprophila* HTGs.

*Glyoxylate shunt*

The glyoxylate shunt consists of two consecutive steps that bypass the decarboxylation reactions of the citric acid cycle and thereby enable the net conversion of fatty-acid-derived acetyl-CoA to glucose via gluconeogenesis[77,78]. While it is a key feature of plant[77] and microbial[78,79] metabolism under glucose scarcity, the shunt is absent in most metazoans[21,80]. Both enzymes of the glyoxylate shunt, isocitrate lyase and malate synthase, are predicted to be encoded by adjacent HTGs in the *B. coprophila* genome (Figure 3D1).

Both *B. coprophila* glyoxylate shunt proteins have predicted mitochondrial localization, and expression of both genes peaks in non-feeding adults and is lowest in feeding larvae (Table S27). These localization and expression patterns are consistent with a possible function in gluconeogenesis under glucose scarcity[79,81]. Expression of both genes is also correlated across development with expression of key regulatory enzymes of β-oxidation (carnitine palmitoyltransferase 1) and gluconeogenesis (phosphoenolpyruvate carboxykinase 1) (Table S27). The latter two genes exhibit conserved transcriptional upregulation upon starvation across Metazoa[81,82] and would act upstream (β-oxidation) and downstream (gluconeogenesis) of the predicted glyoxylate shunt, respectively[21,77].

The glyoxylate shunt enzymes have been independently acquired via HGT in the nematode *Caenorhabditis elegans*[80], where they have been experimentally demonstrated to contribute not only to starvation tolerance[81], but also to tolerance of desiccation in the non-feeding dauer stage[79], and to tolerance of protein misfolding resulting from oxidative stress[63]. Future studies should investigate whether glyoxylate shunt HTGs are similarly implicated in *B. coprophila* stress and starvation tolerance.

*Cell signaling*

We predicted that 63 HTGs might function in cell signaling, including two receptor tyrosine kinases with predicted cell membrane localization, four serine/threonine kinases with predicted intracellular localization, four protein phosphatases and three predicted caspases. We predict that 14 HTGs function in neuronal signaling, including two predicted extracellular acetylcholinesterases, three acetylcholine receptor subunits (two with predicted extracellular localization, one with cell membrane localization), an ionotropic receptor (predicted lysosomal/vacuolar localization), and three dopamine β-hydroxylase-like monooxygenases (all with predicted cell membrane localization)[21,83]. HTGs with predicted function in immune signaling are discussed below. Considering that HGT has been more rarely documented for signaling than defense and metabolism genes[84–86], we suggest that these putative signaling genes represent particularly compelling candidates for future experimental analysis.

*Immunological defenses, protein toxins, and host-microbe interaction*

A total of 23 HTGs are predicted to function in canonical metazoan immunity, including nine signaling and pattern recognition genes, and 14 innate immune effector genes, including the antimicrobial peptide defensin (two HTGs)[87], the antiviral effector viperin (one HTG)[88], and two antimicrobial lysozymes[89].

Aside from HTGs canonically associated with metazoan immunity, other horizontally acquired genes not necessarily associated with immune defenses in donor lineages may have been co-opted into roles in host-microbe interaction. We find evidence for HGT of a type VI bacterial cell wall amidase gene, previously reported to have been transferred from bacteria to eukaryotes at least six independent times and functionally co-opted into antibacterial immune defenses in the tick *Ixodes scapularis*[90,91]. A total of 13 HTGs are predicted to function in degradation or recycling of bacterial cell walls, which may reflect analogous co-option into immune defenses or interaction with the *B. coprophila* rickettsial bacterial symbiont[92,93]. Consistent with the latter hypothesis and with reports of the participation of host

HTGs in the maintenance of endosymbiont physiology[85,94], we find that HTGs may participate in biosynthesis of the bacterial cell wall (15 HTGs) and cell membrane (nine HTGs).

HGT may additionally contribute to immune defenses through the biosynthesis of toxins[7,95]. We predict that 12 HTGs encode toxin-like proteins, including three aerolysin-type pore-forming toxins, one hemolysin, and five cytolytic pore-forming toxins. Horizontally acquired pore-forming toxins have been shown in other insects to participate in immunological defenses against parasitoid wasps[95], suggesting a possible defensive role for these *B. coprophila* HTGs. Similarly, 56 HTGs may participate in the biosynthesis of secondary metabolites (e.g., β-lactam antibiotics; Figure S6) that frequently participate in antimicrobial defenses. As was previously proposed for collembolans[7,53], antimicrobial HTGs could function in defense against the high density of potentially pathogenic microbes in the soil niche[96]. Alternatively, antimicrobial HTGs may participate in the control of host-associated microbes such as the putative *B. coprophila* rickettsial endosymbiont. Accordingly, we find the protein target of β-lactam antibiotics (penicillin-binding protein[97]) in our annotation (see Dryad directory PROKKA_06122024 within the repository for this study) of a previously published genome assembly for the *B. coprophila* endosymbiont[93], along with six *B. coprophila* HTGs annotated as putative β-lactamases, which confer resistance to β-lactams[97]. Viewed in the context of emerging literature on the contributions of HGT to the evolution of eukaryotic immune systems and symbioses[90,94,98–100], these results highlight multiple *B. coprophila* HTGs as promising candidates for future functional work on the evolution of immunity and host-microbe interactions.

### Supporting Information Materials: Software and Algorithms

| Name & version | Reference | URL |
|---|---|---|
| BEDTools v.2.31.1 | [101] | https://biocontainers.pro/ |
| Benchling | | https://www.benchling.com/ |
| Biopython v.1.83 | [102] | https://biopython.org/ |
| BLAST+ v.2.12.0 | [103] | https://ftp.ncbi.nlm.nih.gov/blast/executables/blast+/LATEST/ |
| BRAKER3 | [104] | Galaxy BRAKER3 |
| BUSCO Phylogenomics release 20240919 | [105] | https://github.com/jamiemcg/BUSCO_phylogenomics |
| BUSCO v.5.7.1 | [106,107] | https://biocontainers.pro/ |
| CD-HIT v.4.8.1 | [108] | https://biocontainers.pro/ |
| ColabFold v.1.5.5 | [109,110] | https://colab.research.google.com/github/sokrypton/ColabFold/blob/main/AlphaFold2.ipynb |
| DeepLoc2.0 | [8] | https://services.healthtech.dtu.dk/services/DeepLoc-2.0/ |
| DESeq2 v.1.38.3 | [111] | https://biocontainers.pro/ |
| DIAMOND v.2.2.0 | [112] | https://biocontainers.pro/ |
| ETE v.3.1.2 | [113] | https://pypi.org/project/ete3/ |
| FigTree v.1.4.4 | [114] | https://tree.bio.ed.ac.uk/software/figtree/ |
| Gene loss probabilistic recursion | [115] | https://doi.org/10.1093/nar/gkl792 |
| GffRead v.0.12.9 | [116] | https://biocontainers.pro/ |
| InterProScan webserver, release 5.77-108.0 | [15] | https://www.ebi.ac.uk/interpro/search/sequence/ |
| IQ-TREE v.2.2.0.3 | [117] | https://biocontainers.pro/ |
| iTOL webserver and API | [118] | https://github.com/iBiology/iTOL; https://itol.embl.de/ |
| KEGG release 118.0 | [119] | https://www.genome.jp/kegg/pathway.html |
| KofamKOALA release 2026-06-01 | [120] | https://www.genome.jp/tools/kofamkoala/ |
| lifelines v.0.30.0 | [121] | https://zenodo.org/records/14007206 |
| Liftoff v.1.6.3 | [122] | https://biocontainers.pro/ |
| matplotlib v.3.9.2 | [123] | https://matplotlib.org/ |
| MetaCyc release 29.5 | [24] | https://metacyc.org/ |
| Minimap2 v.2.28 | [124] | https://biocontainers.pro/ |
| MUSCLE v.5.1 | [125,126] | https://biocontainers.pro/ |
| NCBI Genome Data Viewer | [127] | https://www.ncbi.nlm.nih.gov/gdv/ |
| NCBI Primer-BLAST | [128] | https://www.ncbi.nlm.nih.gov/tools/primer-blast/ |
| NetworkX v.2.8.8 | [129] | https://github.com/networkx/networkx |
| NumPy v.1.23.5 | [130] | https://numpy.org/ |
| PAL2NAL v.14.1 | [131] | https://biocontainers.pro/ |
| PAML v.4.10.6 | [132–134] | https://biocontainers.pro/ |
| Prokka v.1.14.6 | [135] | https://biocontainers.pro/ |
| pyCirclize v.1.7.1 | [136] | https://github.com/moshi4/pyCirclize |
| PyMOL v.3.0.3 | [137] | https://pymol.org/ |
| Python v.3.9.7 | [138] | https://www.python.org/ |
| R v.4.2.3 | [139] | https://www.R-project.org/ |
| SAMtools v.1.21 | [140] | https://biocontainers.pro/ |

| | | |
|---|---|---|
| scikit-learn v.1.2.0 | [141] | https://github.com/scikit-learn/scikit-learn |
| SciPy v.1.8.1 | [142] | https://scipy.org/ |
| seaborn v.0.13.2 | [143] | https://seaborn.pydata.org/ |
| SRA toolkit v.3.1.1 | [144] | https://biocontainers.pro/ |
| statsmodels v.0.14.0 | [145] | https://www.statsmodels.org/stable/index.html |
| Tracer v.1.7.2 | [146] | https://tree.bio.ed.ac.uk/software/tracer/ |
| Treemble v.1.6.2 | [147] | https://treemble.org/ |
| Trim Galore v.0.6.9 | [148] | https://biocontainers.pro/ |
| trimAl v.1.4.1 | [149] | https://biocontainers.pro/ |
| Twist codon optimization tool | [150] | https://codon-optimization.twistdna.com/ |
| UniProtKB | [151] | https://www.uniprot.org/uniprotkb/ |
| UpSetPlot v.0.9.0 | [152] | https://pypi.org/project/UpSetPlot/ |

### Supporting Information Materials: Experimental Materials

| REAGENT or RESOURCE | SOURCE | IDENTIFIER |
|---|---|---|
| Chemicals, peptides, and recombinant proteins | | |
| Hydrogen Peroxide solution 30%, AR® ACS | Macron Fine Chemicals | Cat# 7722-84-1 |
| Paraquat Dichloride hydrate, PESTANAL | Sigma-Aldrich | Cat# 36541-100MG |
| Potassium cyanide, ACS, 96.0% min | BeanTown Chemical | Cat# 128615-50G |
| Sodium arsenite, 0.1N Standardized Solution | Thermo Scientific Chemicals | Cat# 041533AP |
| Formula 4-24® Instant Drosophila Medium, Plain | Carolina Biological Supply | Cat# 173204 |
| Methyl Methanesulfonate | Sigma-Aldrich | Cat# 129925-5G |
| Yeast Nitrogen base w/o Amino acids and Nicotinic Acid | Formedium | Cat# CYN3901 |
| Experimental models: Organisms/strains | | |
| *B. coprophila* HoLo2 | Yamashita Lab (MIT) | |
| *D. melanogaster*: *actin5c Gal4/CyO:y[1] w[*]; P{w[+mC]=Act5C-GAL4}25FO1/CyO, y[+]* | Bloomington Drosophila Stock Center (BDSC) | RRID:BDSC 4414 |
| *D. melanogaster*: *Oregon-R-C* | BDSC | RRID:BDSC 5 |
| *D. melanogaster*: *yw; UAS Bcop APEX2/CyO* | This study | |
| *D. melanogaster*: *yw; UAS Bcop ArsM/CyO* | This study | |
| D. melanogaster: *yw; UAS Bcop β-CAS/CyO* | This study | |
| *D. melanogaster*: *yw; UAS Bcop 8ODP/CyO* | This study | |
| *S. cerevisiae* BY4741 bna1Δ | Yeast Deletion Collection: Invitrogen | #95401.H2 |
| *S. cerevisiae* BY4741 bna5Δ | Yeast Deletion Collection: Invitrogen | #95401.H2 |
| *S. cerevisiae* BY4741 bna6Δ | Yeast Deletion Collection: Invitrogen | #95401.H2 |
| *S. cerevisiae* BY4741 apn1Δ | Yeast Deletion Collection: Invitrogen | #95401.H2 |
| *S. cerevisiae* BY4741 mag1Δ | Yeast Deletion Collection: Invitrogen | #95401.H2 |
| *S. cerevisiae* BY4741 bna1Δ + pPN352 (Empty Vector) | This study | |
| *S. cerevisiae* BY4741 bna5Δ + pPN352 (Empty Vector) | This study | |
| S. cerevisiae BY4741 bna6Δ + pPN352 (Empty Vector) | This study | |
| *S. cerevisiae* BY4741 apn1Δ + pPN352 (Empty Vector) | This study | |
| *S. cerevisiae* BY4741 mag1Δ + pPN352 (Empty Vector) | This study | |
| *S. cerevisiae* BY4741 bna1Δ + pPN356 | This study | |
| *S. cerevisiae* BY4741 bna5Δ + pPN357 | This study | |
| *S. cerevisiae* BY4741 bna6Δ + pPN358 | This study | |
| *S. cerevisiae* BY4741 apn1Δ + pPN362 | This study | |
| *S. cerevisiae* BY4741 mag1Δ + pPN370 | This study | |
| Recombinant DNA | | |
| Plasmid: pJFRC12-10XUAS-IVS-myr::GFP | [153] | Addgene #26222 |
| Plasmid: UAS: Bcop APEX2 | This study | |
| Plasmid: UAS: Bcop Arsenite methyltransferase | This study | |
| Plasmid: UAS: Bcop β-CAS | This study | |
| Plasmid: UAS: Bcop 8ODP | This study | |
| Plasmid: pPN352 (Episomal expression plasmid with Actin promoter and alcohol dehydrogenase terminator) | This study | |
| Plasmid: pPN356 (pPN352-BcHAAO) | This study | |
| Plasmid: pPN357 (pPN352-BcKYNU) | This study | |
| Plasmid: pPN358 (pPN352-BcQPRT) | This study | |
| Plasmid: pPN362 (pPN352-BcAPEX2-Cterm:2X SV40 NLS) | This study | |

| | | |
|---|---|---|
| Plasmid: pPN370 (pPN352-BcMPG-Cterm:2X SV40 NLS) | This study | |
| Other | | |
| Fly Injection Service | | BestGene Inc. |

### SI Methods

#### Phylogenetic inference

*Sciarid database construction*

We generated a custom DIAMOND database of sciarid proteins consisting of the longest protein isoforms per gene for the publicly available annotations of *Bradysia coprophila* (RefSeq accession GCF_014529535.2), *B. odoriphaga* (GenBank accession GCA_016920775.1), and *Pseudolycoriella hygida* (GCA_029228625.1)[27]. We additionally supplemented this database with protein sequences we generated via BRAKER3[104] annotation of the unannotated assemblies of *Lycoriella ingenua* (GCA_965233655.1) and *B. impatiens* (GCA_965233325.1)[154]. Soft-masked genomic FASTA files were annotated on the Galaxy webserver[155] (Galaxy BRAKER3) with a protein evidence FASTA containing the longest isoform per *B. coprophila* protein-coding gene along with the OrthoDB[156] version 12 protein set for Arthropoda (Bioinformatics Web Server - University of Greifswald). Protein FASTA files were extracted using GffRead[116] (version 0.12.9), and the single longest isoform per protein-coding gene was selected, for a total of 35,264 and 32,751 proteins for *L. ingenua* and *B. impatiens*, respectively. Annotation completeness was assessed with BUSCO[106] (version 5.7.1) against the diptera_odb10 set, with a score of 95.9% for *L. ingenua* and 95.0% for *B. impatiens*. Protein FASTA files for these two species were clustered at 90% identity and 90% length coverage using CD-HIT[108] (version 4.8.1, resulting in 31,776 sequences for *L. ingenua* and 29,074 sequences for *B. impatiens*) and subsequently concatenated to generate a combined sciarid database, with a resulting total of 111,399 sequences.

*Maximum likelihood gene tree construction*

Sequences for maximum likelihood phylogenetic inference were selected from separate DIAMOND BLASTp runs against the clusteredNR database[157] as well as the sciarid database generated above. To reduce redundant phylogenetic inferences, we used a DIAMOND search against the sciarid database above to cluster candidate HTG sequences by similarity. We generated a NetworkX[129] (version 2.8.8) protein similarity network in which HTGs that were DIAMOND hits of other HTGs were connected with an edge if their alignment to other HTGs satisfied all of the following criteria:

1. DIAMOND E-value < $10^{-10}$.
2. DIAMOND bit-score > 0.80*max (bit-score outside of Sciaridae in the clusteredNR database), selecting sequences likely to be more closely related to the focal HTG than non-sciarid sequences.
3. > 30% alignment overlap with the region of the query that aligned with its top non-sciarid hit.

Similarity clusters were defined as weakly connected components on this network. For each of the 924 resulting similarity clusters, we selected as its representative sequence the longest sequence within the cluster.

Within each cluster, we selected non-sciarid DIAMOND hits of the representative sequence from the clusteredNR database (https://ftp.ncbi.nlm.nih.gov/blast/db/experimental/, access date 5/26/2026), filtering by E-value < $10^{-3}$ and > 30% alignment overlap with the region of the query that aligns with its top non-sciarid hit. For each HTG candidate, we selected the top 300 non-dipteran hits and top 30 non-sciarid dipteran hits. We additionally included the top 20 non-dipteran metazoan hits and top 20 non-metazoan hits if these hits were not represented in the top 300 non-dipteran hits. We chose this search strategy to promote inclusion of sequences from potential metazoan and non-metazoan donor lineages, as well as dipteran sequences to test the alternative hypothesis of vertical descent.

From the combined sciarid database, we selected the following proteins (E-value < $10^{-10}$ and same coverage criterion as for the clusteredNR search):

- **Other *B. coprophila* HTG candidates**: all *B. coprophila* hits to the cluster representative that were DIAMOND-identified HTG candidates, to test HTG origins via single or multiple transfer events.
- ***B. coprophila* dipteran ancestry candidates**: The top five *B. coprophila* hits of the cluster representative that were DIAMOND-identified as being of dipteran origin, along with the top 10 non-sciarid dipteran and top 10 non-dipteran arthropod hits (against clusteredNR) of the top dipteran origin *B. coprophila* protein. These proteins were included to test the hypothesis of pseudoparology[2] (i.e. coexistence of vertically inherited and horizontally acquired homologs in the *B. coprophila* genome).
- **Putative HTG homologs in other sciarids**: non-*B. coprophila* sciarid proteins with bitscore > 0.80 of the bitscore of the top non-sciarid hit for all cluster members, limited to a maximum of 15 proteins per species per query. We reasoned that these proteins were candidate homologs of the query *B. coprophila* proteins, descended from a common transfer event.

These sequences were retrieved from the NR database with the BLAST+[103] (version 2.12.0) ‘blastdbcmd’ command and aligned using MUSCLE[125,126] (version 5.1) with default parameters. Multiple sequence alignments were then successively trimmed in two ways: 1. to solely include regions overlapping with the region of the top non-sciarid hit, 2. to include columns with > 60% column occupancy with trimAl[149] (version 1.4.1). The resulting trimmed alignment was used to produce maximum likelihood trees with IQ-TREE[117] (version 2.2.0.3) using 1,000 iterations of ultrafast bootstrapping and automated model selection. All 918 trees with ≥ 4 sequences, containing 1,714/1,721 DIAMOND-identified putative HTGs, were analyzed in subsequent steps. Trees were annotated with taxonomic information and visualized using the iTOL Python API and web browser[118].

*Automated gene tree analysis*

We wrote an ETE[113] (version 3.1.2)-based Python script to automate analysis of the closest phylogenetic relatives (“sisters”) to putative HTGs from midpoint-rooted gene trees (Figure S2A). First, the script identified monophyletic groups of sciarid sequences consisting of at least one putative *B. coprophila* HTG in each tree’s respective similarity cluster (see

“Maximum likelihood gene tree construction”). For each such monophyletic group, potential donors were identified as the immediate sister sequence or multi-sequence clade. For all multi-sequence sister clades, the potential donor taxon was determined as the lowest common ancestor taxon of all non-sciarid sister sequences using the ‘ncbi.get_topology’ function in ETE. Automated tree analysis results per HTG sequence (see below) are available in Table S3 (sheet “sister_automated_inspection”).

*Manual gene tree inspection and HGT inference*

To validate the inferences of our automated tree analysis, and also consider the evolutionary information potentially contained in non-sister clades of the trees, we manually inspected all the gene trees analyzed above. The taxonomic label of non-sciarid sequences in each tree was assigned as follows according to NCBI taxonomic ranks:

1. “Diptera” for non-sciarid dipterans.
2. Taxonomic order for non-dipteran insects.
3. Taxonomic class for non-insect arthropods.
4. Taxonomic phylum for non-arthropod metazoans.
5. Kingdom or higher level for non-metazoans, including Fungi, Viridiplantae, Viruses, and Bacteria. All Eukaryotes outside Fungi, Viridiplantae, and Metazoa were assigned the paraphyletic label “protist” for simplicity, and due to the unresolved deep branching relationships of Eukarya[137,138].

Within each gene tree, we separately examined the phylogenetic context surrounding each monophyletic group of sciarid sequences (hereafter referred to as HTG clades) containing at least one *B. coprophila* sequence in the tree’s respective similarity cluster. Trees were midpoint rooted by default, with alternative non-sciarid outgroups selected (55 cases) when midpoint rooting led to ambiguous topologies with polyphyly in sister or cousin clades, or placed sciarid sequences as the outgroup to a larger clade of non-sciarid sequences. Alternative outgroups were selected according to known lineage relationships between taxonomic kingdoms and domains, animal phyla[158], and/or arthropod classes[159] when possible, and at random for clades of bacterial or fungal sequences. HTG clades were assigned to topological categories according to the following rules:

1. **Reject** (Figure S2G). Sciarid sequences had sister sequences from non-sciarid dipteran taxa, or were not separated from a multi-sequence non-sciarid dipteran clade by at least one node with > 70% bootstrap support. These topologies are consistent with vertical descent from a dipteran common ancestor.
2. **Other direction transfer** (Figure S2F). Non-sciarid sequences were nested within an otherwise monophyletic group of dipteran sequences in which they were most closely related to sciarid sequences. These topologies are consistent with a transfer event from sciarids to non-sciarids and were excluded as a possible transfer to sciarids.
3. **One-step transfer** (Figure 3A1-A2, Figure S3). Closest (“sister”) and second closest (“cousin”) non-sciarid clades to an HTG clade were of the same taxonomic label such

that the sciarid clade is nested within an otherwise monophyletic group of non-sciarid sequences. This topology allowed for unambiguous inference of a non-sciarid origin in the taxon of the sister/cousin clades. In cases in which sisters and cousins both consisted of non-dipteran arthropod sequences of different classes, we considered this a one-step transfer from the arthropod taxon of the sister clade. Similarly, when both sisters and cousins were non-metazoan of different kingdoms or domains, we considered this a one-step transfer from the taxon of the sister clade.

4. **Two-step transfer** (Figure 3A3, Figure S4)**.** In these topologies, the sciarid HTG clade is nested within a non-sciarid metazoan donor clade ("A") consisting of sequences from a single phylum, and the combined sciarid + metazoan clade was in turn nested within a non-metazoan clade ("B"). This is consistent with two successive horizontal transfer events: non-metazoan clade B to metazoan clade A, followed by transfer from clade A to sciarids. The metazoan donor is inferred from the sister clade within A, and the non-metazoan donor is inferred from the sister clade within B.
5. **Two-step, other direction transfer** (Figure S2C, Figure S4C)**.** These cases consisted of non-sciarid metazoan sequences ("A") nested within a paraphyletic group of sciarid sequences, which were in turn nested within a different non-metazoan clade ("B"). This topology is consistent with a transfer from non-metazoan clade B to sciarids, followed by a transfer to non-sciarid clade A. All sciarid sequences within the paraphyletic group were considered to be the results of a single HGT event.
6. **Two-step, unclear direction transfer** (Figure S2D)**.** A sciarid HTG clade and non-sciarid metazoan clade consisting of sequences of at most one phylum were each other's closest relatives, without nesting of one clade within the other. These two clades were nested within a clade of non-metazoan sequences. We interpreted this topology as consistent with either a two-step transfer or multiple independent transfer events from a non-metazoan donor to sciarids and the clade of the sister sequences.
7. **Ambiguous transfer** (Figure S2E, Figure S7)**.** Like other HGT categories, these tree topologies are inconsistent with dipteran ancestry due to non-dipteran sister clades. These topologies did not fall into any of the above categories with straightforward interpretation of transfer source and/or direction due to polyphyly of sister clades, or mismatch between the taxonomic label of sister and cousin clades. Following prior work in which HGT was inferred even when topologies did not unambiguously suggest a single donor source[160], we consider these cases consistent with HGT based on their non-dipteran sister clades.

*Analysis of intron position conservation for two-step transfers*

We first collected 34 *B. coprophila* HTGs satisfying two criteria: (1) a manual tree annotation of *Two-step* or *Two-step, unclear direction*, with an inferred transfer from Bacteria→ Collembola → Sciaridae; and (2) top collembolan DIAMOND hit in *Folsomia candida*. *F. candida* hits were selected exclusively from the RefSeq annotation (GCF_002217175.1). To identify conserved intron positions, we first pairwise aligned protein sequences of each of the 34 *B. coprophila* proteins with its top *F. candida* hit using MUSCLE[125,126] (version 5.1) with default parameters. Protein alignments and nucleotide coding sequences (CDSs) were used to generate codon alignments with PAL2NAL[131] (version 14.1), on which GFF-derived positions of the final base of each coding exon were annotated. Positions of the resulting annotated alignment in which both *B. coprophila* and *F. candida* shared a final position of an exon were considered conserved intron positions (Table S9).

*Analysis of putative convergent transfers to other metazoans*

To identify possible instances of convergent horizontal transfer events to non-sciarid metazoans and sciarids, we manually inspected *one-step* (Figure 3A1, A2), *two-step* (Figure 3A3), *two-step, other direction* (Figure S2C), and *two-step unclear direction* (Figure S2D) gene trees with evidence of gene origination in a non-metazoan taxon for clades of non-sciarid metazoan sequences that met all of the following criteria:

1. In the same gene tree as a *B. coprophila*-containing sciarid clade with an HTG confirmed via manual inspection.
2. Not more closely related to the confirmed sciarid HGT clade(s) than the sciarid clade(s) are to their closest non-metazoan relatives.
3. Contained sequences from no more than one metazoan phylum, to reduce spurious detection of vertically inherited sequences.

We reported the number of sciarid HGT events that co-occurred with putative transfers to other metazoans per non-sciarid recipient taxon in Table S10. We caution that these inferences of independent HGT events in non-sciarids are preliminary, given that we did not investigate the potential for contamination as an explanation for these sequences in non-sciarid genomes. Illustrative trees are provided in Figure S5.

*Gene family expansion analysis: post-acquisition gene duplication*

To quantify post-acquisition gene duplication, we computed the total number of *B. coprophila* genes contained in each confirmed HGT sciarid clade, where each *B. coprophila* gene within a multi-sequence clade was interpreted as a paralog. We limited our tabulation (Table S4) to only those genes that were identified as HTG candidates via both DIAMOND and phylogenetic inference, and appeared in the tree's corresponding similarity cluster. In total, we found 247 HTG clades with at least one inferred post-acquisition gene duplication by these criteria.

*Gene family expansion analysis: multiple putative independent transfers to the B. coprophila genome*

All clades of confirmed HTGs that were found in the same gene tree as at least one other HTG clade were considered the products of multiple independent transfers. For each tree, we relaxed the requirement that all sciarid clades analyzed per tree contained at least one representative of the tree's DIAMOND similarity cluster, such that sciarid clades containing sequences belonging to a different tree's similarity cluster (but independently confirmed as HTGs in their respective trees) were also considered as possible instances of independent HGT.

*Expansions of gene families via HGT of genes with pseudoparalogs in the B. coprophila genome*

We searched trees with at least one HGT clade of inferred one-step origin from metazoan sources for other *B. coprophila* sequences of dipteran ancestry. We inferred dipteran ancestry for sequences in which the top non-sciarid DIAMOND hit was dipteran, and the sequence grouped with non-sciarid dipteran sequences in the gene tree. Trees containing these sequences alongside HTG clades supported gene family expansion via co-occurrence of vertically and horizontally inherited genes ('pseudoparalogs'[2]) in the *B. coprophila* genome. The numbers of HTG clades and HTG sequences found in these trees are reported in Figure S2J.

**Analysis of HTG synteny between *B. coprophila* and other sciarid genomes**

We first searched for reciprocal best BLASTp hits (RBBs) between the genome annotations for *B. coprophila* (GCF_014529535.2), and four other sciarid genome annotations (*B. odoriphaga* [GCA_016920775.1], *B. impatiens* [BRAKER annotation], *L. ingenua* [BRAKER annotation], and *P. hygida* [GCA_029228625.1]) using DIAMOND (parameters --evalue 1e-20 -k 1 --very-sensitive), restricting our search to the single longest protein isoform per protein-coding gene.

We obtained gene localizations from gff files. To each locus with an RBB, we assigned a unique numeric identifier common to both *B. coprophila* and non-*B. coprophila* proteins in the RBB. For each RBB and gene distance δ, we queried the RBB identifiers δ genes away on the same scaffold (Figure S1E-F). The gene distance δ excludes any locus without an RBB. If an RBB identifier δ genes away from the query gene (on either side) was common to both *B. coprophila* and the corresponding non-*B. coprophila* species, then we tabulated the RBB as having a conserved RBB δ away. We performed RBB analysis separately on dipteran genes and HTGs, and plotted the fraction of genes with RBBs with a syntenic neighbor over varying values of δ.

### Search for possible HGT donor sequences in a putative *B. coprophila* endosymbiont genome

We obtained the metagenome-assembled genome of a putative *B. coprophila* rickettsial endosymbiont[93] from GenBank (accession JAHXDM000000000.1) and submitted it to Prokka[135] (version 1.14.6) for annotation. We used the resulting protein FASTA to build a DIAMOND database and then queried via DIAMOND BLASTp (parameters --evalue 0.001 -k 10 --sensitive) with a FASTA of putative *B. coprophila* HTGs. For each *B. coprophila* HTG with a hit in the predicted endosymbiont genome, we compared the top bitscore of the alignment to an endosymbiont protein to the top bitscore from NR.

### Liftoff annotation transfer

We used Liftoff[122] (version 1.6.3) to transfer the annotation (gff file) from GCF_014529535.2 to the independently assembled GCA_965233685.1[154], using the following parameters: -p 16 -polish -flank 0.25 -a 0.9 -s 0.9. 1,392 HTGs transferred with an identical open reading frame; 63 transferred with a modified but intact open reading frame; 31 transferred with a partial open reading frame; 31 did not transfer at all.

### Analysis of Oxford Nanopore (ONT) reads

We downloaded Oxford Nanopore read FASTQ files for 17 runs associated with NCBI BioProject PRJNA291918 from NCBI using the SRA Toolkit[144] (version 3.1.1). We mapped reads to the *B. coprophila* genome assembly (accession GCF_014529535.2) using the 'map-ont' function of Minimap2[124] (version 2.28). We excluded SRR9971648 from subsequent analysis due to zero successfully mapped reads. We converted the resulting SAM files to sorted BAM files using SAMtools[140] 'view' and 'sort' (version 1.21), and subsequently to BED files using the 'bamtobed' function of BEDTools[101] (version 2.31.1). We extracted BED files for annotated genes from the gff file for GCF_014529535.2. The BEDTools 'intersect' function was used to determine the protein-coding genes overlapping each ONT read. The BEDTools 'coverage' function was used to determine the number of reads mapping to each protein-coding gene. Reads were pooled across runs and normalized by gene length (in kilobases). To reduce the influence of multi-mapping (e.g., to recently diverged paralogs) and low-quality alignments, secondary alignments were excluded with Minimap2 ("–secondary=no"), and SAMtools view was run excluding unmapped reads, reads not in the primary alignment, supplementary alignments, and alignments with MAPQ quality score < 30 ("-F 2308 -q 30"). Chimeric reads were defined as reads containing at least one dipteran ancestry gene and at least one HTG.

### Nucleotide composition analyses

We analyzed GC content and codon usage for dipteran ancestry genes and HTGs. GC content and GC3 values were inferred using Biopython[102] (version 1.83). Codon usage was first analyzed using MILC (Measure Independent of Length and Composition) values[1], with expected codon usage set to transcriptome-wide frequency vectors for all dipteran ancestry genes. MILC,

GC, and GC3 values were also computed for the longest CDS sequence per protein-coding gene in the RefSeq annotation for *Folsomia candida*[53] (accession GCF_002217175.1, access date 6/28/2026). MILC reference values for *F. candida* were set to the *B. coprophila* transcriptome-wide reference to measure deviation from *B. coprophila* genes. We also visualized codon usage using principal component analysis on standard-scaled codon frequency vectors using Scikit-learn[141] (version 1.2.0). Two-sample Kolmogorov-Smirnov tests were performed with SciPy (version 1.8.1).

**Arthropod lineage tree construction**

To determine the relationships between putative arthropod donor organisms and sciarids, we obtained a reference arthropod lineage tree by integrating and pruning published time-calibrated phylogenomic trees for Hexapoda[161], Chelicerata[162], Pancrustacea[163], Myriapoda[164], and Diptera[165]. Newick trees were obtained directly from the publication for Pancrustacea[163], with permission from TimeTree.org[166] for Hexapoda[161], Chelicerata[162], and Myriapoda[164] , and using Treemble (version 1.6.2)[147] to extract from Figure S3 for Diptera[165]. Backbone-level relationships among Hexapoda, Chelicerata, and non-hexapod Pancrustacea were selected from a published pancrustacean tree[163]. The NR database consists of sequences derived from genome annotations as well as protein- or gene-specific studies[167], but we reasoned that species with annotated genomes were most likely to be detected as potential HTG donors. We therefore pruned the lineage tree such that it only contained non-dipteran orders represented in GenBank- or RefSeq-annotated genomes as of 5/27/26 (NR access date). We additionally included in the tree the order Notostraca, which lacked a GenBank annotated genome but was a top DIAMOND hit for two HTGs; all other orders of top DIAMOND hits were represented by annotated genomes. Dipteran families were included to allow for within-Diptera losses in analysis of the minimum number of losses under parsimony. Families belonging to the infraorders Muscomorpha and Culicomorpha were collapsed to the infraorder level.

**Analysis of alternative hypothesis of vertical descent with differential loss**

We considered both a probabilistic model and empirical percent identity scores to evaluate the collembolan-affinity putative HTGs.

*Probabilistic expectation of collembolan-affinity genes under vertical inheritance and loss*

We computed the maximum-likelihood estimate of the probability of gene presence in Collembola (considering a single order only) and Sciaridae alone, assuming its presence in the hexapod common ancestor and allowing for recurrent gene loss. For this estimation, we adapted a previously described dynamic programming scheme[115] that considers all possible histories of losses consistent with the gene distribution across the phylogeny.

First, we estimated *P(phyletic pattern|T,u)*, where *T* is the time-calibrated tree of hexapod relationships (see "Arthropod lineage tree construction" above), *phyletic pattern* is the distribution of the gene at the tree tips (in this case, Collembola and Sciaridae only) and *u* is a free parameter reflecting the (constant) rate of gene loss. The previously derived recursion[115] for $-log_{10}[P(phyletic\ pattern|T,u)]$ was minimized with respect to *u* using optimize.fmin in SciPy[142] (version 1.8.1). To compute the probability mass function shown in Figure 2D, the conditional probability under the maximum-likelihood loss rate was treated as a binomial probability parameter *p*, with *N* = 20,000 to approximate the order of magnitude of the number of genes in the collembolan-sciarid common ancestor. This mass function was compared to the number of HTGs with top DIAMOND hit in Collembola and the number of *one-step* (Figure 3A1, A2) or *two-step* (Figure 3A3) transfer events of collembolan affinity.

*Collembolan percent identity analysis*

First, we collected a set of tree-validated HTGs of inferred collembolan ancestry, defined as all *B. coprophila* proteins inside HGT clades with *one-step* or *two-step* annotations in which the inferred donor lineage was Collembola. For each of the resulting 646 HTGs, we then extracted the DIAMOND percent identity to the lowest E-value collembolan hit, and compared this statistic to the percentage identity of DIAMOND-defined dipteran ancestry proteins to their respective lowest E-value collembolan DIAMOND hits (for all 9,664 dipteran genes with collembolan hits E-value < 0.01 in the top 2,000 hits). Histograms were plotted using the 'histplot' function in seaborn[143] (version 0.13.2).

**Genome localization analysis**

Localization of HTGs and dipteran genes was analyzed using the gff file for the *B. coprophila* reference annotation (GCF_014529535.2). A chi-squared contingency test was performed with SciPy (version 1.8.1) on the distribution of HTGs across each of the four chromosomes in this assembly relative to the distribution of DIAMOND-identified dipteran-ancestry (*dipt*) genes. The Circos plot of HTG and dipteran gene density per 100,000 base pairs was generated using pyCirclize[136] (version 1.7.1).

A null distribution for the proportion of HTGs immediately adjacent to another HTG (i.e. with no other intervening genes) was generated by 10,000 iterations of the following procedure:

1. Drawing a randomly selected subset of 1,517 genes from the *B. coprophila* genome, equivalent to the observed number of tree-validated HTGs.
2. Assigning the randomly selected subset a "simulated HTG" label.
3. Computing the proportion of simulated HTGs immediately adjacent to other simulated HTGs, pooling across all chromosomes and scaffolds.

To generate Figure 3D, genes encoding top DIAMOND hits to *B. coprophila* HTGs were selected for visualization for the collembolan *F. candida* and tardigrade *Paramacrobiotus metropolitanus* (the latter only for β-lactam synthesis genes). Genomic localization of glyoxylate shunt and β-lactam biosynthesis genes along with associated RNA-Seq tracks was obtained from the NCBI Genome Data Viewer[127]. Intergenic regions were removed and genes were rescaled for ease of visualization. Supporting amino acid and codon alignments are available from the accompanying Dryad repository under "pairwise_folsomia_alignments_intron" and "pairwise_paramacrobiotus_alignments_intron.tar.gz".

### Transcriptomic analysis

Raw RNA-Seq counts and transcripts per million (TPM)-normalized expression values were obtained from the NCBI ftp site associated with the reference *B. coprophila* annotation (https://ftp.ncbi.nlm.nih.gov/genomes/all/GCF/014/529/535/GCF_014529535.2_BU_Bcop_v2.1, access date 7/7/2026). Mean expression data in Figure 4B were pooled across BioProject accessions PRJNA1109384, PRJNA1220056, PRJNA291918, PRJNA928089; Figure 4A and 4C contain non-salivary gland data from PRJNA291918; and Figure S8C-D were generated using PRJNA1109384 and salivary gland data from PRJNA291918. Pairwise differential expression analyses were executed with DESeq2[111] (version 1.38.3) in R (version 4.2.3) on raw counts for the following comparisons:

1. Larval vs. non-larval expression from BioProject PRJNA291918, excluding all salivary gland transcriptomes.
2. Irradiated larval expression vs control (non-irradiated) larval samples from BioProject PRJNA928089.

Heatmaps were generated with the seaborn (version 0.13.2) clustermap function. Stage or tissue of maximal expression was obtained from TPM values pooled per stage or tissue. Pearson correlation coefficients of glyoxylate shunt enzyme expression with phosphoenolpyruvate carboxykinase 1 (PEPCK1, RefSeq identifier Pepck1) and carnitine palmitoyltransferase 1 (CPT1, RefSeq identifier whd) expression were computed from raw TPM values across all developmental stages in BioProject PRJNA291918.

### Sciarid species tree inference

We extracted 760 complete and single-copy BUSCO orthologs[106] from eight dipteran NCBI genome annotations, two sciarid genomes we annotated via BRAKER3 (see "Sciarid database construction" section above), and two unannotated dipteran genomes (Table S28) using the BUSCO software[107] (version 5.7.1) and the diptera_odb10 BUSCO set (release 2024-01-08). The BUSCO Phylogenomics pipeline[168] (release 20240919) was used to generate multiple sequence alignments and concatenate trimmed alignments into a supermatrix. The supermatrix was used for species tree inference using IQ-TREE (version 2.2.0.3) with automated model selection and 1,000 iterations of ultrafast bootstrapping. The resulting species tree topology (Figure S8A) agrees with previously published relationships among sciarid species[169] and dipteran families[170]. Partitioned tree inference yielded an identical topology, so subsequent analyses were performed on the unpartitioned supermatrix.

We used the approximate likelihood method implemented in MCMCtree[133,171] (PAML version 4.10.6) to produce a time-calibrated phylogeny from literature-derived fossil constraints and the maximum likelihood topology and concatenated supermatrix specified above (Tables S29-S30). Following Álvarez-Carretero and colleagues[172], we specified the gamma rate prior by assuming an unrestrictive value of 2 for the shape ('alpha') parameter, and set the 'beta' parameter such that the mean rate equaled the *B. coprophila* to root distance on the maximum-likelihood topology divided by 250 million years, an approximation of the time to the crown-group dipteran common ancestor[165]. To approximate the best-fitting model determined by ModelFinder in IQ-TREE (Q.insect+F+I+I+R9), we selected the Q.insect amino substitution matrix[173] and modeled among-site heterogeneity with a discrete gamma distribution (ncatG = 9, alpha = 0.5). Due to the close phylogenetic relatedness of the multiple Sciaroidea genomes in our phylogeny, we selected the autocorrelated rates model for our molecular clock. We discarded the first 100,000 iterations of MCMC sampling as burn-in, and subsequently collected samples every 100 iterations until a total of 300,000 samples were collected.

Using Tracer[146] (version 1.7.2), we confirmed that the dipteran root and sciarid divergence time effective sample size (ESS) values were > 800, and posterior means of all parameters differed by less than two percent across two independent runs of MCMCtree. Time-calibrated species trees with highest posterior density (HPD) intervals from the first independent run were visualized with FigTree[114] (version 1.4.4).

### Phylostratigraphic dating

HTG clades on gene trees were assigned to one of five possible age categories (phylostrata[174]; Figure 1H) based on presence of sequences from the genomes of *B. odoriphaga, B. impatiens, L. ingenua, and P. hygida*. In Figure 1H and Tables S3 and S15, HTG acquisition events were assigned to the duration between (1) (minimum) the divergence of *B. coprophila* from the most distantly related species containing a gene tree-determined homolog of the *B. coprophila* sequence(s) in each HTG clade; and (2) (maximum) the divergence of *B. coprophila* from the next most closely related species lacking a gene tree-determined homolog of the *B. coprophila* sequence. For sequences found in *P. hygida* (and hence assigned to the oldest phylostratum) we considered the upper bound on the HTG acquisition as the divergence time of Sciaridae from Cecidomyiidae, since annotated Mycetophilidae or Diadocidiidae genomes were not available at the time of writing and hence were not analyzed.

### Selective pressure (dN/dS) analysis

A total of 13,046 reciprocal best DIAMOND BLASTp alignments between the *B. coprophila* reference annotation and our BRAKER3-based annotation of the *B. impatiens genome* (see "Sciarid database construction" above) were subjected to dN/dS analysis.

Pairwise amino acid alignments were first obtained with MUSCLE[125,126] (version 5.1), which were submitted alongside nucleotide CDS sequences to PAL2NAL[131] (version 14.1) to produce codon alignments. Codon alignments with dS < 2 were successfully generated for 12,764/13,046 input protein pairs, including 990 HTGs. Pairwise dN/dS estimates were subsequently obtained with CODEML in PAML[132–134] (version 4.10.6) with the M0 model (model = 0, cleandata = 1, Confer = 7). A null model of dN/dS = 1 was also fit using the same model with fix omega = 1 and omega = 1. Likelihood ratio test p-values were obtained with $\chi^2 = 2\Delta logL$ using SciPy[142] (version 1.8.1), and were false discovery-rate adjusted according to the Benjamini-Hochberg procedure implemented in statsmodels[145] (version 0.14.0).

Mann-Whitney comparisons of dN/dS between tree-inferred HTGs and DIAMOND-inferred dipteran genes were performed using SciPy[142] (version 1.8.1). To determine whether the observed increase in dN/dS for HTGs vs. dipteran genes held after adjustment for covariates, we used the 'smf.ols' tool in statsmodels[145] (version 0.14.0) to fit an ordinary least squares model for dN/dS of "dipteran" or "HGT" ancestry genes, with the following predictor variables:

- *$log_{10}(TPM+10^{-2})$*, with TPM being the mean transcripts per million expression value across all analyzed *B. coprophila* RNA-Seq datasets (BioProjects PRJNA1109384, PRJNA1220056, PRJNA291918, PRJNA928089).
- *Extracellular*: a binary variable, with "1" indicating predicted "extracellular" subcellular localization (see Methods).
- *HGT*: a binary variable, with "1" indicating tree-confirmed HGT ancestry and "0" indicating DIAMOND-inferred dipteran ancestry.
- *CDS length*: the number of nucleotides in each gene's coding sequence.

Mann-Whitney comparisons in Figure 4E were performed for 10,535 DIAMOND-inferred dipteran genes, 466 DIAMOND-inferred orphan genes (i.e., no DIAMOND hits outside of Sciaridae), 76 "middle"-aged HTGs, and 910 "old"-aged HTGs (see Methods).

### Mapping of accessions from previous *B. coprophila* genome annotation to current annotation

Our analysis used reference annotation version GCF_014529535.2, which differs from the version used in previously published studies of *B. coprophila* HGT and carbohydrate-active enzymes (GCF_014529535.1). To facilitate comparisons between these prior publications and our results (Table S1), we performed a DIAMOND BLASTp search (--very-sensitive) of proteins from the old annotation vs. the current annotation. Proteins without an exact accession match in the current annotation were mapped to their top (lowest E-value) match in the current annotation if available (pident > 95% and qcovhsp > 85%). Protein matches were mapped to their respective genes and then to the longest protein isoform selected for HGT inference in the current annotation.

### SI Tables S1-S30

SI Tables and legends are listed in numerical order. SI Tables for which only the legend is included here, are available to download as individual Excel files.

**Table S1. Previously published *B. coprophila* HTGs.** Table excludes studies of HGT and/or introgression into *Bradysia* germline-restricted chromosomes[92,199], which were not analyzed here, as well as hypothesized HGT of carbohydrate catabolism genes based upon sequence similarity and gene presence/absence patterns without gene tree validation[37]. RefSeq protein accessions from published studies were mapped to gene accessions before comparison to our set of HTGs to mitigate inconsistencies in protein isoform selection. Mappings from the previous annotation (GCF_014529535.1) to closest match in current (GCF_014529535.2) RefSeq annotation (85% length coverage, 95% identity) are indicated in the "new_accession" column, where "removed" indicates that the sequence has been removed from the current annotation with no replacement. "Confirmed" column indicates whether the gene appeared in our final set of tree-validated HTGs.

| Annotation | published_accession | new_accession | Confirmed | Reference |
|---|---|---|---|---|
| Cytochrome P450 | XP_037037647.1 | | Yes | [51] |
| Cytochrome P450 | XP_037047887.1 | XP_080113996.1 | Yes | [51] |
| Cytochrome P450 | XP_037048891.1 | XP_080119038.1 | Yes | [51] |
| Cytochrome P450 | XP_037051484.1 | | Yes | [51] |
| Cytochrome P450 | XP_037049572.1 | | Yes | [51] |
| Cytochrome P450 | XP_037025072.1 | XP_080120198.1 | Yes | [51] |
| Cytochrome P450 | XP_037036945.1 | XP_037036944.1 | Yes | [51] |
| Cytochrome P450 | XP_037042570.1 | | Yes | [51] |
| Cytochrome P450 | XP_037042572.1 | | Yes | [51] |
| Cytochrome P450 | XP_037052143.1 | | Yes | [51] |
| Cytochrome P450 | XP_037034543.1 | | Yes | [51] |
| Cytochrome P450 | XP_037044115.1 | XP_080112887.1 | Yes | [51] |
| Cytochrome P450 | XP_037050080.1 | | Yes | [51] |
| Cytochrome P450 | XP_037045845.1 | | Yes | [51] |
| Cytochrome P450 | XP_037025508.1 | | Yes | [51] |
| Cytochrome P450 | XP_037036450.1 | | Yes | [51] |
| Cytochrome P450 | XP_037049686.1 | XP_080118275.1 | Yes | [51] |
| Cytochrome P450 | XP_037049098.1 | XP_037049099.1 | Yes | [51] |
| Cytochrome P450 | XP_037029925.1 | XP_080112043.1 | Yes | [51] |
| Cytochrome P450 | XP_037049483.1 | | Yes | [51] |
| Cytochrome P450 | XP_037025913.1 | | Yes | [51] |
| Cytochrome P450 | XP_037037225.1 | | Yes | [51] |
| Cytochrome P450 | XP_037035924.1 | | Yes | [51] |
| Cytochrome P450 | XP_037043806.1 | | Yes | [51] |
| Cytochrome P450 | XP_037047695.1 | | Yes | [51] |
| Cytochrome P450 | XP_037052058.1 | | No | [51] |

| | | | | |
|---|---|---|---|---|
| Cytochrome P450 | XP_037032840.1 | | Yes | [51] |
| Cytochrome P450 | XP_037029512.1 | removed | No | [51] |
| Chimeric gene | XP_037032039.1 | removed | No | [200] |
| Chimeric gene | XP_037036061.1 | | Yes | [200] |
| Chimeric gene | XP_037047876.1 | XP_037047876.2 | Yes | [200] |
| Chimeric gene | XP_037051404.1 | XP_080119310.1 | Yes | [200] |
| Chimeric gene | XP_037041958.1 | XP_080119927.1 | Yes | [200] |
| Chimeric gene | XP_037048916.1 | XP_080113585.1 | Yes | [200] |
| Chimeric gene | XP_037049533.1 | | No | [200] |
| Chimeric gene | XP_037030969.1 | | No | [200] |
| Chimeric gene | XP_037026007.1 | | No | [200] |
| Chimeric gene | XP_037047875.1 | XP_037047875.2 | Yes | [200] |
| Chimeric gene | XP_037038002.1 | XP_080118104.1 | No | [200] |
| Terpene synthase | XP_037024969.1 | XP_080120004.1 | Yes | [67] |
| Terpene synthase | XP_037027991.1 | | Yes | [67] |
| Terpene synthase | XP_037028003.1 | | Yes | [67] |
| Terpene synthase | XP_037049366.1 | XP_080114915.1 | Yes | [67] |
| Tryptophanase/tryptophan-indole lyase (TNase) | XP_037029494.1 | | Yes | [201] |
| Alternative oxidase (AOX) | XP_037036163.1 | | Yes | [39,59] |
| DASH cryptochrome/photolyase | XP_037034936.1 | | Yes | [60,61] |
| Uncharacterized β-propeller gene (propellin) | XP_037046651.1 | removed | No | [202] |
| Glycoside hydrolase family 32 (GH32) | XP_037030094.1 | | No | [35] |
| Glycoside hydrolase family 32 (GH32) | XP_037025415.1 | | Yes | [35] |
| Glycosyl hydrolase group 43 (GH43) | XP_037044875.1 | removed | No | [36] |
| Apoptosis inducing protein 56 (toxin) | XP_037050353.1 | removed | No | [203] |
| Lysozyme | XP_037029278.1 | | Yes | [203] |
| Bacterial extracellular solute-binding protein | XP_037031050.1 | XP_037031050.2 | Yes | [204] |

**Table S2 (attached file). Per-genome results of DIAMOND screen for HGT across 54 dipteran genomes**. Columns are as follows: "Genome" is the RefSeq accession of the genome analyzed; "Organism Name", "family", and "taxid" provide NCBI taxonomic information for the genome analyzed; "orphan", "dipt", "alien", "non_dipt", "none" quantify the number of protein ancestries in each ancestry class described in Methods; "HGT" and "p_HGT" are the total number and percentage of DIAMOND-inferred HTGs in the genome computed as the sum of proteins in the "alien" and "non_dipt" categories.

**Table S3 (attached file). HGT gene tree analysis.** Results of maximum-likelihood gene tree inspection, in two sheets:

- **sister_automated_inspection:** Taxonomic information of closest non-sciarid phylogenetic relatives ("sisters") to manual inspection-confirmed *B. coprophila* sequences, determined via automated analysis of midpoint-rooted trees. Each row corresponds to a single HTG ("accession") confirmed via manual inspection. Columns are as follows: "tree_number" is a unique numerical identifier for each tree available on iTOL, with some trees containing multiple clades/rows; "name" and "taxid" are the taxonomic name and NCBI taxid of the lowest common taxonomic rank to all sequences in the sister clade per HGT clade; remaining columns are various higher taxonomic ranks if applicable to the rank in "name" else blank.
- **manual_inspection_merged:** Results of manual inspection of gene trees, per clade of sciarid sequences. Clades in this table are restricted to those containing at least one *B. coprophila* sequence in each tree's respective similarity cluster. Column description: "tree.cluster" a unique identifier for each clade, with format "iTOL tree identifier.clade number"; "annotation", inferred HGT or non-HGT annotation for each clade; "primary neighbor" the taxonomic labels of donors for "One step" and "Two step" transfers, closest non-sciarid relatives for "Two step unclear direction" (possibly donors to sciarids or recipients of independent transfer events), recipient lineage for "Other direction" and "Two-step other direction" transfers; "secondary neighbor" are donors to the proximate neighbors for "Two-step" transfers, either donors to both sciarids and non-sciarid metazoans, or donors to proximate neighbors for "Two step unclear direction", and donors to sciarids for "Two step other direction"; "re-root" indicates manually selected outgroup rooting scheme if applicable, and is blank for default midpoint rooting; "notes" are any specific factors considered for HGT inference of this clade; "all_seqs" are all sequences in the clade, including both *B. coprophila* sequences and those from other sciarids (GenBank accessions for *P. hygida* and *B. odoriphaga*; accessions generated via BRAKER3 annotation for *L. ingenua* and *B. impatiens*); "cluster_seqs" are *B. coprophila* sequences in the clade that were identified as HTGs via DIAMOND and are in the tree's respective similarity cluster; "n_cluster_seqs" are the number of sequences in "cluster_seqs"; "sciarid_species" is a list of all sciarid species with sequences in the clade; "age" is the phylostratigraphic age category from youngest (1) to oldest (5) assigned according to "sciarid_species"; "convergent" contains the metazoan taxon/taxa inferred to have independently received the HTG, if detected (otherwise blank); "pseudoparalog" is TRUE for metazoan origin HGT-confirmed clades that co-occur with an inferred vertically-inherited/dipteran-ancestry sciarid clade; "multi_transfer" is TRUE for HGT clades inferred to represent independent transfer events to *B. coprophila* in the same tree (including those not in each tree's respective similarity cluster and hence absent from this table). Blank entries in the latter two columns represent clades not assessed for pseudoparalogy or multiple transfers.

- **Table S4. Distribution of the numbers of inferred *B. coprophila* genes per HGT clade**. 247 clades with > 1 *B. coprophila* sequence are inferred to contain paralogs resulting from post-transfer gene duplication. Tabulation excludes *B. coprophila* proteins in HGT clades that did not pass DIAMOND-based thresholds.
-

| Number of genes | Number of transfers |
|---|---|
| 1 | 687 |
| 2 | 127 |
| 3 | 54 |
| 4 | 24 |
| 5 | 16 |
| 6 | 7 |
| 7 | 6 |
| 8 | 4 |
| 9 | 3 |
| 13 | 2 |
| 16 | 2 |
| 17 | 1 |
| 20 | 1 |

**Table S5 (attached file). HTG presence across other genomes,** in two sheets:

- **other_sciarids.** Gene-tree-based assessment of presence of homologs of *B. coprophila* HTGs in the genomes of the other sciarid species *B. odoriphaga, B. impatiens*, *L. ingenua*, and *P. hygida* (“TRUE” if a homolog is detected in the column species). “phylostratum” provides the age category from youngest (1) to oldest (5) based on the most distantly related sciarid species in which a homolog is found (tabulated in Table S15).
- **other_b.coprophila_genome.** Assessment of HTG presence in the independent *B. coprophila* genome assembly GCA_965233685.1 using Liftoff. Value descriptions for “GCA_965233685.1_liftoff” are as follows: “full” indicates the full ORF transferred to the updated assembly with identical protein sequence; “modified_ORF” indicates transfer with a modified protein sequence; “partial_ORF” indicates that the transferred ORF is incomplete; “no_transfer” indicates that the sequence could not be transferred to the new assembly by Liftoff. “full" and “modified_ORF” transfers were considered intact for the purpose of this analysis.

**Table S6 (attached file)**. **Per-gene results of DIAMOND screen for HGT across the *B. coprophila* genome**. Columns are as follows: RefSeq “accession” of the protein; “description” NCBI RefSeq annotation description of the protein; “gene”: accession of the gene; “n_non_diptera”: number of non-dipteran protein hits; “n_out_fam_diptera”: number of non-sciarid dipteran hits; “AI”: alien index as explained in SI Methods; “sseqid”, “sciname”, “tid” (NCBI taxid), “evalue”, “bitscore”, “qcovhsp”, “qstart”, “qend”, and “pident” indicate standard DIAMOND output fields; “length”: length of the protein; “type”: annotation to “orphan”, “dipt”, “alien”, “non_dipt”, or “none” categories as described in SI Methods; “domain”, “kingdom”, “phylum”, “class”, “order” are taxonomic information of the top non-sciarid hit; “BLAST_HTG”: TRUE if protein was determined as a putative HTG via DIAMOND; “tree_HTG”: TRUE if protein was confirmed as an HTG via manual inspection of gene trees. Seven proteins that failed to return any DIAMOND hits were excluded from this table.

**Table S7. Top 20 species by top non-sciarid DIAMOND hits to HTGs**. "N_HTGs" indicates the number of DIAMOND-defined HTGs with a top non-sciarid DIAMOND hit in the "Species" column. "Soil-dwelling" indicates whether the species or corresponding genus has a reported niche in soil, leaf litter or moss according to the cited source. All species with at least seven HTGs are reported. Note: Collembola is considered a class in the NCBI Taxonomy database[175]. *Rotaria* Silwood species were sampled from a pond subject to periodic drying[176] and are members of a genus that harbors other soil-dwelling species[177–179].

| **Species** | **Domain** | **Kingdom** | **Phylum** | Class | **Soil-dwelling** | **N_HTGs** |
|---|---|---|---|---|---|---|
| *Folsomia candida* | Eukaryota | Metazoa | Arthropoda | Collembola | Yes[53] | 461 |
| *Allacma fusca* | Eukaryota | Metazoa | Arthropoda | Collembola | Yes[180] | 133 |
| *Orchesella dallaii* | Eukaryota | Metazoa | Arthropoda | Collembola | Yes[181] | 129 |
| *Orchesella cincta* | Eukaryota | Metazoa | Arthropoda | Collembola | Yes[52] | 61 |
| *Halotydeus destructor* | Eukaryota | Metazoa | Arthropoda | Arachnida | Yes[182] | 59 |
| *Oppia nitens* | Eukaryota | Metazoa | Arthropoda | Arachnida | Yes[183] | 32 |
| *Rotaria* sp. Silwood1 | Eukaryota | Metazoa | Rotifera | Bdelloidea | Yes[176–178] | 18 |
| *Leptotrombidium deliense* | Eukaryota | Metazoa | Arthropoda | Arachnida | Yes[184] | 17 |
| *Archispirostreptus syriacus* | Eukaryota | Metazoa | Arthropoda | Diplopoda | Yes[185] | 15 |
| *Venturia canescens* | Eukaryota | Metazoa | Arthropoda | Insecta | No | 13 |
| *Oppiella nova* | Eukaryota | Metazoa | Arthropoda | Arachnida | Yes[186] | 11 |
| *Medioppia subpectinata* | Eukaryota | Metazoa | Arthropoda | Arachnida | Yes[187] | 10 |
| *Adineta steineri* | Eukaryota | Metazoa | Rotifera | Bdelloidea | Yes[177–179] | 9 |
| *Rotaria* sp. Silwood2 | Eukaryota | Metazoa | Rotifera | Bdelloidea | Yes[176–178] | 8 |
| *Adineta ricciae* | Eukaryota | Metazoa | Rotifera | Bdelloidea | Yes[177–179] | 8 |
| *Pectobacterium aroidearum* | Bacteria | Pseudomonadati | Pseudomonadota | Gammaproteobacteria | Yes[188] | 8 |
| *Adineta vaga* | Eukaryota | Metazoa | Rotifera | Bdelloidea | Yes[177,178] | 8 |
| *Blomia tropicalis* | Eukaryota | Metazoa | Arthropoda | Arachnida | No | 8 |
| *Rotaria sordida* | Eukaryota | Metazoa | Rotifera | Bdelloidea | Yes[178] | 8 |
| *Ralstonia pseudosolanacearum* | Bacteria | Pseudomonadati | Pseudomonadota | Betaproteobacteria | Yes[189] | 7 |

**Table S8. Taxonomic origins of HTGs as determined via manual inspection of gene trees.** Taxonomic labels were assigned at the class level for non-insect arthropods, order level for insects, phylum level for non-arthropod metazoans, and kingdom or domain level for non-metazoans. "Protists" encompasses eukaryotes outside Metazoa, Fungi and Viridiplantae.

| Taxon | # Transfers |
|---|---|
| A. Origins of putative one-step transfers (n = 565; Figs 3A1, A2) | |
| Collembola | 304 |
| Arachnida | 80 |
| Fungi | 60 |
| Bacteria | 41 |
| Rotifera | 16 |
| Protist | 14 |
| Hymenoptera | 13 |
| Coleoptera | 11 |
| Hemiptera | 8 |
| Diplopoda | 4 |
| Lepidoptera | 3 |
| Viridiplantae | 2 |
| Psocodea | 2 |
| Ostracoda | 2 |
| Branchiopoda | 1 |
| Blattodea | 1 |
| Mollusca | 1 |
| Thysanoptera | 1 |
| Nematoda | 1 |
| B. Primary origins of putative two-step transfers, determined as the likely donor lineage to sciarids (n = 102; Fig. 3A3) | |
| Collembola | 82 |
| Rotifera | 14 |
| Arachnida | 6 |
| C. Secondary origins of putative two-step transfers, determined as the likely donor to the primary donor lineage (n = 102) | |
| Fungi | 44 |
| Bacteria | 39 |
| Ambiguous | 14 |
| Protist | 4 |
| Viruses | 1 |
| D. Primary origins of putative two-step unclear direction transfers, determined as the likely donor to sciarids, or recipient of a convergent transfer event from a non-metazoan donor (n = 65; Fig. S2D) | |
| Collembola | 50 |

| | |
|---|---|
| Rotifera | 12 |
| Arachnida | 3 |
| E. Second-closest non-sciarid relatives for two-step unclear direction transfers, which are interpreted either as donors to both sciarids and their proximate neighbors (above) or donors to the proximate neighbors (n = 65 Fig. S2D) | |
| Bacteria | 24 |
| Fungi | 21 |
| Protist | 12 |
| Ambiguous | 7 |
| Viridiplantae | 1 |
| F. Inferred recipients for putative other direction transfers, from sciarids to non-sciarids (n = 8; Fig. S2F) | |
| Collembola | 4 |
| Bacteria | 1 |
| Branchiopoda | 1 |
| Arachnida | 1 |
| Archaea | 1 |
| G. Inferred donors to sciarids for putative two-step other direction transfers (n = 10; Fig. S2C) | |
| Bacteria | 6 |
| Fungi | 2 |
| Protist | 2 |
| H. Inferred recipients of HGT from sciarids for putative two-step other direction transfers (n = 10; Fig. S2C) | |
| Collembola | 8 |
| Rotifera | 2 |

**Table S9 (attached file). Data on intron conservation with respect to *Folsomia candida*** for 34 tree-confirmed “two-step” or “two-step unclear direction” HTGs of primary collembolan origin and secondary bacterial origin with top collembolan hit in *F. candida*. Column descriptions are as follows: “Bradysia_accession” is the *B. coprophila* protein RefSeq accession; “Folsomia_accession” is the RefSeq accession of the top DIAMOND hit in *F. candida*; “Annotation” is the type of HGT determined during manual tree inspection (see SI Methods ”Manual gene tree inspection and HGT inference”); “n_int” is the number of CDS-spanning introns with conserved position in *F. candida and B. coprophila* genes (set intersection); “n_union” is the total number of CDS-spanning introns across either species (set union); “n_fols” is the number of CDS-spanning introns in the *F. candida* gene; “n_brad” is the number of CDS-spanning introns in the *B. coprophila gene*.

**Table S10. Non-sciarid recipients of convergent HGT.** Metazoan taxa inferred to have received putative convergent HGT of genes in the same tree as a transfer to sciarids, reported at the phylum level for non-arthropod metazoans, class level for non-insect arthropods, and order level for insects. "Number of sciarid transfers" indicates the number of sciarid clades inferred to co-occur in the same phylogenetic tree as a convergent HGT event to each taxon. A total of 128 sciarid HGT events were inferred to co-occur in the same tree as a transfer to another metazoan taxon. The total in this table (194) exceeds 128 because 46 transfers co-occurred with transfers to multiple metazoan groups.

| Recipient taxon | Number of sciarid transfers |
|---|---|
| Rotifera | 50 |
| Collembola (non-insect arthropod) | 21 |
| Arachnida (non-insect arthropod) | 20 |
| Hemiptera (insect) | 14 |
| Porifera | 12 |
| Nematoda | 10 |
| Branchiopoda (non-insect arthropod) | 7 |
| Cnidaria | 6 |
| Annelida | 4 |
| Diplopoda (non-insect arthropod) | 4 |
| Xenacoelomorpha | 4 |
| Lepidoptera (insect) | 4 |
| Tardigrada | 4 |
| Chordata | 4 |
| Hymenoptera (insect) | 3 |
| Ostracoda (non-insect arthropod) | 3 |
| Mollusca | 3 |
| Placozoa | 2 |
| Zygentoma (insect) | 2 |
| Ctenophora | 2 |
| Neuroptera (insect) | 2 |
| Coleoptera (insect) | 2 |
| Malacostraca (non-insect arthropod) | 2 |
| Copepoda (non-insect arthropod) | 2 |
| Amphipoda (non-insect arthropod) | 1 |
| Trichoptera (insect) | 1 |
| Siphonaptera (insect) | 1 |
| Merostomata (non-insect arthropod) | 1 |
| Diptera (insect) | 1 |
| Psocodea (insect) | 1 |
| Platyhelminthes | 1 |

**Table S11. *B. coprophila* HTGs with literature-derived evidence consistent with transfers of functionally similar HTGs to other eukaryotes.** Recipient taxa in the "other recipient(s)" column are stated at the taxonomic level reported in published studies, with the taxonomic phylum for non-arthropod metazoans and taxonomic order for arthropods indicated in parentheses. The "phytophagy-associated" column indicates whether at least some of the recipient lineage(s) are phytophagous and the HTG was previously hypothesized to support phytophagy/herbivory. CAZy family designations (https://www.cazy.org/) for carbohydrate-active enzymes are indicated in parentheses. Accessions of corresponding HTGs are provided in Table S18, sheet "carbohydrate_catabolism" for carbohydrate-related genes, and Table S18, sheet "manual_annotations" for all others.

| **Protein (predicted function)** | **N_HTGs** | **Other recipient(s)** | **Phytophagy-associated** |
|---|---|---|---|
| Cyanate hydratase (cyanide detoxification) | 1 | *Meloidogyne incognita* (Nematoda)[29,190], *Tetranychus urticae* (Trombidiformes)[191] | Yes |
| β-cyanoalanine synthase (cyanide detoxification) | 1 | Lepidoptera[192], *Tetranychus urticae* (Trombidiformes)[193] | Yes |
| Alternative oxidase | 1 | Nematoda, Collembola, Rotifera[59] | Yes |
| Arsenite methyltransferase (arsenite detoxification) | 1 | Fungi, diatoms, possibly the last metazoan common ancestor[194] | No |
| Isocitrate lyase (glyoxylate shunt) | 1 | Nematoda, Cnidaria, *Dictyostelium*, and Euglena[80] | No |
| Malate synthase (glyoxylate shunt) | 1 | Nematoda, Cnidaria, *Dictyostelium*, and Euglena[80] | No |
| Kynureninase (kynurenine pathway) | 1 | *Bombyx mori* (Lepidoptera)[195,196] | No |
| Pectin/pectate lyase (carbohydrate catabolism) | 7 | Nematoda[29] and Chrysomelidae (Coleoptera)[31] | Yes |
| Pectin methylesterase (CE8) (carbohydrate catabolism) | 2 | Curculionoidea (Coleoptera)[41] | Yes |
| Polygalacturonase (GH28) (carbohydrate catabolism) | 5 | Phytophaga (Coleoptera), Lamiinae (Coleoptera), Mirinae (Hemiptera), Bruchinae (Coleoptera), Verophasmatodea (Phasmatodea)[41]; *Aphelenchus avenae* and *Meloidogyne incognita* (Nematoda)[29], Chrysomelidae (Coleoptera)[31] | Yes |
| Endo-β-1,3-glucanase | 1 | Aphelenchoidea (Nematoda)[29] | Yes |
| Exoglucanase/xylanase (GH10) (carbohydrate catabolism) | 1 | *Hypothenemus hampei* (Coleoptera)[41], Cassidinae (Coleoptera) and Criocerinae (Coleoptera)[31] | Yes |
| Endo-1,4-β-xylanase (GH11) (carbohydrate catabolism) | 2 | *Hypothenemus hampei* (Coleoptera)[41], *Phaedon cochleariae* (Coleoptera)[31] | Yes |

| | | | |
|---|---|---|---|
| Cellulase/endoglucanase (GH45) (carbohydrate catabolism) | 7 | Phytophaga (Coleoptera)[41], Chrysomelidae (Coleoptera)[31], Aphelenchoidea (Nematoda)[29] | Yes |
| Glycosyl hydrolase, family 5 (GH5) (carbohydrate catabolism) | 7 | Lamiinae (Coleoptera)[41], Nematoda[29], Bruchinae (Coleoptera)[31] | Yes |
| Exo-α-(1-5)-L-arabinofuranosidase (GH43) (carbohydrate catabolism) | 2 | *Globodera pallida* and *Meloidogyne incognita* (Nematoda)[29], *Lema* (Coleoptera)[31] | Yes |
| Glycosyl hydrolase, family 31 (GH31) (carbohydrate catabolism) | 4 | Lepidoptera[41] | Yes |
| β-fructofuranosidase (GH32) (carbohydrate catabolism) | 1 | Lepidoptera, *Agrilus planipennis* (Coleoptera), Curculionidae (Coleoptera), *Mayetiola destructor* (Diptera), Tetranychidae (Trombidiformes)[41] | Yes |
| Pyridoxal 5'-phosphate synthase subunit PdxS/SNZ (Pyridoxal 5'-phosphate synthesis) | 1 | *Heterodera glycines* (Nematoda)[73] | Yes |
| Pyridoxal 5'-phosphate synthase subunit PdxT/SNO (Pyridoxal 5'-phosphate synthesis) | 1 | *Heterodera glycines* (Nematoda)[73] | Yes |
| Chorismate mutase (amino acid biosynthesis) | 1 | Tylenchida (Nematoda)[29] | Yes |
| Polyketide/non-ribosomal peptide synthesis-associated enzymes | 12 | Bdelloidea (Rotifera)[197] | No |
| D-alanine--D-alanine ligase (Bacterial cell wall biosynthesis) | 4 | *Planococcus citri* (Hemiptera)[94] | No |
| Alanine racemase (Bacterial cell wall biosynthesis) | 2 | SAR group protozoans; Excavata; Holomycota; *Schizosaccharomyces pombe*[198] | No |
| Domesticated amidase effector/type VI secretion amidase effector (Immune effector; Bacterial cell wall degradation/turnover) | 1 | Acari (mites and ticks), *Daphnia*, Ciliophora, Annelida and Mollusca, possibly the metazoan common ancestor[90]. | No |
| Lysozyme/endolysin (Immune effector; Bacterial cell wall degradation/turnover) | 2 | Aphidinae (Hemiptera)[99], Heterodonta and Palaeoheterodonta (Mollusca)[89] | No |
| Intradiol ring-cleavage dioxygenase (Xenobiotic metabolism) | 1 | Sarcoptiformes and Trombidiformes, Collembola, Tardigrada, *Bemisia tabaci* (Hemiptera)[50] | Yes |
| Hemolysin (Pore-forming toxin) | 1 | Drosophilidae (Diptera)[95] | No |

**Table S12 (attached file). Genomic clustering of HTGs**. Data on the genomic clustering of HTGs in the *B. coprophila* genome. Each row corresponds to a pair of HTGs that are each other's immediate genomic neighbors (i.e. no intervening genes) on the same scaffold or chromosome indicated with "location", with RefSeq gene locus identifiers in the "locus_" columns, and RefSeq protein accessions in the "prot" columns. The "paralogs" column indicates whether both members of the pair are inferred to be the products of post-transfer gene duplication from gene trees. The "cluster_number" column contains a unique numerical identifier assigned to 199 contiguous clusters of HTGs uninterrupted by non-HTGs; note that 42 clusters contain multiple adjacent pairs.

**Table S13. Clusters of adjacent HTGs predicted to catalyze consecutive biochemical reactions.** "Sequence" and "Enzyme" provide protein accessions and manual annotations for pairs of adjacent HTGs predicted to catalyze consecutive steps in the same biochemical "Pathway" as determined from MetaCyc[24]. "Ancestry" indicates the phylogenetic affinity of the sequence as in Table S18. "Enzyme" predictions for all proteins were verified with InterProScan (Table S18, sheet "manual_annotations").

| Pathway | Sequence 1 | Enzyme 1 | Ancestry 1 | Sequence 2 | Enzyme 2 | Ancestry 2 |
|---|---|---|---|---|---|---|
| Isopenicillin/β-lactam synthesis | XP_037037668.1 | N-(5-amino-5-carboxypentanoyl)-L-cysteinyl-D-valine synthase | Collembola | XP_037040229.1 | Isopenicillin-N synthase | Collembola |
| Pyridoxal 5′-phosphate biosynthesis | XP_037038870.1 | Pyridoxal 5'-phosphate synthase subunit PdxT/SNO | Fungi | XP_037038869.1 | Pyridoxal 5'-phosphate synthase subunit PdxS/SNZ | Bacteria |
| Glyoxylate shunt | XP_037048618.1 | Malate synthase | Collembola | XP_037048622.1 | Isocitrate lyase | Collembola |
| Kynurenine pathway | XP_037051528.1 | 3-hydroxyanthranilate 3,4-dioxygenase (HAAO) | Arachnida | XP_080118561.1 | 2-amino-3-carboxymuconate-6-semialdehyde decarboxylase (ACMSD) | Arachnida |

**Table S14 (attached file). HTG expression data,** in six sheets:

- **Datasets:** File with accession numbers of publicly available RNA-Seq accessions analyzed for tree-confirmed HTG expression and differential expression analysis. NCBI SRA accessions are in the "SRA run accession" column; "description" contains a description of the sex, developmental stage, and tissue; "Bioproject accession" NCBI identifier for submitting publication.
- **mean_tpms_per_stage_organ:** Mean transcripts per million (TPM) expression values per developmental stage, sex, and organ/tissue type for HTGs. "description" indicates the RefSeq description for the selected protein isoform, and "manual_annotation" indicates the function assigned via manual curation (if applicable; Table S18). Data are derived from previously published RNA-Seq data.
- DESeq2 differential expression data derived from publicly available RNA-Seq datasets for tree-confirmed, significantly upregulated (adjusted p-value < 0.05, log2 fold change > 0) HTGs. Data are split across two sheets: **irradiated_upregulated** for irradiated vs. non-irradiated larval comparisons; **larval_upregulated** for larval vs. non-larval samples. Columns correspond to standard DESeq2 output fields as follows: "gene" and "description" contain RefSeq data for the selected gene/protein; "baseMean" mean DESeq2-normalized counts across all samples; "log2FoldChange" log base 2 fold changes across each sheet's respective comparison (e.g., larval/non-larval for "larval_upregulated"); "lfcSE" standard error of the log2FoldChange; "stat" Wald test statistic; "pvalue" unadjusted Wald test p-value; "padj" Benjamini-Hochberg adjusted p-value; description indicates the RefSeq annotation description of the HTG; "manual_annotation" indicates the function annotation assigned by manual inspection of InterProScan results (Table S18, sheet "manual_annotations"). Columns with TRUE/FALSE entries indicate whether the gene was assigned to the biological pathway indicated in the column name via inspection of InterProScan results (Table S18, sheet "manual_annotations").
- **p.hygida_salivary_proteome:** HTGs with tree-determined homologs in *P. hygida*, with mass spectrometry proteomics expression derived from larval saliva as reported by Trinca and colleagues[27]. "P.hygida_protein" is the GenBank accession of the previously validated protein [27]; "B.copro_homologs" are tree-inferred homologous HTGs in *B. coprophila*; "Name" is the previously assigned [27] annotation; "Unique peptides" is the number of unique detected peptides from the *P. hygida* protein of interest. Columns with TRUE/FALSE entries indicate whether the gene was assigned to the biological pathway indicated in the column name via inspection of InterProScan results (Table S18, sheet "manual_annotations"). All data except homology and function inferences are from Table S10 of Trinca and colleagues 2023 [27].
- **Sanger_results**: Sanger sequencing results and primer information for RT-PCR of eight *B. coprophila* HTGs selected for experimental analysis (Figures 5, S3C-E, S4D-E, S7C-E, S11). Protein accessions of the sequences derived from the RefSeq annotation are available in the "RefSeq accession" column; sequencing products were submitted to GenBank with accessions in the "Product accession" column. For QPRT, the RT-PCR primers were initially constructed using an accession (XP_037047822.1) from the previous *B. coprophila* RefSeq genome annotation, which was updated to a new accession (XP_037047821.1) with identical amino acid sequence in the most recent RefSeq annotation.

**Table S15. Phylostratigraphic age assignments of *B. coprophila* HTGs and transfer events. "**Phylostratum" age categories are from youngest (1) to oldest (5). **"**Lower bound" age values in millions of years ago (MYA) were assigned according to the posterior mean divergence time of *B. coprophila* from the species in column "spanning species", which is the most distantly related species that contains tree-determined homolog(s) of each HTG; "upper bound" estimates are determined as the divergence time from the next most closely related sciarid species lacking homologs of each HTG, or the divergence time from Cecidomyiidae, the most closely related family to Sciaridae with RefSeq-annotated genomes, for phylostratum 5; "N_HTGs" is the number of HTGs that trace their origins to the phylostratum, including subsequent gene duplications; "N_transfers" is the number of inferred transfer events during the phylostratum.

| Phylostratum | Spanning species | Lower bound (MYA) | Upper bound (MYA) | N_HTGs | N_transfers |
|---|---|---|---|---|---|
| **1** | *B. coprophila* | 0 (present) | 14.1 | 53 | 45 |
| **2** | *B. odoriphaga* | 14.1 | 27.1 | 28 | 27 |
| **3** | *B. impatiens* | 27.1 | 58.2 | 106 | 79 |
| **4** | *L. ingenua* | 58.2 | 73.2 | 330 | 250 |
| **5** | *P. hygida* | 73.2 | 216.0 | 1,000 | 533 |

**Table S16 (attached file). Pairwise dN/dS estimates.** dN/dS statistics computed for reciprocal best DIAMOND BLASTp hits between *B. coprophila* and *B. impatiens.* Column descriptions are as follows: “B.coprophila” protein accession from *B. coprophila*; “B.impatiens” protein identifier for *B. impatiens* from BRAKER3 annotation (available on Dryad); “dNdS” PAML dN/dS (omega) maximum-likelihood estimate; “dS" synonymous substitution rate; “lnL” log-likelihood for the maximum-likelihood dN/dS estimate; “null_lnL”: log-likelihood for a null model with dN/dS fixed at 1; “2delta(lnL)” statistic used for likelihood ratio test for the null hypothesis that dN/dS=1; “p_val” p-value obtained from likelihood ratio test; “p_val_corrected” p-values adjusted via the Benjamini-Hochberg procedure; “type” is the ancestry annotation of the *B. coprophila* protein, with “Other” indicating genes with a “none” DIAMOND-based ancestry annotation (Table S6), or DIAMOND-inferred HTGs rejected by tree inspection. Only dN/dS values with corresponding dS < 2 were analyzed.

**Table S17. Ordinary least squares statistics for dN/dS** modeled as ~logTPM+Extracellular+HGT+CDS_length. logTPM = log10(TPM+$10^{-2}$). Extracellular = 1 if DeepLoc2 predicts an extracellular localization; HGT = 1 if HTG and 0 if dipteran. HGT remains significantly associated with dN/dS even when accounting for the other covariates. Bracketed values indicate bounds of the 95% confidence interval.

| **Test statistic** | **coef** | **std err** | **t-statistic** | **P-value** | **[0.025** | **0.975]** |
|---|---|---|---|---|---|---|
| Intercept | 0.136 | 0.002 | 69.627 | $<10^{-100}$ | 0.132 | 0.140 |
| logTPM | -0.018 | 0.001 | -16.755 | $2.82\times10^{-62}$ | -0.021 | -0.016 |
| Extracellular | 0.031 | 0.003 | 12.115 | $1.40\times10^{-33}$ | 0.026 | 0.036 |
| HGT | 0.036 | 0.003 | 11.324 | $1.42\times10^{-29}$ | 0.030 | 0.042 |
| CDS_len | $-1.83\times10^{-6}$ | $5.27\times10^{-7}$ | -3.476 | $5.11\times10^{-4}$ | $-2.86\times10^{-6}$ | $-7.98\times10^{-7}$ |
| R-squared | 0.060 | | | | | |
| Adj. R-squared | 0.059 | | | | | |
| F-statistic | 182.40 | | | | | |
| Prob (F-statistic) | $7.89\times10^{-152}$ | | | | | |
| Log-Likelihood | 10,902 | | | | | |
| AIC | $-2.18\times10^{4}$ | | | | | |
| BIC | $-2.18\times10^{4}$ | | | | | |

**Table S18 (attached file). Functional predictions for *B. coprophila* HTGs**, across the following nine sheets:

- **localization_predictions:** sheet contains DeepLoc2 subcellular localization predictions for HTGs, with the following columns: “Protein_ID” is the RefSeq protein accession; “Localizations” is the predicted label(s), separated with a “|” in the case of multiple predictions per protein; “Signals” are the predicted signal motif(s) if found; and the remaining columns are the localization probabilities per location.
- **localization_enrichment:** sheet contains enrichment statistics for DeepLoc2 predictions for HTGs, with “HGT” indicating the number of HTGs with each predicted localization; “FC” as the fold-change over non-HGT proteins; “p_value” the unadjusted p-value from Fisher’s exact test; “padj” the Benjamini-Hochberg adjusted p-value. The total number of proteins across localizations (1,853) exceeds the number of HTGs because DeepLoc2 assigned multiple predictions to 325 proteins.
- **molecular_function_enrichment:** data for Gene Ontology (“GO”) molecular functions, with the following columns: “GO term”; “Name” as the common name assigned to each GO term; “HTGs” gene accessions of HTGs; “HTGs_desc” with the annotations of all predicted HTGs with the predicted GO term; “HTG_count” with the number of HTGs with the GO term; “FC” as the fold-change over non-HGT proteins; “p_value” the unadjusted p-value from Fisher’s exact test; “padj” the Benjamini-Hochberg adjusted p-value.
- **biological_process_enrichment:** data for Gene Ontology (“GO”) biological processes, with the same columns as “molecular_function_enrichment”.
- **predicted_CAZymes:** tree-confirmed *B. coprophila* HTGs that were previously annotated as predicted carbohydrate-active enzymes (CAZymes)[27], with the following columns: “original_accession” is the *B. coprophila* HTG previously reported[27] accession from *B. coprophila* annotation GCF_014529535.1; “new_accession” is the equivalent accession obtained from the latest annotation GCF_014529535.2; “Final annotation” is the previously reported[27] manual CAZy family annotation (https://www.cazy.org/) with corresponding CAZy family code in “CAZy family”; “Pfam” is the automated Pfam domain(s); “EC Number” is the Enzyme Commission number of the predicted enzymatic reaction. All data except “new_accession” are from Table S9 of Trinca and colleagues’ report[27], with protein accessions mapped to the latest genome annotation as described in SI Methods.
- **confirmed_CAZymes:** *P. hygida* CAZymes validated via proteomics and enzyme assays performed against larval homogenates[27] that we infer to be orthologs of *B. coprophila* HTGs, with the following columns: “P.hygida_protein” is the GenBank accession of the previously validated enzyme[27]; “B.copro_homologs” are tree-inferred homologous HTGs in *B. coprophila* (RefSeq accession); “Annotation” is the confirmed enzymatic activity; “CAZy family” is the CAZy family code; “EC Number” is the Enzyme Commission number; “Substrate” is the carbohydrate substrate against which activity was validated (note: CMC and Avicel are cellulose derivatives); “Activity” is PCW for substrates found in plant cell walls, IE for insect endoskeleton, and FCW for fungal cell walls. All data except orthology inferences are from Table 1 of Trinca and colleagues, 2023[27].
- **InterProScan_counts:** numbers of HTGs and HTG accessions assigned to each InterProScan domain.
- **carbohydrate_catabolism:** a curated set of HTGs predicted to participate in the catabolism of carbohydrates, combining previously reported information[27] (see **predicted_CAZymes**) and manual annotations based on InterProScan results (see **manual_annotations**) as indicated in the “Source” column; “CAZy family” is the CAZy family code; “Cell wall substrate” indicates a plant, fungal or bacterial cell wall substrate

for the enzyme (if applicable); “Ancestry” indicates inferred HTG source via closest non-sciarid phylogenetic relative or top non-sciarid DIAMOND hit; “Localization” indicates DeepLoc-predicted subcellular localization. Rows highlighted in yellow are predicted to act on non-carbohydrate components of peptidoglycan. Rows with substrates in “Cell wall substrate” were used as inputs for Figure S9.

- **manual_annotations:** Functional predictions (“Manual annotation”) and predicted pathway membership (“Pathway(s)”) of select proteins discussed in this study. Data supporting functional inferences include: “RefSeq description” from current (GCF_014529535.2) and previous (GCF_014529535.1) RefSeq annotations “Domains”: InterProScan domain annotations with accessions in parentheses and delimited by semicolons; “Localization” is the DeepLoc2 predicted subcellular localization; “Source(s)” provides URLs for publications used for pathway assignments if applicable (blank if taken directly from InterPro domain descriptions); “Ancestry” reflects the likely origin of the HTG inferred from closest non-sciarid relatives on trees if possible, or from top non-sciarid DIAMOND hit if phylogenetic closest relative was polyphyletic.

**Table S19. Total numbers of HTGs assigned via manual annotation** to biological pathways (bold), their constituent sub-pathways (bold italicized) and gene-level functional annotations (normal text). Note that some gene-level functional annotations have been assigned to multiple biological pathways. Source data and RefSeq accessions for non-carbohydrate catabolism terms are provided in Table S18, sheet "manual_annotations"; carbohydrate catabolism data are derived from Table S18, sheet "carbohydrate_catabolism". Immune defense annotations are distributed across immune effectors and "Cell signaling-immune signaling/recognition" (cell signaling) annotations. CAZy family designations (https://www.cazy.org/) for carbohydrate-active enzymes are indicated in parentheses.

| Pathway or Annotation | N_HTGs |
|---|---|
| **Carbohydrate catabolism** | **126** |
| Endo-β-1,2-glucanase SGL (GH?) | 1 |
| Arylsulfatase H-like | 1 |
| Exo-β-1,4-glucuronan lyase (PL8_4) | 1 |
| Gluconokinase | 1 |
| Glucose-6-phosphatase | 1 |
| Glycoside hydrolase | 3 |
| Glycoside hydrolase family 31 (GH31) | 1 |
| Glycoside hydrolase family 65 (GH65) | 1 |
| Glycoside hydrolase, family 5 (GH5) | 1 |
| DegT/DnrJ/EryC1/StrS aminotransferase (GH5) | 1 |
| Glycosyl Hydrolase 5 (GH5) | 2 |
| Lactase (GH1) | 5 |
| Levanase/endo-inulinase (GH32) | 1 |
| O-Glycosyl hydrolase (GH30_1) | 1 |
| Unsaturated glucuronyl hydrolase (GH88) | 1 |
| Endo-α-1,4-polygalactosaminidase (GH114) | 1 |
| Exo-α-1,6-mannosidase (GH125) | 1 |
| α-L-fucosidase (GH29) | 1 |
| α-glucosidase (GH31) | 3 |
| β-galactosidase (GH35) | 3 |
| ***Carbohydrate catabolism-cellulose catabolism*** | ***15*** |
| Endoglucanase (GH45) | 7 |
| Endoglucanase (GH9) | 8 |
| ***Carbohydrate catabolism-chitin catabolism*** | ***19*** |
| Chitin deacetylase (CE4) | 1 |
| Chitinase (GH18) | 2 |
| Chitinase (GH19) | 6 |
| Endochitinase-like (GH?) | 1 |
| Lytic chitin monooxygenase (AA15) | 5 |
| N-acetyl-β-hexosaminidase (GH20) | 4 |
| ***Carbohydrate catabolism-mannan/hemicellulose catabolism*** | ***4*** |
| Endo-1,4-β-mannosidase (GH5_10) | 1 |
| β-mannosidase (GH130) | 1 |

| | |
|---|---|
| β-mannosidase (GH2) | 2 |
| ***Carbohydrate catabolism-pectin catabolism*** | ***14*** |
| Pectin methylesterase (CE8) | 2 |
| Pectin/pectate lyase (PL1_7) | 6 |
| Pectin/pectate lyase-like (PL?) | 1 |
| Polygalacturonase (GH28) | 5 |
| ***Carbohydrate catabolism-peptidoglycan catabolism*** | ***9*** |
| Glycosyl hydrolase YngK-like (Bacterial cell wall hydrolase) | 2 |
| Cell wall hydrolase SleB (Bacterial cell wall hydrolase) | 1 |
| Domesticated amidase effector/Type VI secretion system amidase effector | 1 |
| Lysozyme (GH24) | 2 |
| Murein tetrapeptide carboxypeptidase | 2 |
| N-acetylmuramic acid 6-phosphate etherase MurQ | 1 |
| ***Carbohydrate catabolism-xylan/hemicellulose catabolism*** | ***12*** |
| Endo-1,4-β-xylanase (GH11) | 2 |
| Endo-1,4-β-xylanase (GH43_34) | 1 |
| Exo-α-(1-5)-L-arabinofuranosidase (GH43_26) | 2 |
| Exoglucanase/xylanase (GH10) | 1 |
| Xylan exo-α-(1-2)-glucuronosidase (GH67) | 1 |
| α-L-arabinofuranosidase (GH51) | 2 |
| β-D-xylosidase (GH3) | 3 |
| ***Carbohydrate catabolism-α-1,3-glucan*** | ***2*** |
| Glucan-endo-1,3-α-glucosidase (GH71) | 2 |
| ***Carbohydrate catabolism-β-1,3-glucan*** | ***7*** |
| Glucan-1,3-β-glucosidase (GH5) | 1 |
| Glucan-1,3-β-glucosidase (GH55) | 1 |
| Exo-β-1,3-glucanase (GH5_9) | 1 |
| β-1,3-glucanase (GH17) | 3 |
| Endo-β-1,3-glucanase (GH16) | 1 |
| ***Carbohydrate catabolism-monosaccharide import*** | ***13*** |
| L-fucose-proton symporter | 1 |
| Sodium-dependent glucose transporter (SGLT) | 12 |
| **Protein catabolism** | **81** |
| Acylamino-acid-releasing enzyme | 2 |
| Arylamine N-acetyltransferase; Cysteine peptidase | 3 |
| Calpain peptidase | 1 |
| Carboxypeptidase Q | 2 |
| Carboxypeptidase Q-like | 1 |
| Caspase | 3 |
| Chymotrypsin-1-like | 2 |
| Chymotrypsin-2-like | 1 |
| Cyanophycinase-like | 1 |
| Cytosolic non-specific dipeptidase | 1 |
| Metalloendopeptidase OMA1, mitochondrial-like | 1 |
| Metallopeptidase | 15 |

| | |
|---|---|
| Neurolysin, mitochondrial-like | 1 |
| PPPDE peptidase | 1 |
| Peptidase | 1 |
| Peptidase C45-like | 1 |
| Peptidase M50B-like | 1 |
| Peptidase M64 | 1 |
| Prenylcysteine oxidase | 1 |
| Proline iminopeptidase | 5 |
| Prolyl endopeptidase | 2 |
| Rhomboid peptidase | 7 |
| Secernin-3-like | 1 |
| Serine β-lactamase-like protein LACTB, mitochondrial | 7 |
| Serine protease | 1 |
| Subtilisin protease | 13 |
| TRPM8 channel-associated factor 3-like/Peptidase M60 | 1 |
| Thimet oligopeptidase-like | 1 |
| Trypsin-1-like | 1 |
| Venom serine carboxypeptidase-like | 2 |
| **Amino acid catabolism** | **24** |
| Aspartate dehydrogenase | 1 |
| 2-iminobutanoate/2-iminopropanoate deaminase | 1 |
| 4-hydroxy-2-oxoglutarate aldolase, mitochondrial-like | 1 |
| Argininosuccinate lyase | 1 |
| Aspartate/prephenate aminotransferase | 1 |
| Cysteine desulfurase | 1 |
| D-amino-acid oxidase | 2 |
| D-aminoacylase | 1 |
| Fumarylacetoacetase | 1 |
| Histidine ammonia-lyase | 2 |
| L-serine dehydratase/L-threonine deaminase | 7 |
| Prenylcysteine oxidase | 1 |
| Saccharopine dehydrogenase | 2 |
| Selenocysteine lyase | 1 |
| Tryptophanase | 1 |
| **Lipid catabolism** | **67** |
| 2,4-dienoyl-CoA reductase | 1 |
| 2-hydroxy-palmitic acid dioxygenase Mpo1-like | 1 |
| 3-hydroxybutyrate dehydrogenase | 1 |
| Acyl-CoA dehydrogenase | 1 |
| Acyl-CoA dehydrogenase family member 11-like | 1 |
| Acyl-coenzyme A synthetase ACSM3 | 1 |
| Alkaline ceramidase | 1 |
| Amidase signature (AS) superfamily/Vitamin D3 hydroxylase-associated protein-like | 1 |
| Amidase signature (AS) superfamily/monoglyceride lipase faah | 1 |
| Arylsulfatase H-like | 1 |

| | |
|---|---|
| CD36 family (scavenger receptor) | 1 |
| Carboxyl esterase | 3 |
| D-β-hydroxybutyrate dehydrogenase, mitochondrial-like | 1 |
| Enoyl-CoA hydratase | 2 |
| Enoyl-CoA hydratase echA8 | 1 |
| Fungal lipase-like | 9 |
| GDXG lipase | 3 |
| Glycerophosphodiester phosphodiesterase | 1 |
| Glycoside hydrolase family 30/lysosomal acid glucosylceramidase | 1 |
| Lipase/calcium-activated chloride channel regulator | 2 |
| Low-density lipoprotein (LDL) receptor | 1 |
| Lysophospholipase D GDPD1-like | 1 |
| Lysoplasmalogenase | 1 |
| Mitochondrial ABHD10 acyl-protein thioesterase | 1 |
| Mitochondrial carnitine/acylcarnitine carrier protein | 1 |
| Mitochondrial coenzyme A diphosphatase | 1 |
| N-acyl-phosphatidylethanolamine-hydrolyzing phospholipase D | 1 |
| PLC-like phosphodiesterase | 1 |
| Peroxisomal trans-2-enoyl-CoA reductase | 2 |
| Phosphatidyl-N-methylethanolamine/Phosphatidylethanolamine N-methyltransferase | 1 |
| Phospholipase A and acyltransferase | 5 |
| Phospholipase A and acyltransferase 1-like | 1 |
| Phospholipase B | 1 |
| Phytanoyl-CoA dioxygenase | 6 |
| Prenylcysteine oxidase | 1 |
| SGNH hydrolase | 7 |
| Sterol transport protein NPC2-like | 1 |
| **Glyoxylate shunt** | **2** |
| Isocitrate lyase | 1 |
| Malate synthase | 1 |
| **Proteostasis** | **58** |
| Acylamino-acid-releasing enzyme | 2 |
| Cystatin-A2 | 2 |
| DSBA-like thioredoxin | 1 |
| F-box associated (FBA) domain protein | 2 |
| F-box domain protein | 7 |
| LanC-like protein | 1 |
| LanC-like protein, eukaryotic | 3 |
| Metalloendopeptidase OMA1, mitochondrial-like | 1 |
| Protein IMPACT homolog | 1 |
| Protein mono-ADP-ribosyltransferase PARP16-like | 2 |
| Serine/threonine-protein kinase/endoribonuclease IRE1 | 2 |
| Serine/threonine-protein kinase/endoribonuclease IRE1/2-like | 1 |
| Speckle-type POZ protein-like | 2 |
| Ubiquitin carboxyl-terminal hydrolase | 1 |

| | |
|---|---|
| Ubiquitin-conjugating enzyme E2-like | 3 |
| ***Proteostasis-E3 ubiquitin-protein ligase*** | ***16*** |
| IBR-containing E3 ligase | 1 |
| RBR-family/RNF14-like E3 ubiquitin-protein ligase | 1 |
| RING-SPRY/B30.2-domain protein; putative E3 ubiquitin ligase | 4 |
| RNF-type E3 ubiquitin-protein ligase | 1 |
| E3 ligase/Deltex family protein | 4 |
| E3 ubiquitin-protein ligase NRDP1-like | 1 |
| E3 ubiquitin-protein ligase RNF14-like | 1 |
| E3 ubiquitin-protein ligase SHPRH-like | 1 |
| E3 ubiquitin-protein ligase SINA | 1 |
| E3 ubiquitin-protein ligase parkin-like | 1 |
| ***Proteostasis-chaperones*** | ***11*** |
| Mitochondrial chaperone BCS1-like | 1 |
| Cyclophilin-like (chaperone) | 1 |
| Hsp70 chaperone | 1 |
| DNAJ domain protein (chaperone) | 5 |
| Peptidyl-prolyl cis-trans isomerase FKBP-like (chaperone) | 3 |
| **Transmembrane transport** | **68** |
| ABC transporter G family | 2 |
| ATP-binding cassette subfamily G transporter | 1 |
| Acetyl-coenzyme A transporter 1-like | 1 |
| Ammonium transporter | 2 |
| Aquaporin | 1 |
| Bacterial-type extracellular solute-binding protein | 1 |
| CD36 family (scavenger receptor) | 1 |
| Choline/ethanolamine transporter | 1 |
| Citrulline exporter C365.16-like | 1 |
| Failed axon connections homolog | 2 |
| Ionotropic receptor | 1 |
| L-fucose-proton symporter | 1 |
| Large-conductance mechanosensitive channel | 2 |
| Lipase/calcium-activated chloride channel regulator | 2 |
| Low-density lipoprotein (LDL) receptor | 1 |
| MFS transporter | 1 |
| MFS transporter/putative peptidoglycan muropeptide transporter SLC46 | 3 |
| Magnesium-transporting ATPase, P-type | 1 |
| Mg2+ transporter protein, CorA-like/Zinc transport protein | 1 |
| Mitochondrial carnitine/acylcarnitine carrier protein | 1 |
| Mitochondrial substrate carrier | 1 |
| Multi Antimicrobial Extrusion Protein | 1 |
| Multidrug transporter EmrE superfamily | 1 |
| Na(+)/citrate cotransporter | 1 |
| Na(+)/citrate cotransporter-like | 1 |
| Nucleobase cation symporter | 1 |

| | |
|---|---|
| Probable citrulline exporter | 1 |
| Protein translocase subunit SecA | 1 |
| Proton-linked Monocarboxylate Transporter/MFS transporter | 5 |
| Putative ABC-transporter type IV | 1 |
| SecA domain protein | 1 |
| Sodium-coupled monocarboxylate transporter | 4 |
| Sodium-dependent glucose transporter | 12 |
| Sodium-dependent multivitamin transporter | 1 |
| Sodium/anion cotransporter | 1 |
| Sodium/potassium-transporting ATPase subunit alpha-1-like | 1 |
| Sodium:solute symporter (SSF) | 1 |
| Solute carrier family 23 member | 1 |
| Sterol transport protein NPC2-like | 1 |
| UNC93-like protein | 2 |
| Voltage-gated hydrogen channel 1 | 1 |
| Voltage-gated hydrogen channel 1-like | 1 |
| **ADP-ribosylation** | **9** |
| ADP-ribosyltransferase | 2 |
| Mono-ADP-ribosyltransferase | 1 |
| Mono-ADP-ribosyltransferase and antiviral protein | 2 |
| NAD(P)(+)--arginine ADP-ribosyltransferase | 1 |
| Protein mono-ADP-ribosyltransferase PARP16-like | 2 |
| poly [ADP-ribose] polymerase (PARP) tankyrase-like | 1 |
| **Bacterial cell membrane biosynthesis** | **9** |
| Cyclopropane-fatty-acyl-phospholipid synthase | 8 |
| GDP-D-glycero-alpha-D-manno-heptose dehydrogenase-like | 1 |
| **Bacterial cell wall biosynthesis** | **15** |
| Alanine racemase | 1 |
| Alanine racemase/D-amino acid metabolism enzyme | 1 |
| Anhydro-N-acetylmuramic acid kinase | 1 |
| D-alanine--D-alanine ligase | 4 |
| Diaminopimelate epimerase-like | 1 |
| N-acetylmuramate/N-acetylglucosamine kinase | 1 |
| β-lactamase/transpeptidase | 6 |
| **Bacterial cell wall degradation/turnover** | **13** |
| Anhydro-N-acetylmuramic acid kinase | 1 |
| Cell wall hydrolase SleB | 1 |
| D-amino-acid oxidase | 2 |
| Domesticated amidase effector | 1 |
| Glycosyl hydrolase YngK-like | 2 |
| Lysozyme | 2 |
| Murein tetrapeptide carboxypeptidase | 2 |
| N-acetylmuramate/N-acetylglucosamine kinase | 1 |
| N-acetylmuramic acid 6-phosphate etherase MurQ | 1 |
| **Protein toxins** | **12** |

| | |
|---|---|
| Aerolysin-like toxin | 3 |
| Cytolytic pore-forming toxin | 5 |
| Hemolysin E (pore-forming toxin) | 1 |
| TcdA/TcdB toxin | 1 |
| Tox-ART-HYD1 | 2 |
| **Immune effectors** | **14** |
| Defensin | 2 |
| Domesticated amidase effector | 1 |
| Gamma-interferon-inducible lysosomal thiol reductase | 1 |
| Interferon-induced very large GTPase | 1 |
| Lysozyme | 2 |
| Membrane attack complex component/perforin (MACPF) domain protein | 2 |
| Mono-ADP-ribosyltransferase and antiviral protein | 2 |
| Pathogen-related defense protein | 2 |
| Viperin | 1 |
| **Cell signaling** | **63** |
| Arrestin domain (GPCR signaling) | 3 |
| Calmodulin-alpha | 1 |
| Caspase | 3 |
| Differentiation-associated signaling protein | 1 |
| Dual specificity protein phosphatase | 2 |
| E3 ligase/Deltex family protein | 4 |
| Ecdysteroid kinase-like | 2 |
| Ephexin | 1 |
| Integrin domain protein | 3 |
| Peptidyl-prolyl cis-trans isomerase FKBP-like (chaperone) | 2 |
| Platelet-derived growth factor receptor-like | 1 |
| Receptor tyrosine kinase | 2 |
| Serine/threonine-protein kinase | 4 |
| Serine/threonine-protein kinase/endoribonuclease IRE1 | 2 |
| Serine/threonine-protein kinase/endoribonuclease IRE1/2-like | 1 |
| Serine/threonine-protein phosphatase | 2 |
| Store-operated calcium entry-associated regulatory factor | 1 |
| Sushi domain-containing protein 2-like | 5 |
| ***Cell signaling-immune signaling/recognition*** | ***9*** |
| 2-5-oligoadenylate synthetase 1 (OAS1) | 1 |
| MFS transporter/putative peptidoglycan muropeptide transporter SLC46 | 3 |
| Macrophage migration inhibitory factor-like | 1 |
| NFX1-type zinc finger-containing protein 1 (RNA sensing) | 1 |
| Pfs47 receptor/natterin-like | 2 |
| Tumor Necrosis Factor Receptor-Associated Factor (TRAF) domain | 1 |
| ***Cell signaling-neuronal signaling*** | ***14*** |
| (Dopamine β-hydroxylase) DBH-like monooxygenase protein 1 homolog | 3 |
| Acetylcholine receptor subunit | 3 |
| Acetylcholinesterase | 2 |

| | |
|---|---|
| Amine oxidase | 1 |
| Aromatic-L-amino-acid decarboxylase | 1 |
| Glutamate decarboxylase | 1 |
| Ionotropic receptor | 1 |
| UNC93-like protein | 2 |
| **DNA repair** | **16** |
| ATP-dependent DNA helicase PcrA | 1 |
| Argonaute 2 | 1 |
| Cryptochrome DASH/pyrimidine photolyase | 1 |
| DNA polymerase kappa | 1 |
| E3 ubiquitin-protein ligase SHPRH-like | 1 |
| Fanconi-associated nuclease | 1 |
| Replication protein A | 1 |
| Smr domain protein | 1 |
| Tyrosyl-DNA phosphodiesterase | 1 |
| poly [ADP-ribose] polymerase (PARP) tankyrase-like | 1 |
| ***DNA repair-Base excision or damage reversal*** | ***6*** |
| Methylpurine DNA glycosylase (MPG) | 1 |
| Uracil-DNA glycosylase (UDG) | 1 |
| AP endonuclease family 1 member (APEX1) | 1 |
| AP endonuclease family 2 member (APEX2) | 1 |
| Oxidized purine nucleoside triphosphate hydrolase (8ODP) | 1 |
| DNA oxidative demethylase ALKBH2 | 1 |
| **RNA repair** | **2** |
| RNA ligase domain, REL/Rln2 protein | 1 |
| Swiss Army Knife | 1 |
| **Xenobiotic metabolism/toxin resistance** | **115** |
| 2,6-Dihydroxypyridine 3-monooxygenase | 1 |
| 2-iminobutanoate/2-iminopropanoate deaminase | 1 |
| Aldehyde dehydrogenase | 1 |
| Amine oxidase | 1 |
| Arsenite methyltransferase (ArsM) | 1 |
| Arylamine N-acetyltransferase; Cysteine peptidase | 3 |
| Dehydrogenase/reductase SDR family member 7 | 2 |
| L-2-Haloacid dehalogenase | 1 |
| Methanethiol oxidase | 2 |
| NADH:cytochrome b5 reductase | 1 |
| Nitronate monooxygenase | 1 |
| Oxalate--CoA ligase Pcs60 | 1 |
| Renalase | 1 |
| ***Xenobiotic metabolism/toxin resistance-cyanide resistance*** | ***5*** |
| Alternative oxidase | 1 |
| B-cyanoalanine synthase | 1 |
| Bifunctional nitrilase/nitrile hydratase NIT4A | 2 |
| Cyanate hydratase | 1 |

| | |
|---|---|
| ***Xenobiotic metabolism/toxin resistance-Phase I xenobiotic metabolism*** | ***60*** |
| Alpha-ketoglutarate-dependent 2,4-dichlorophenoxyacetate dioxygenase (TfdA) | 1 |
| Trans-1,2-dihydrobenzene-1,2-diol dehydrogenase-like | 2 |
| Aldo/keto reductase | 5 |
| Carboxyl esterase | 3 |
| Bifunctional epoxide hydrolase 2-like | 1 |
| Extradiol ring-cleavage dioxygenase | 1 |
| Intradiol ring-cleavage dioxygenase | 1 |
| Mitochondrial amidoxime-reducing component 2-like | 1 |
| Mitochondrial amidoxime-reducing component 1-like | 1 |
| Cytochrome P450 | 27 |
| L-xylulose reductase/carbonyl reductase | 3 |
| Epoxide hydrolase | 1 |
| Carbonyl reductase | 1 |
| Flavin monooxygenase FMO | 12 |
| ***Xenobiotic metabolism/toxin resistance-Phase II xenobiotic metabolism*** | ***30*** |
| Sulfotransferase | 1 |
| Thiol S-methyltransferase | 6 |
| UDP-glucosyltransferase | 6 |
| Glutathione S-transferase | 17 |
| ***Xenobiotic metabolism/toxin resistance-Phase III xenobiotic metabolism*** | ***3*** |
| ATP-binding cassette subfamily G transporter | 1 |
| Multidrug transporter EmrE superfamily | 1 |
| Multi antimicrobial extrusion protein | 1 |
| **Redox homeostasis/oxidative stress response** | **67** |
| Acylamino-acid-releasing enzyme | 2 |
| Catalase | 11 |
| Cytochrome b561/ferric reductase | 1 |
| Delta(24)-sterol reductase | 1 |
| Esterase FE4-like | 1 |
| Haem oxygenase | 1 |
| Metalloendopeptidase OMA1, mitochondrial-like | 1 |
| Mitochondrial amidoxime reducing component 1-like | 1 |
| Mitochondrial amidoxime reducing component 2-like | 1 |
| Nitronate monooxygenase | 1 |
| Oxidized purine nucleoside triphosphate hydrolase (8ODP) | 1 |
| Oxidoreductase Oye-like | 3 |
| Peroxidase | 5 |
| Thioredoxin | 1 |
| DSBA-like thioredoxin | 1 |
| Peroxiredoxin/Thioredoxin-like | 1 |
| Peroxynitrite isomerase THAP4-like | 1 |
| Pirin | 1 |
| Pyridine nucleotide-disulfide oxidoreductase/amino oxidase | 1 |
| Sirtuin (mitochondrial) | 1 |

| | |
|---|---|
| Gamma-interferon-inducible lysosomal thiol reductase | 1 |
| ***Redox homeostasis/oxidative stress response-glutathione metabolism*** | ***27*** |
| Gamma-glutamylcyclotransferase | 1 |
| Glutamate--cysteine ligase | 1 |
| Glutathione S-transferase | 17 |
| Glutathione hydrolase | 3 |
| Glutathione reductase | 1 |
| LanC-like protein | 1 |
| LanC-like protein, eukaryotic | 3 |
| ***Redox homeostasis/oxidative stress response-alternative respiratory chain protein*** | ***2*** |
| Alternative NADH dehydrogenase | 1 |
| Alternative oxidase | 1 |
| **Epigenetic regulation** | **27** |
| 5-hydroxymethylcytosine (5hmC) reader, thymocyte nuclear protein 1-like | 1 |
| Acyl-CoA N-acyltransferase with predicted nuclear localization | 4 |
| DNA oxidative demethylase ALKBH2 | 1 |
| JmjC domain histone demethylation protein | 1 |
| Myb/SANT-like DNA-binding domain 5 | 1 |
| SET and MYND domain-containing protein | 4 |
| SET domain-containing protein | 1 |
| SET domain-containing protein SmydA-8 | 3 |
| SWR1-complex protein 5/Craniofacial development protein 1/2 | 2 |
| Chromo domain protein | 1 |
| Zinc finger (nuclear) | 4 |
| ***Epigenetic regulation-Transposon suppression*** | ***4*** |
| Endoribonuclease Dicer | 1 |
| Argonaute 2 | 1 |
| Histone-lysine N-methyltransferase SUV39H2-like | 2 |
| **Antioxidant/cofactor biosynthesis** | 34 |
| ***Antioxidant/cofactor biosynthesis-ergothioneine*** | ***4*** |
| Ergothioneine synthesis protein 1 (histidine-specific methyltransferase) | 1 |
| Ergothioneine synthesis protein 2 (hercynylcysteine sulfoxide lyase) | 3 |
| ***Antioxidant/cofactor biosynthesis-Vitamin C*** | ***21*** |
| L-gulono-1,4-lactone oxidase | 17 |
| Regucalcin | 4 |
| ***Antioxidant/cofactor biosynthesis pyridoxal 5-phosphate*** | ***2*** |
| Pyridoxal 5’-phosphate synthase subunit PdxS/SNZ | 1 |
| Pyridoxal 5’-phosphate synthase subunit PdxT/SNO | 1 |
| ***Antioxidant/cofactor biosynthesis-kynurenine pathway (NAD$^+$)*** | ***7*** |
| 2-amino-3-carboxymuconate-6-semialdehyde decarboxylase (ACMSD) | 2 |
| 3-hydroxyanthranilate 3,4-dioxygenase (HAAO) | 1 |
| Indoleamine 2,3-dioxygenase | 1 |
| Kynureninase | 1 |
| Kynurenine formamidase | 1 |
| Nicotinate-nucleotide pyrophosphorylase/quinolinate phosphoribosyl transferase (QPRT) | 1 |

| | |
|---|---|
| **Amino acid biosynthesis** | **13** |
| Asparagine synthetase | 1 |
| Aspartate/prephenate aminotransferase | 1 |
| B-cyanoalanine synthase | 1 |
| Chorismate mutase | 1 |
| Cobalamin-independent methionine synthase MetE, C-terminal/archaeal | 1 |
| Diaminopimelate epimerase-like | 1 |
| Glutamine synthetase | 1 |
| Homoserine/serine acetyltransferase | 1 |
| Threonine synthase | 1 |
| Argininosuccinate lyase | 1 |
| ***Amino acid biosynthesis-D-amino acids*** | ***3*** |
| Serine racemase-like | 1 |
| Alanine racemase | 1 |
| Alanine racemase/D-amino acid metabolism enzyme | 1 |
| **Antimicrobial/secondary metabolite biosynthesis** | **56** |
| Anthrone oxygenase | 1 |
| Baeyer-Villiger monooxygenase-like | 4 |
| Berberine/berberine-like biosynthetic oxidoreductase | 18 |
| Biosynthetic oxidoreductase | 2 |
| Fungal secondary metabolite SDR | 1 |
| Glycoside hydrolase, family 5;DegT/DnrJ/EryC1/StrS aminotransferase | 1 |
| Hydroxylase/desaturase AsaB-like | 2 |
| Isoprenoid/terpene synthase | 6 |
| Phenazine biosynthesis-like domain-containing protein | 3 |
| Questin oxidase-like | 1 |
| Secondary metabolite biosynthesis oxidoreductase | 1 |
| Strictosidine synthase | 4 |
| ***Polyketide/non-ribosomal peptide synthesis-associated enzymes*** | ***12*** |
| Polyketide cyclase SnoaL-like | 6 |
| Polyketide synthase acyl carrier protein | 2 |
| Polyketide synthase-like | 1 |
| Non-ribosomal peptide synthetase-like phosphopantetheine binding ACP domain-containing protein | 1 |
| N-(5-amino-5-carboxypentanoyl)-L-cysteinyl-D-valine synthase | 1 |
| Isopenicillin-N synthase | 1 |
| **Lipid biosynthesis/modification** | **92** |
| 17-β-hydroxysteroid dehydrogenase 13-like | 1 |
| 3-oxo-5-alpha-steroid 4-dehydrogenase | 3 |
| 3-oxoacyl-[acyl-carrier-protein] reductase FabG | 1 |
| Acyl carrier protein | 1 |
| Carotenoid oxygenase | 1 |
| Cholesterol 7-desaturase | 1 |
| Choline kinase alpha-like | 13 |
| Cyclopropane-fatty-acyl-phospholipid synthase | 8 |

| | |
|---|---|
| Dehydrogenase/reductase SDR family member 7 | 2 |
| Dehydrogenase/reductase SDR family member 9-like | 1 |
| Delta(24)-sterol reductase | 1 |
| Ecdysteroid kinase-like | 2 |
| Fatty acid desaturase | 24 |
| Fatty acid desaturase/Acyl-lipid (8-3)-desaturase B-like | 1 |
| Fatty acid desaturase/Delta(5) fatty acid desaturase A-like | 1 |
| Fatty acid hydroxylase/Sterol desaturase | 2 |
| Isoprenoid/terpene synthase | 6 |
| NADH:cytochrome b5 reductase | 1 |
| O-acyltransferase WSD1 | 1 |
| Peroxisomal trans-2-enoyl-CoA reductase | 2 |
| Phosphatidyl-N-methylethanolamine/Phosphatidylethanolamine N-methyltransferase | 1 |
| Phospholipase A and acyltransferase | 5 |
| Phospholipase A and acyltransferase 1-like | 1 |
| Prostaglandin reductase 1-like | 1 |
| Prostaglandin reductase 3 | 1 |
| Putative fatty acyl-CoA reductase | 1 |
| Retinol dehydrogenase | 6 |
| Sphingomyelin synthase-like | 1 |
| Sterol biosynthesis ERG24/DHCR-like | 1 |
| Very long chain fatty acid elongase 6 | 1 |

**Table S20. Selected HTGs absent or rare in non-sciarid dipterans.** Table contains evidence of the absence of HTG gene families in non-sciarid dipterans to support inferences of novel biochemical functionality conferred by HTGs. Analysis restricted to HTGs highlighted in text and figures. "Absence rank" indicates the highest taxonomic rank at or above Diptera (excluding Sciaridae) at which gene absence was inferred from the UniProt taxonomy browser in InterPro (https://www.ebi.ac.uk/interpro/entry/InterPro/, access date 7/21/2026) for the domain accession in "InterPro entry", with exceptions (possibly due to genomic contamination or independent HGT events) noted in parentheses. Evidence for the absence of β-cyanoalanine synthase from non-sciarid dipterans is provided in Figure S4D.

| **Accession** | **Name** | **Pathway** | **InterPro entry** | **Absence rank** |
|---|---|---|---|---|
| XP_037049870.1 | Oxidized purine nucleoside triphosphate hydrolase (8ODP) | DNA repair | IPR003563 | Diptera |
| XP_037040393.1 | Uracil-DNA glycosylase (UDG) | DNA repair | IPR002043 | Diptera |
| XP_037036837.1 | Methylpurine DNA glycosylase (MPG) | DNA repair | IPR003180 | Diptera |
| XP_037042473.1 | Apurinic or apyrimidinic site endonuclease family 2 (APEX2) | DNA repair | IPR001719 | Insecta (except *Lasius niger*, *Aphis craccivora*, *Zophobas morio*) |
| XP_037035483.1 | $NAD^+$ diphosphatase | $NAD^+$ metabolism | IPR049734 | Endopterygota (except *Anopheles maculatus*) |
| XP_037041147.1 | Kynureninase | Kynurenine pathway; $NAD^+$ biosynthesis | IPR010111 | Insecta (except *Ladona fulva*, Hemiptera, *Polypedilum vanderplanki*, and a previously noted HGT to Lepidoptera[196]) |
| XP_037051528.1 | 3-hydroxyanthranilate 3,4-dioxygenase (HAAO) | Kynurenine pathway; $NAD^+$ biosynthesis | IPR010329 | Insecta (except *Lucilia cuprina*) |
| XP_037047821.1 | Nicotinate-nucleotide pyrophosphorylase/quinolinate phosphoribosyl transferase (QPRT) | Kynurenine pathway; $NAD^+$ biosynthesis | IPR002638 | Insecta (except *Glossina* spp. and *Dendroctonus ponderosae*) |
| XP_037047040.1 | 2-amino-3-carboxymuconate-6-semialdehyde decarboxylase (ACMSD) | Kynurenine pathway; $NAD^+$ biosynthesis | IPR032465 | Insecta |
| XP_080118561.1 | 2-amino-3-carboxymuconate-6-semialdehyde decarboxylase (ACMSD) | Kynurenine pathway; $NAD^+$ biosynthesis | IPR032465 | Insecta (except *Photinus pyralis*) |
| XP_037029494.1 | Tryptophanase/tryptophan-indole lyase (TNase) | Tryptophan catabolism | IPR018176 | Metazoa (except *Trichuris trichiura*, *Leptidea sinapis*, *Cyprideis torosa*, *Darwinula stevensoni*, *Limulus polyphemus*) |
| XP_080117808.1 (and 16 others) | L-gulono-1,4-lactone oxidase | Vitamin C biosynthesis | IPR010031 | Insecta (except for *Gryllus longicercus*, *Eumeta variegata*, *Cloeon dipterum,* and Blattodea) |
| XP_037049532.1 | Ergothioneine synthesis protein 1 (histidine-specific methyltransferase) | Ergothioneine biosynthesis | IPR051128 | Arthropoda (except for *Anopheles melas*, *Cinara cedri*, and *Bemisia tabaci*) |

| | | | | |
|---|---|---|---|---|
| XP_037038870.1 | Pyridoxal 5'-phosphate synthase subunit PdxT/SNO | Pyridoxal 5'-phosphate synthesis | IPR002161 | Bilateria, except for Spiralia, plant-parasitic nematodes[73], Copepoda, three chordates, and *Lasius niger*. |
| XP_037038869.1 | Pyridoxal 5'-phosphate synthase subunit PdxS/SNZ | Pyridoxal 5'-phosphate synthesis | IPR001852 | Bilateria, except for Spiralia, a previously characterized HTG in plant-parasitic nematodes[73], Copepoda, three chordates, *Lasius niger* and *Hirondellea gigas* |
| XP_037042500.1 | Arsenite methyltransferase | Arsenite detoxification | IPR026669 | Diptera |

**Table S21. Selected dipteran genes and HTGs with predicted function in tryptophan catabolism and de novo NAD$^+$ synthesis.** Ancestry for putative HTGs was inferred directly from closest relatives when tree topologies unambiguously indicated a single HGT donor, or else from the top non-sciarid DIAMOND hit. “EC number” reports the Enzyme Commission number for the predicted reaction using RefSeq assigned annotations alongside InterProScan evidence presented in Table S18.

| Accession | Annotation | Predicted function | EC number | Ancestry |
|---|---|---|---|---|
| XP_037029494.1 | Tryptophanase/tryptophan-indole lyase (TNase) | Tryptophan catabolism (non-kynurenine pathway) | 4.1.99.1 | Bacteria |
| XP_037030629.1 | Indoleamine 2,3-dioxygenase (IDO) | Kynurenine pathway (catabolic portion) | 1.13.11.11/1.13.11.52 | Fungi |
| XP_037038643.1 | Tryptophan 2,3-dioxygenase (TDO) | Kynurenine pathway (catabolic portion) | 1.13.11.11/1.13.11.52 | Diptera |
| XP_037038012.2 | Kynurenine formamidase (KFase) | Kynurenine pathway (catabolic portion) | 3.5.1.9 | Diptera |
| XP_037051522.1 | Kynurenine formamidase (KFase) | Kynurenine pathway (catabolic portion) | 3.5.1.9 | Diptera |
| XP_037041193.1 | Kynurenine formamidase (KFase) | Kynurenine pathway (catabolic portion) | 3.5.1.9 | Collembola |
| XP_080113447.1 | Kynurenine 3-monooxygenase (KMO) | Kynurenine pathway (catabolic portion) | 1.14.13.9 | Diptera |
| XP_037041147.1 | Kynureninase (KYNU) | Kynurenine pathway (de novo NAD+ synthesis) | 3.7.1.3 | Collembola |
| XP_037051528.1 | 3-hydroxyanthranilate 3,4-dioxygenase (HAAO) | Kynurenine pathway (de novo NAD+ synthesis) | 1.13.11.6 | Arachnida |
| XP_037047821.1 | Nicotinate-nucleotide pyrophosphorylase/quinolinate phosphoribosyl transferase (QPRT) | Kynurenine pathway (de novo NAD+ synthesis) | 2.4.2.19 | Arachnida |
| XP_037029936.1 | Nicotinamide/nicotinic acid mononucleotide adenylyltransferase (NMNAT) | De novo NAD+ synthesis/NAD+ salvage pathway | 2.7.7.18 | Diptera |
| XP_037028206.1 | Glutamine-dependent NAD$^+$ synthetase (NADsyn) | De novo NAD+ synthesis/NAD+ salvage pathway | 6.3.5.1 | Diptera |
| XP_037035483.1 | NAD$^+$ diphosphatase (NADPase) | NAD+ salvage pathway/NAD+ degradation | 3.6.1.22 | Collembola |
| XP_037047040.1 | 2-amino-3-carboxymuconate-6-semialdehyde decarboxylase (ACMSD) | Kynurenine pathway shunt; reduces the concentrations of the toxic intermediate quinolinate | 4.1.1.45 | Collembola |
| XP_080118561.1 | 2-amino-3-carboxymuconate-6-semialdehyde decarboxylase (ACMSD) | Kynurenine pathway shunt; reduces the concentrations of the toxic intermediate quinolinate | 4.1.1.45 | Arachnida |

**Table S22. Selected dipteran genes and HTGs with predicted function in biosynthetic pathways.** Ancestry for putative HTGs was inferred directly from closest relatives when tree topologies unambiguously indicated a single HGT donor, or else from the top non-sciarid DIAMOND hit. Note: XP_037037356.2, indicated with an asterisk, returned a top non-sciarid DIAMOND hit in Hymenoptera but did not meet our alien index threshold for HTG inference. Annotations were assigned using RefSeq descriptions (when available) alongside InterProScan evidence presented in Table S18 (sheet "manual_annotations").

| Accession | Annotation | Pathway | Ancestry |
|---|---|---|---|
| XP_037043224.1 | Aldo-keto reductase family 1 | Vitamin C | Diptera |
| XP_037037530.1 | Aldo-keto reductase family 1 | Vitamin C | Diptera |
| XP_037040291.1 | Aldo-keto reductase family 1 | Vitamin C | Diptera |
| XP_037037486.1 | Aldo-keto reductase family 1 | Vitamin C | Diptera |
| XP_037040241.1 | Aldo-keto reductase family 1 | Vitamin C | Diptera |
| XP_037025978.1 | Aldo-keto reductase family 1 | Vitamin C | Diptera |
| XP_037049259.1 | Aldo-keto reductase family 1 | Vitamin C | Diptera |
| XP_037049256.1 | Aldo-keto reductase family 1 | Vitamin C | Diptera |
| XP_037052243.1 | Aldo-keto reductase family 1 | Vitamin C | Diptera |
| XP_037037005.1 | Aldo-keto reductase family 1 | Vitamin C | Diptera |
| XP_080118818.1 | Aldo-keto reductase family 1 | Vitamin C | Diptera |
| XP_080118066.1 | Aldo-keto reductase family 1 | Vitamin C | Diptera |
| XP_080114100.1 | Aldo-keto reductase family 1 | Vitamin C | Diptera |
| XP_037044813.1 | Aldo-keto reductase family 1 | Vitamin C | Diptera |
| XP_037039507.1 | Aldo-keto reductase family 1 | Vitamin C | Diptera |
| XP_037048086.1 | Aldo-keto reductase family 1 | Vitamin C | Diptera |
| XP_037031486.1 | Aldo-keto reductase family 1 | Vitamin C | Diptera |
| XP_080119631.1 | Aldo-keto reductase family 1 | Vitamin C | Diptera |
| XP_037036457.2 | Aldo-keto reductase family 1 | Vitamin C | Diptera |
| XP_037052277.1 | Aldo-keto reductase family 1 | Vitamin C | Diptera |
| XP_037028837.1 | Regucalcin | Vitamin C | Arachnida |
| XP_080114029.1 | Regucalcin | Vitamin C | Collembola |
| XP_037044301.1 | Regucalcin | Vitamin C | Collembola |
| XP_037028785.1 | Regucalcin | Vitamin C | Collembola |
| XP_037033188.1 | Regucalcin | Vitamin C | Diptera |
| XP_037037501.1 | Regucalcin | Vitamin C | Diptera |
| XP_037043734.1 | Regucalcin | Vitamin C | Diptera |
| XP_037044175.1 | Regucalcin | Vitamin C | Diptera |
| XP_037044176.1 | Regucalcin | Vitamin C | Diptera |
| XP_080113076.1 | Regucalcin | Vitamin C | Diptera |
| XP_037028311.1 | Regucalcin | Vitamin C | Diptera |

| | | | |
|---|---|---|---|
| XP_037052276.1 | Regucalcin | Vitamin C | Diptera |
| XP_037046716.1 | Regucalcin | Vitamin C | Diptera |
| XP_080115552.1 | Regucalcin | Vitamin C | Diptera |
| XP_037037356.2 | Regucalcin | Vitamin C | *Diptera |
| XP_037033252.1 | Regucalcin | Vitamin C | Diptera |
| XP_037040989.1 | Regucalcin | Vitamin C | Diptera |
| XP_080117808.1 | L-gulono-1,4-lactone oxidase | Vitamin C | Collembola |
| XP_080112003.1 | L-gulono-1,4-lactone oxidase | Vitamin C | Collembola |
| XP_037049943.1 | L-gulono-1,4-lactone oxidase | Vitamin C | Collembola |
| XP_037029701.1 | L-gulono-1,4-lactone oxidase | Vitamin C | Collembola |
| XP_037051389.1 | L-gulono-1,4-lactone oxidase | Vitamin C | Collembola |
| XP_037042731.1 | L-gulono-1,4-lactone oxidase | Vitamin C | Collembola |
| XP_080119402.1 | L-gulono-1,4-lactone oxidase | Vitamin C | Collembola |
| XP_037034048.2 | L-gulono-1,4-lactone oxidase | Vitamin C | Collembola |
| XP_037034050.2 | L-gulono-1,4-lactone oxidase | Vitamin C | Collembola |
| XP_080114438.1 | L-gulono-1,4-lactone oxidase | Vitamin C | Collembola |
| XP_037038882.1 | L-gulono-1,4-lactone oxidase | Vitamin C | Collembola |
| XP_080117800.1 | L-gulono-1,4-lactone oxidase | Vitamin C | Collembola |
| XP_037049935.1 | L-gulono-1,4-lactone oxidase | Vitamin C | Collembola |
| XP_080117809.1 | L-gulono-1,4-lactone oxidase | Vitamin C | Collembola |
| XP_037025866.1 | L-gulono-1,4-lactone oxidase | Vitamin C | Collembola |
| XP_037045984.1 | L-gulono-1,4-lactone oxidase | Vitamin C | Collembola |
| XP_080114557.1 | L-gulono-1,4-lactone oxidase | Vitamin C | Collembola |
| XP_037049532.1 | Ergothioneine synthesis protein 1 (histidine-specific methyltransferase) | Ergothioneine | Fungi |
| XP_080112150.1 | Ergothioneine synthesis protein 2 (hercynylcysteine sulfoxide lyase) | Ergothioneine | Collembola |
| XP_037026235.1 | Ergothioneine synthesis protein 2 (hercynylcysteine sulfoxide lyase) | Ergothioneine | Collembola |
| XP_037037559.1 | Ergothioneine synthesis protein 2 (hercynylcysteine sulfoxide lyase) | Ergothioneine | Collembola |
| XP_037038869.1 | Pyridoxal 5'-phosphate synthase subunit PdxS/SNZ | Pyridoxal 5’-phosphate | Bacteria |
| XP_037038870.1 | Pyridoxal 5'-phosphate synthase subunit PdxT/SNO | Pyridoxal 5’-phosphate | Fungi |
| XP_037040229.1 | Isopenicillin-N synthase | β-lactam antibiotics | Collembola |
| XP_037037668.1 | N-(5-amino-5-carboxypentanoyl)-L-cysteinyl-D-valine synthase | β-lactam antibiotics | Collembola |

**Table S23. Results of Cox proportional hazards regression for *D. melanogaster* transgenic experiments.** Column descriptors are as follows: "Comparison" indicates the statistical comparison made, with "B" in "A vs. B" indicating the baseline; "Treatment" indicates the experimental or control exposure; "N_reps" indicates the number of independent replicate vials for each genotype or condition, in the same order as the "Comparison" column, with 25 flies per vial; "HR" is the hazard ratio, with values < 1 indicating reduced death risk and values > 1 indicating increased death risk relative to the baseline; "CI" columns indicate the lower and upper bounds of the 95% confidence interval of the hazard ratio; the "p-value" indicates significance relative to the null hypothesis that HR = 1. Gray shaded rows indicate significant protective effects (HR < 1 and p-value < 0.05). N/A for sex columns indicates that Cox regression was not split by sex (but sex and genotypes were included as covariates); N/A for "Treatment" indicates that the comparison was between different treatments.

| **Comparison** | **Treatment** | **N_reps** | **Sex** | **HR** | **CI (low)** | **CI (upper)** | **p-value** |
|---|---|---|---|---|---|---|---|
| 1%suc + 1%$H_2O_2$ vs. 1%suc | N/A | 50 vs. 50 | N/A | 35.71 | 21.29 | 59.92 | $8.59 \times 10^{-42}$ |
| Actin5c>Bc8ODP vs. UAS Bc8ODP/+ | 1%suc + 1%$H_2O_2$ | 5 vs. 5 | Male | 0.78 | 0.51 | 1.19 | $2.48 \times 10^{-1}$ |
| Actin5c>Bc8ODP vs. Actin5c> | 1%suc + 1%$H_2O_2$ | 5 vs. 5 | Male | 0.28 | 0.16 | 0.49 | $1.01 \times 10^{-5}$ |
| Actin5c>Bc8ODP vs. UAS Bc8ODP/+ | 1%suc + 1%$H_2O_2$ | 5 vs. 5 | Female | 0.43 | 0.29 | 0.65 | $6.80 \times 10^{-5}$ |
| Actin5c>Bc8ODP vs. Actin5c> | 1%suc + 1%$H_2O_2$ | 5 vs. 5 | Female | 0.52 | 0.32 | 0.84 | $7.51 \times 10^{-3}$ |
| Actin5c>Bc8ODP vs. UAS Bc8ODP/+ | 1%suc | 5 vs. 5 | Male | 8.06 | 3.86 | 16.83 | $2.72 \times 10^{-8}$ |
| Actin5c>Bc8ODP vs. Actin5c> | 1%suc | 5 vs. 5 | Male | 0.89 | 0.43 | 1.82 | $7.40 \times 10^{-1}$ |
| Actin5c>Bc8ODP vs. UAS Bc8ODP /+ | 1%suc | 5 vs. 5 | Female | 0.72 | 0.48 | 1.08 | $1.14 \times 10^{-1}$ |
| Actin5c>Bc8ODP vs. Actin5c> | 1%suc | 5 vs. 5 | Female | 0.48 | 0.27 | 0.84 | $1.06 \times 10^{-2}$ |
| Actin5c>BcAPEX2 vs. UAS BcAPEX2/+ | 1%suc + 1%$H_2O_2$ | 5 vs. 5 | Male | 0.83 | 0.49 | 1.40 | $4.80 \times 10^{-1}$ |
| Actin5c>BcAPEX2 vs. Actin5c> | 1%suc + 1%$H_2O_2$ | 5 vs. 5 | Male | 0.32 | 0.16 | 0.62 | $6.65 \times 10^{-4}$ |
| Actin5c>BcAPEX2 vs. UAS BcAPEX2/+ | 1%suc + 1%$H_2O_2$ | 5 vs. 5 | Female | 0.66 | 0.42 | 1.04 | $7.53 \times 10^{-2}$ |
| Actin5c>BcAPEX2 vs. Actin5c> | 1%suc + 1%$H_2O_2$ | 5 vs. 5 | Female | 0.63 | 0.39 | 1.02 | $5.87 \times 10^{-2}$ |
| Actin5c>BcAPEX2 vs. UAS BcAPEX2/+ | 1%suc | 5 vs. 5 | Male | 3.91 | 2.77 | 5.51 | $6.47 \times 10^{-15}$ |
| Actin5c>BcAPEX2 vs. Actin5c> | 1%suc | 5 vs. 5 | Male | 0.77 | 0.41 | 1.44 | $4.19 \times 10^{-1}$ |
| Actin5c>BcAPEX2 vs. UAS BcAPEX2/+ | 1%suc | 5 vs. 5 | Female | 0.57 | 0.41 | 0.79 | $8.39 \times 10^{-4}$ |
| Actin5c>BcAPEX2 vs. Actin5c> | 1%suc | 5 vs. 5 | Female | 0.61 | 0.42 | 0.89 | $1.07 \times 10^{-2}$ |
| 1mM Arsenite + 4-24 Diet vs. 4-24 Diet | N/A | 44 vs. 43 | N/A | 34.27 | 15.71 | 74.75 | $6.56 \times 10^{-19}$ |
| Actin5c>BcArsM vs. UAS BcArsM/+ | 4-24 diet | 10 vs. 7 | Female | 3.77 | 1.59 | 8.93 | $2.55 \times 10^{-3}$ |
| Actin5c>BcArsM vs. Actin5c> | 4-24 diet | 10 vs. 7 | Female | 0.62 | 0.19 | 2.07 | $4.41 \times 10^{-1}$ |

| | | | | | | | |
|---|---|---|---|---|---|---|---|
| Actin5c>BcArsM vs. UAS BcArsM /+ | 4-24 diet | 7 vs. 7 | Male | 0.89 | 0.18 | 4.34 | $8.90 \times 10^{-1}$ |
| Actin5c>BcArsM vs. Actin5c> | 4-24 diet | 7 vs. 5 | Male | 0.83 | 0.37 | 1.86 | $6.44 \times 10^{-1}$ |
| Actin5c>BcArsM vs. UAS BcArsM/+ | 1mM arsenite + 4-24 diet | 9 vs. 7 | Female | 0.49 | 0.37 | 0.66 | $2.42 \times 10^{-6}$ |
| Actin5c>BcArsM vs. Actin5c> | 1mM arsenite + 4-24 diet | 9 vs. 8 | Female | 0.40 | 0.30 | 0.54 | $2.36 \times 10^{-9}$ |
| Actin5c>BcArsM vs. UAS BcArsM /+ | 1mM arsenite + 4-24 diet | 8 vs. 7 | Male | 0.20 | 0.14 | 0.28 | $2.73 \times 10^{-22}$ |
| Actin5c>BcArsM vs. Actin5c> | 1mM arsenite + 4-24 diet | 8 vs. 5 | Male | 0.29 | 0.21 | 0.40 | $2.65 \times 10^{-14}$ |

**Table S24. Selected dipteran genes and HTGs with predicted function in repair of damaged DNA bases,** related to Figure 5B. Ancestry for putative HTGs was inferred directly from closest relatives when tree topologies unambiguously indicated a single HGT donor, or else from the top non-sciarid DIAMOND hit. XP_080113008.1 (TDG) was identified as a putative HTG of hymenopteran affinity via DIAMOND but found to group phylogenetically with non-sciarid dipteran sequences; XP_037030920.1 (MGMT) was identified as a putative HTG of bacterial affinity via DIAMOND and nested within bacterial sequences in its gene tree, but was excluded from the final set of HTGs due to a single non-sciarid dipteran sequence in its sister clade. We detected nine putative ADP-ribosylation HTGs, some of which may participate in functions outside of DNA repair[65]. Here, we report the Poly (ADP-ribose) polymerase (PARP) with evidence of a role in DNA repair via significant transcriptional upregulation upon irradiation of larvae (Table S14) and predicted nuclear localization. The APEX2 annotation was assigned from InterProScan domains (Table S18), and PARP1 was identified as the top BLASTp hit of its respective *Drosophila melanogaster* homolog; the remaining dipteran proteins were identified directly from RefSeq annotation descriptions and their associated NCBI Conserved Domain Database annotations. All HTGs in this table have predicted nuclear localization.

| Accession | Annotation | Predicted function | Ancestry |
|---|---|---|---|
| XP_037049870.1 | Oxidized purine nucleoside triphosphate hydrolase (8ODP) | Oxidized base pair degradation | Insecta |
| XP_037051802.1 | DNA oxidative demethylase AlkB homolog 1 (ALKBH1) | Direct base pair damage reversal | Diptera |
| XP_037050925.1 | DNA oxidative demethylase AlkB homolog 1 (ALKBH1) | Direct base pair damage reversal | Diptera |
| XP_080119171.1 | DNA oxidative demethylase AlkB homolog 2 (ALKBH2) | Direct base pair damage reversal | Arachnida |
| XP_080118158.1 | RNA/DNA oxidative demethylase AlkB homolog 6 (ALKBH6) | Direct base pair damage reversal | Diptera |
| XP_037030920.1 | Methylated-DNA--protein-cysteine methyltransferase (MGMT) | Direct base pair damage reversal | Bacteria* |
| XP_037040393.1 | Uracil-DNA glycosylase (UDG) | Monofunctional glycosylase | Arachnida |
| XP_037036837.1 | Methylpurine DNA glycosylase (MPG) | Monofunctional glycosylase | Collembola |
| XP_080113008.1 | Thymine-DNA glycosylase (TDG) | Monofunctional glycosylase | Diptera |
| XP_037042473.1 | Apurinic or apyrimidinic site endonuclease family 2 member (APEX2) | AP site endonuclease | Arachnida |
| XP_037027975.1 | Apurinic or apyrimidinic site endonuclease family 1 member (APEX1) | AP site endonuclease | Collembola |
| XP_037038972.1 | Apurinic or apyrimidinic site endonuclease family I member (APEX1) | AP site endonuclease | Diptera |
| XP_037038372.1 | 8-oxoguanine DNA glycosylase (OGG1) | Bifunctional glycosylase | Diptera |
| XP_037035410.1 | 8-oxoguanine DNA glycosylase (OGG1) | Bifunctional glycosylase | Diptera |
| XP_037043837.1 | Nth-like DNA glycosylase (NTH) | Bifunctional glycosylase | Diptera |
| XP_037028270.1 | DNA polymerase epsilon catalytic subunit 1 (POLE) | DNA polymerization | Diptera |
| XP_037025554.1 | DNA polymerase delta catalytic subunit (POLD) | DNA polymerization | Diptera |
| XP_037047531.1 | Replication factor C subunit 1 (RFC1) | DNA polymerization | Diptera |
| XP_037033130.1 | Flap endonuclease 1 (FEN1) | Flap excision | Diptera |
| XP_037042621.1 | Proliferating cell nuclear antigen (PCNA) | Flap excision | Diptera |
| XP_037038333.1 | DNA repair protein XRCC1 | DNA ligation | Diptera |

| | | | |
|---|---|---|---|
| XP_037029027.1 | DNA ligase 1 (LIG1) | DNA ligation | Diptera |
| XP_037039700.1 | Poly [ADP-ribose] polymerase 1 (PARP1) | DNA ligation, DNA polymerization | Diptera |
| XP_037051671.1 | Poly [ADP-ribose] polymerase (PARP) tankyrase | DNA ligation, DNA polymerization | Collembola |

**Table S25. RefSeq gene and protein accessions for glyoxylate shunt and β-lactam biosynthesis** genes across *B. coprophila*, *F. candida*, and *P. metropolitanus*.

| Enzyme name | Species | Protein accession | Gene accession |
|---|---|---|---|
| N-(5-amino-5-carboxypentanoyl)-L-cysteinyl-D-valine synthase | *B. coprophila* | XP_037037668.1 | LOC119075341 |
| N-(5-amino-5-carboxypentanoyl)-L-cysteinyl-D-valine synthase | *F. candida* | XP_035703636.1 | LOC110843573 |
| N-(5-amino-5-carboxypentanoyl)-L-cysteinyl-D-valine synthase | *P. metropolitanus* | XP_055346233.1 | LOC129593796 |
| Isopenicillin-N synthase | *B. coprophila* | XP_037040229.1 | LOC119077147 |
| Isopenicillin-N synthase | *F. candida* | XP_021945242.1 | LOC110843583 |
| Isopenicillin-N synthase | *P. metropolitanus* | XP_055346169.1 | LOC129593743 |
| Malate synthase | *B. coprophila* | XP_037048618.1 | LOC119083097 |
| Malate synthase | *F. candida* | XP_021962959.1 | LOC110858471 |
| Isocitrate lyase | *B. coprophila* | XP_037048622.1 | LOC119083098 |
| Isocitrate lyase | *F. candida* | XP_021962860.1 | LOC110858398 |

**Table S26. Putative HTGs predicted to function in carbohydrate catabolism,** associated with Figure S9. “Enzyme name” indicates the functional prediction assigned via manual annotation; “Substrate” is the carbohydrate the enzyme is predicted to catabolize; “N_HTGs” indicates the number of tree-confirmed putative HTGs assigned to each enzyme name; “N_Ex/M” indicates the number of HTGs with DeepLoc2-predicted extracellular or cell membrane localization.

| Enzyme name | N_HTGs | N_Ex/M | Substrate |
|---|---|---|---|
| Endoglucanase (GH9) | 8 | 7 | Cellulose |
| Endoglucanase (GH45) | 7 | 7 | Cellulose |
| Endochitinase-like (GH?) | 1 | 1 | Chitin |
| Chitin deacetylase (CE4) | 1 | 1 | Chitin |
| Chitinase (GH18) | 2 | 2 | Chitin |
| Chitinase (GH19) | 6 | 6 | Chitin |
| N-acetyl-β-hexosaminidase (GH20) | 4 | 4 | Chitin |
| Lytic chitin monooxygenase (AA15) | 5 | 5 | Chitin |
| β-mannosidase (GH2) | 2 | 2 | Mannan/hemicellulose |
| Endo-1,4-β-mannosidase (GH5_10) | 1 | 1 | Mannan/hemicellulose |
| β-mannosidase (GH130) | 1 | 1 | Mannan/hemicellulose |
| Pectin/pectate lyase-like (PL?) | 1 | 1 | Pectin |
| Polygalacturonase (GH28) | 5 | 5 | Pectin |
| Pectin methylesterase (CE8) | 2 | 2 | Pectin |
| Pectin/pectate lyase (PL1_7) | 6 | 6 | Pectin |
| N-acetylmuramic acid 6-phosphate etherase MurQ | 1 | 0 | Peptidoglycan |
| Murein tetrapeptide carboxypeptidase | 2 | 1 | Peptidoglycan |
| Bacterial cell wall hydrolase | 3 | 2 | Peptidoglycan |
| Lysozyme (GH24) | 2 | 2 | Peptidoglycan |
| Domesticated amidase effector | 1 | 1 | Peptidoglycan |
| β-D-xylosidase (GH3) | 3 | 2 | Xylan/hemicellulose |
| α-L-arabinofuranosidase (GH51) | 2 | 1 | Xylan/hemicellulose |
| Endo-1,4-β-xylanase (GH11) | 2 | 2 | Xylan/hemicellulose |
| Xylan exo-α-(1-2)-glucuronosidase (GH67) | 1 | 0 | Xylan/hemicellulose |
| Exo-α-(1-5)-L-arabinofuranosidase (GH43_26) | 2 | 2 | Xylan/hemicellulose |
| Exoglucanase/xylanase (GH10) | 1 | 1 | Xylan/hemicellulose |
| Endo-1,4-β-xylanase (GH43_34) | 1 | 1 | Xylan/hemicellulose |
| Glucan-endo-1,3-α-glucosidase (GH71) | 2 | 2 | α-1,3-glucan |
| Endo-β-1,3-glucanase (GH16) | 1 | 1 | β-1,3-glucan |
| Exo-β-1,3-glucanase (GH5_9) | 1 | 1 | β-1,3-glucan |
| Glucan-1,3-β-glucosidase (GH5) | 1 | 1 | β-1,3-glucan |
| β-1,3-glucanase (GH17) | 3 | 3 | β-1,3-glucan |
| Glucan-1,3-β-glucosidase (GH55) | 1 | 1 | β-1,3-glucan |
| L-fucose-proton symporter | 1 | 0 | Fucose |
| Sodium-dependent glucose transporter (SGLT) | 12 | 10 | Glucose |

**Table S27**. **Expression of putative glyoxylate shunt HTGs.** Mean-pooled transcripts per million (TPM) expression values are shown for adult and larval samples, pooled across both sexes. *r* values indicate Pearson correlation coefficients for TPMs across developmental stages with phosphoenolpyruvate carboxykinase 1 (RefSeq gene accession Pepck1) and carnitine palmitoyltransferase 1 (CPT1, RefSeq gene accession whd). **** indicates significant correlations with p-value < $10^{-4}$; ** indicates significant p-values > 0.01 and < 0.05.

| Enzyme name | RefSeq gene accession | Mean adult TPM | Mean larval TPM | *r* with Pepck1 | *r* with CPT1 |
|---|---|---|---|---|---|
| Isocitrate lyase | LOC119083098 | 1,486.35 | 16.81 | 0.91**** | 0.81**** |
| Malate synthase | LOC119083097 | 299.29 | 26.60 | 0.56** | 0.48** |

**Table S28. Genomes used for species tree inference**. In Data type column, “Protein” indicates that inputs to BUSCO were protein FASTA files taken from NCBI annotations; “Nucleotide” indicates that inputs were unannotated genomes; “BRAKER” indicates that BRAKER3 protein predictions were inputs.

| Species | Family | Genome accession | Data type |
|---|---|---|---|
| *Aedes aegypti* | Culicidae | GCF_002204515.2 | Protein |
| *Hermetia illucens* | Stratiomyidae | GCF_905115235.1 | Protein |
| *Drosophila melanogaster* | Drosophilidae | GCF_000001215.4 | Protein |
| *Sitodiplosis mosellana* | Cecidomyiidae | GCF_009176505.1 | Protein |
| *Contarinia nasturtii* | Cecidomyiidae | GCF_009176525.2 | Protein |
| *Mycetophila fungorum* | Mycetophilidae | GCA_981263315.1 | Nucleotide |
| *Diadocidia ferruginosa* | Diadocidiidae | GCA_027564275.1 | Nucleotide |
| *Bradysia odoriphaga* | Sciaridae | GCA_016920775.1 | Protein |
| *Bradysia coprophila* | Sciaridae | GCF_014529535.2 | Protein |
| *Bradysia impatiens* | Sciaridae | GCA_965233325.1 | BRAKER |
| *Lycoriella ingenua* | Sciaridae | GCA_965233655.1 | BRAKER |
| *Pseudolycoriella hygida* | Sciaridae | GCA_029228625.1 | Protein |

**Table S29. Dipteran fossils used for soft minimum and maximum constraints in MCMCtree.** MYA = millions of years ago.

| **Node description** | **Minimum constraint** | **Minimum justification** | **Maximum constraint** | **Maximum justification** |
|---|---|---|---|---|
| *Lycoriella-Bradysia* ancestor | 35.0 MYA | *Lycoriella* fossils have been detected in Baltic amber from 35-50 MYA[205]. | | |
| Sciaridae-Diadocidiidae ancestor | 125.8 MYA | Sciaridae fossils recovered from England from 125.8-132.6 MYA[206]. | | |
| Sciaridae-Mycetophilidae ancestor | 139.0 MYA | *Sciophila defossa*, a possible stem-group impression Mycetophilidae fossil detected from 139.0 – 141.0 MYA[206]. | | |
| Sciaridae-Cecidomyiidae ancestor | 143.1 MYA | *Catotricha mesozoica*, a cecidomyiid fossil recovered from Russia, dated to 149.2-143.1 MYA[206]. | | |
| Neodiptera crown group ancestor | 247.0 MYA | *Protoanisolarva juarezi,* of the extant neodipteran family Anisopodidae, recovered from 247 MYA[207]. | | |
| Dipteran crown group ancestor | 247.0 MYA | *Protoanisolarva juarezi,* of the extant neodipteran family Anisopodidae, recovered from 247 MYA[207]. | 270 MYA | Taken from consensus inter-order phylogenies[161,165]. |
| Brachycera crown group ancestor | 195.0 MYA | *Oligophryne*, a crown group brachyceran fossil, recovered from 195 MYA[165]. | | |

**Table S30. Parameters used for MCMCtree inference.**

| Parameter | Value |
|---|---|
| clock | 3 |
| aaRateFile | insect.dat[173] |
| alpha | 0.5 |
| ncatG | 9 |
| cleandata | 0 |
| BDparas | 1 1 0 |
| rgene_gamma | 2 7.8299 |
| sigma2_gamma | 1 10 |
| burnin | 100000 |
| sampfreq | 100 |
| nsample | 300000 |
